# Small Molecule *in situ* Resin Capture – A Compound First Approach to Natural Product Discovery

**DOI:** 10.1101/2023.03.02.530684

**Authors:** Alexander Bogdanov, Mariam N. Salib, Alexander B. Chase, Heinz Hammerlindl, Mitchell N. Muskat, Stephanie Luedtke, Elany Barbosa da Silva, Anthony J. O’Donoghue, Lani F. Wu, Steven J. Altschuler, Tadeusz F. Molinski, Paul R. Jensen

## Abstract

Microbial natural products remain an important resource for drug discovery. Yet, commonly employed discovery techniques are plagued by the rediscovery of known compounds, the relatively few microbes that can be cultured, and laboratory growth conditions that do not elicit biosynthetic gene expression among myriad other challenges. Here we introduce a culture independent approach to natural product discovery that we call the Small Molecule In situ Resin Capture (SMIRC) technique. SMIRC exploits in situ environmental conditions to elicit compound production and represents a new approach to access poorly explored chemical space by capturing natural products directly from the environments in which they are produced. In contrast to traditional methods, this compound-first approach can capture structurally complex small molecules across all domains of life in a single deployment while relying on Nature to provide the complex and poorly understood environmental cues needed to elicit biosynthetic gene expression. We illustrate the effectiveness of SMIRC in marine habitats with the discovery of numerous new compounds and demonstrate that sufficient compound yields can be obtained for NMR-based structure assignment. Two new compound classes are reported including one novel carbon skeleton that possesses a functional group not previously observed among natural products and a second that possesses potent biological activity. We introduce expanded deployments, in situ cultivation, and metagenomics as methods to facilitate compound discovery, enhance yields, and link compounds to producing organisms. This compound first approach can provide unprecedented access to new natural product chemotypes with broad implications for drug discovery.

**Significance Statement:** Pharmaceutically relevant microbial natural products have traditionally been discovered using a ‘microbe-first’ approach in which bioassays are used to guide the isolation of active compounds from crude culture extracts. While once productive, it is now widely recognized that this approach fails to access the vast chemical space predicted from microbial genomes. Here, we report a new approach to natural product discovery in which compounds are captured directly from the environments in which they are produced. We demonstrate the applications of this technique with the isolation and identification of both known and new compounds including several that possess new carbon skeletons and one with promising biological activity.

## Introduction

Microbial natural products account for many of today’s essential medicines including most antibiotics (1, 2). These compounds are traditionally discovered using a ‘microbe-first’ approach, where individual strains are isolated from Nature, cultured in the laboratory, and screened using bioassays to guide the isolation of active compounds from culture extracts. While once productive, the limitations to this approach are now well documented (3) and include the re-isolation of known compounds (4), the recognition that only a small percentage of bacterial diversity has been cultured (5), and observations that most biosynthetic gene clusters (BGCs) are silent under laboratory growth conditions (6). Efforts to address these limitations include improved cultivation methods (7), co-cultivation or the addition of elicitors to activate silent BGCs (8, 9), and genome mining (10, 11), where BGCs are activated via genetic manipulation (12) or heterologous expression (13). While genome mining has shown promise (14), it is time consuming, limited to the small percentage of microbial diversity that has been cultured (15), and targets one BGC at a time with no a priori evidence that the products, once isolated and characterized, will be chemically novel or have medicinal relevance (16). Recent evidence of unique biosynthetic potential in yet-to-be cultured bacteria (17), as well as meta-analyses of genomic (18) and metagenomic (19) data all suggest that our best discovery efforts have failed to realize the natural product biosynthetic potential encoded in microbial genomes(20).

Here we report a new approach for microbial natural product discovery that bypasses the initial need for laboratory cultivation. Rather than relying on cultured strains to drive discoveries, natural products are captured directly from the environment in which they are produced. This culture independent approach, which we call the Small Molecule In Situ Resin Capture (SMIRC) technique, inverts traditional discovery paradigms by targeting compounds without initial consideration for the producing organism. SMIRC builds upon targeted, solid phase methods developed for toxin monitoring (21) and was first employed by us to detect natural products and their producers in marine sediments (22). The technique is agnostic to the biological source of the compounds and thus can capture small molecules originating from bacteria to phytoplankton and, as we show, some that were previously reported from marine plants and invertebrates. As such, it requires no knowledge of cultivation conditions or the factors required to induce biosynthetic gene expression, both major obstacles to current discovery strategies. Instead, our approach relies on naturally occurring chemical cues and environmental conditions to trigger natural product biosynthesis. It is designed to capture secreted compounds across a wide range of polarities while avoiding the complex mixtures obtained from direct sample extraction. Here we demonstrate the discovery potential of SMIRC with the isolation and characterization of natural products with two new carbon skeletons, one of which possesses promising biological activity, and the detection of many more potentially new compounds all from a single deployment site.

## Results

### SMIRC deployments and compound dereplication

The Small Molecule In Situ Resin Capture (SMIRC) technique was originally developed to detect microbial natural products in marine sediment (22). Given that most compounds captured at that time could not be identified, we revised the method to determine if it could be used as a tool for natural product discovery. These changes included increasing the deployment times from 24 h to multiple days, increasing the amount of HP-20 resin used to 100 g per replicate, and switching to Nitex mesh enclosures and other modifications to reduce chemical contaminants (Methods). With these modifications, we first deployed SMIRC resins in a well-protected *Zostera marina* seagrass meadow (<2 m) in Mission Bay, San Diego (Fig. 1A) and obtained salt-free, dark brown extracts averaging 54.2 ± 6.1 mg per replicate (n=4). A microfractionation antibiotic assay targeting the outer membrane deficient *E. coli* strain lptD4123 (23) revealed a large active peak with a UV maximum at 360 nm (Fig. 1B). Major compounds in the active fraction (*m/z* 381.0906 and 301.1396) were detected using high-resolution mass spectrometry (MS) with the mass difference of 79.951 Da attributed to the loss of SO_3_ (calcd. 79.9568 Da). Bioassay-guided C_18_ fractionation from one SMIRC replicate extract followed by HPLC purification yielded 2 mg of the major compound (*m/z* 301.1396), which was identified as the flavonoid chrysoeriol based on the comparison of MS and NMR data with published reports (Fig. 1C). Sulfated flavonoids with antibacterial activity have been reported from *Z. marina* (24), providing a candidate producer for these compounds. While our goal was to capture novel microbial products, this finding demonstrates the potential applications of SMIRC to address questions in chemical ecology, in this case the role of antibiotic metabolites released by seagrasses in structuring seagrass meadow microbiomes. We estimated that the concentration of chrysoeriol and related flavonoids at the deployment site was <1 µg/L by direct extraction of ambient seawater and quantitative HPLC analysis. Given that SMIRC yielded 20 µg chrysoeriol/mg HP-20 resin, our method concentrated this compound at levels that would be difficult to obtain using direct seawater extraction. This first deployment demonstrated that sufficient compound yields could be obtained using SMIRC for NMR-based structure elucidation.

**Figure 1.**
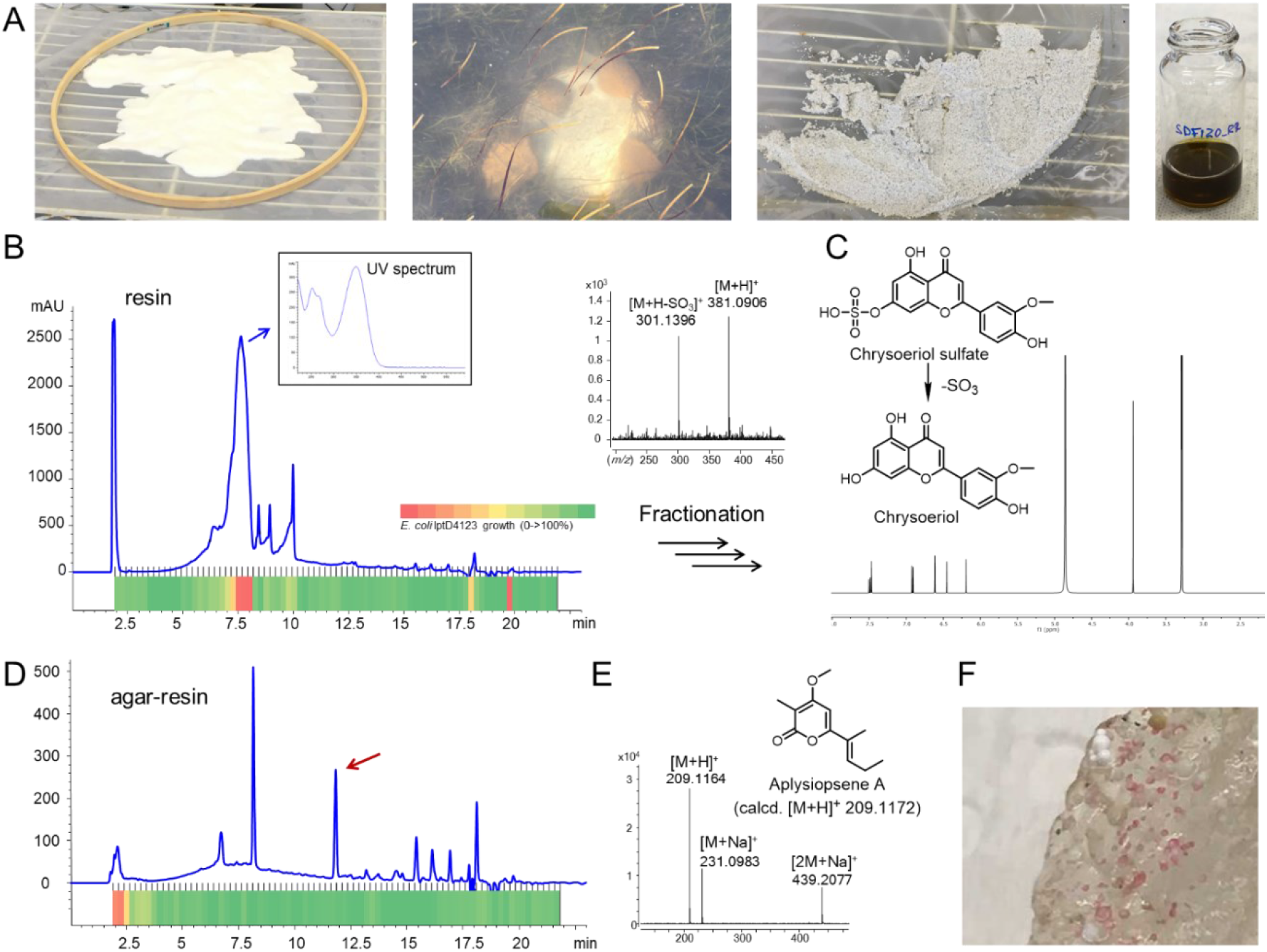
SMIRC deployments in *Zostera marina* meadow (Mission Bay, San Diego). A. From left to right: activated HP-20 resin before enclosing in Nitex, SMIRC deployment, recovered resin before extraction, concentrated extract in MeOH. B. UV_360_ chromatogram of the resin extract and corresponding microfractionation bioassay (80 fractions collected over 20 minutes) with heatmap showing *E. coli* LptD4123 growth inhibition (red = no growth). Blue arrow: UV spectrum of active peak (7.6 min). MS spectrum of the active compound shows loss of a sulfate moiety. C. ^1^H NMR (500 MHz) and structure of the isolated flavonoid chrysoeriol, a degradation product of chrysoeriol sulfate. D. UV_360_ chromatogram of extract generated from resin embedded in agar (in situ cultivation) deployed at the same site revealed an additional peak (red arrow). E. Structure and MS spectrum of aplysiopsene A isolated from the agar/resin peak (red arrow). F. Pink colonies growing on agar/resin matrix that yielded aplysiopsene A.

In a follow-up deployment at the same site, we further modified the SMIRC technique by embedding the resin in an agar matrix with the aim of increasing the yields of microbial-derived compounds by facilitating in situ microbial growth. The addition of agar led to a different antibiotic profile for the extracts and the detection of a major new peak in the UV chromatogram (Fig. 1D). We isolated and identified this compound (0.4 mg) as α-pyrone aplysiopsene A (Fig. 1E) using NMR and tandem MS/MS (SI Appendix). Aplysiopsene A was originally reported from the Atlantic sacoglossan mollusk *Aplysiopsis formosa* (25), while other structurally related compounds have been reported from a sea fan-derived fungus (26) and the actinobacterium *Nocardiopsis dassonvillei* (27). Given that sacoglossans were not observed at the deployment site, the isolation of aplysiopsene A suggests that this compound represents another example of a natural product originally reported from a marine invertebrate that appears to be of microbial origin (28, 29). Of note, we observed pink bacterial colonies (Fig. 1F) on the agar-resin matrix and are working on linking this microorganism to aplysiopsene A production. These results provide a second example in which SMIRC deployments yielded sufficient compound for NMR-based structural elucidation and evidence that embedding the resin in agar can enhance compound discovery, facilitate in situ microbial growth, and provide opportunities to identify the microbial origins of captured natural products. In addition, we identified a variety of known natural products using cheminformatic tools associated with the GNPS platform (30) (SI Appendix, Fig. S1) including lipids, carotenoids, flavonoids, bile acids, and the microbial natural product apratoxin A (SI Appendix, Fig. S2), all of which were validated using reference MS/MS spectra. The detection of apratoxin A expands the geographic distribution of this potent cyanobacterial toxin to temperate waters (31, 32) and demonstrates that untargeted MS analyses can provide important insights into the distribution of diverse natural products captured using the SMIRC technique.

We next deployed SMIRC in the littoral zone at Cabrillo State Marine Reserve (CSMR, Fig. 2A), a relatively pristine habitat comprised of sand patches and rocky reef dominated by the seagrass *Phyllospadix* sp. and inhabited by a diverse assemblage of macroalgae and invertebrates. Deployments (without agar) spanned 2-8 days depending on tides and swells, and yielded extract masses averaging 177.1 mg (±87.6 mg, n=17) with some exceeding 300 mg per 100 g replicate. While no antibiotic activity was detected in the initial screen, LCMS analyses revealed extensive chemical complexity, visualized as congested peaks in the UV (254 nm) and total ion count (TIC) chromatograms (SI Appendix, Fig. S3). Pre-fractionation of the crude extracts greatly facilitated compound detection, increasing the number of detectable molecular features from 732 to 1,231 (68% increase) in crude extracts and fractions, respectively (SI Appendix, Fig. S1). As in the Mission Bay deployments, we detected trace amounts of aplysiopsene A in several of the CSMR extracts, providing further support that this compound has a microbial origin. We also confidently identified okadaic acid, the polyether toxin responsible for diarrhetic shellfish poisoning (33), in several Cabrillo extracts (SI Appendix, Fig. S4). However, numerous compounds detected using the DEREPLICATOR+ algorithm (34) including pectenotoxin-2 (35), callipeltin B, cyanopeptolin 880, amphidinolides, kahalalide Y, and kabiramide D, could not be validated highlighting the need for stringent manual curation of environmental metabolomes (20).

**Figure 2.**
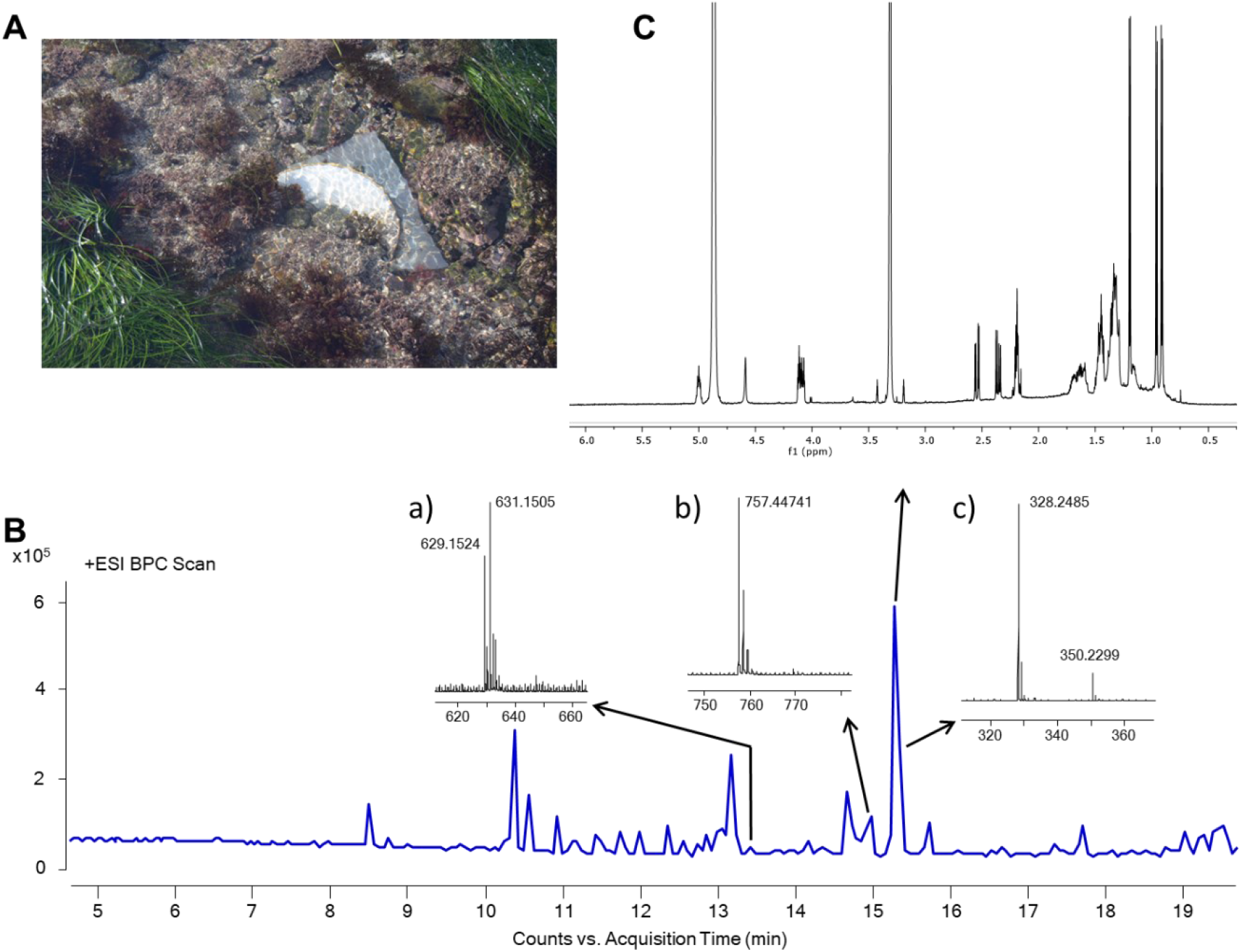
SMIRC deployment at Cabrillo State Marine Reserve (CSMR). A. SMIRC resin deployed on rocky substrate. B. Base peak chromatogram of a SMIRC extract and mass spectra (a-c) of compounds targeted for isolation. C. ^1^H NMR spectrum of the *m/z* 328.2485 compound cabrillostatin (**1**, 40 µg, 600 MHz, 1.7 mm cryoprobe, CD_3_OD).

### Natural product discovery

The large number of molecules that could not be confidently identified in the SMIRC extracts suggests a high degree of chemical novelty and opportunities for natural product discovery. As such, we targeted for isolation a compound with a high-intensity molecular peak (*m/z* 328.2485 [M+H]^+^ Fig. 2B) and calculated molecular formula of C_18_H_33_NO_4_ (calcd [M+H]^+^ 328.2483, 0.61 ppm) with three degrees of unsaturation. This molecular formula matched three compounds in the Dictionary of Natural Products (https://dnp.chemnetbase.com): 3-hydroxy-C_14_-homoserine lactone and curvularides B and E, initially reported from various bacteria and an endophytic fungus (36), respectively. We subjected two extracts with the highest estimated abundance of the compound to LCMS guided isolation (SI Appendix, Methods) augmented by tandem UV and evaporative light-scattering (ELSD) detection given the absence of a strong chromophore. Repeated rounds of HPLC purification yielded ∼40 mg of a pure compound as a colorless solid. The ^1^H NMR spectrum (600 MHz, CD_3_OD, 1.7 mm micro cryoprobe) revealed three doublet resonances attributable to three methyl groups (δ_H_ 0.90, *J* = 6.5 Hz; 0.96, *J* = 6.7; 1.20, *J* = 6.3 Hz) positioned on methine carbons (Fig. 2C), thus indicating that our targeted compound is neither 3-hydroxy-C_14_-homoserine lactone, which possess only one methyl group, nor one of the curvularides, which have five methyl groups. The complete structure elucidation of this compound (SI Appendix, Supplementary Text, Table S1, Figs. S6-S8), which we have called cabrillostatin (**1**, Fig. 3), was accomplished with 2D NMR experiments (^1^H-^1^H COSY, HSQC, HMBC). Cabrillostatin represents a new carbon skeleton comprised of the uncommon non-proteinogenic γ-amino acid statine fused with 9-hydroxydecanoic acid to form a 15-membered macrocycle containing both ester and amide groups. The characterization of this compound by NMR validates the application of SMIRC for natural product discovery.

**Figure 3.**
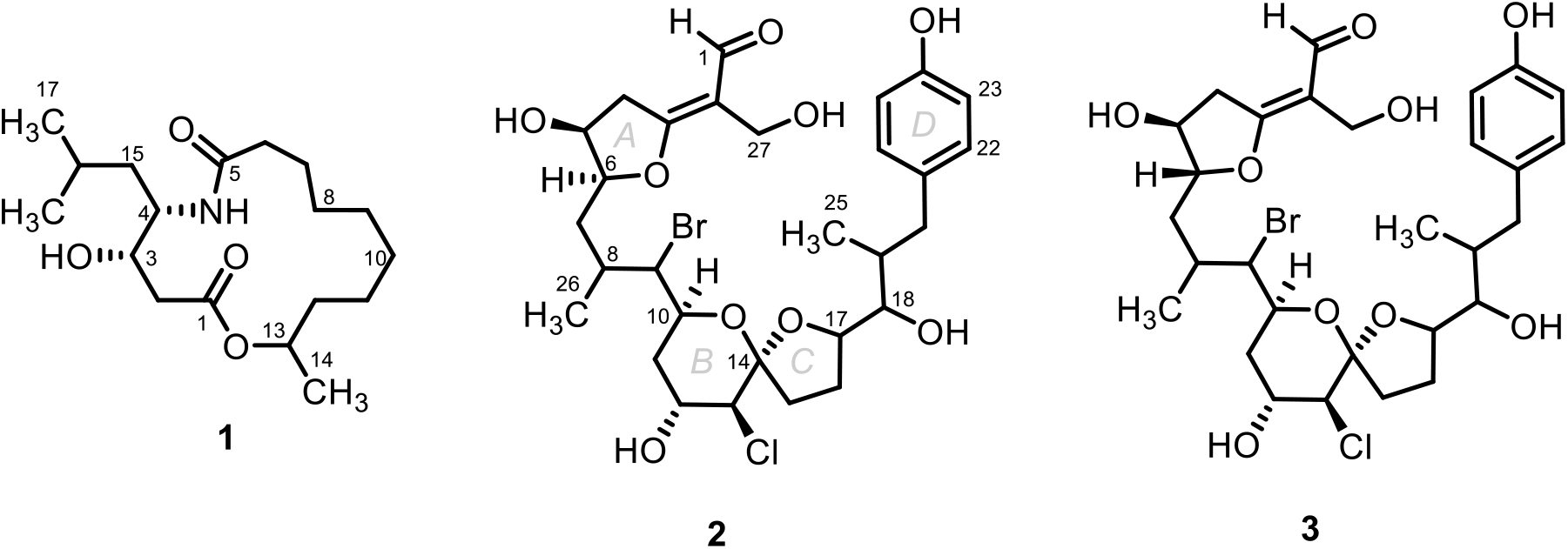
Structures of cabrillostatin (**1**) and cabrillospirals A (**2**) and B (**3**).

While isolating cabrillostatin (**1**), we detected a second compound of interest that displayed an isotopic pattern in the MS spectrum indicative of a dihalogenated molecule bearing one Cl and one Br (Fig. 2B). The best-fit molecular formula C_29_H_40_BrClO_9_, with nine degrees of unsaturation, was calculated from the highest intensity isotopologue (SI Appendix, Fig. S9) at *m/z* 629.1510 (calcd for *m/z* [M-H_2_O+H]^+^ 629.1512, -0.32 ppm). This formula returned no matches in database searches suggesting it represented a second new natural product. Multiple rounds of HPLC purification (SI Appendix, Methods) yielded 50 µg of pure compound from the extract of a 100 mg SMIRC resin deployment. A series of NMR experiments (^1^H, ^1^H-^1^H COSY, HSQC, HMBC, NOESY, 600 MHz, 1.7 mm microcryoprobe, CD_3_CN) conducted over long acquisition times (>72 h for HMBC) left several important correlations missing in the HMBC spectrum. Pooling samples from two SMIRC extracts yielded ∼100 µg of compound while NMR experiments (^1^H, COSY and HSQC) repeated in CD_3_OD improved peak shapes and resolved signals and *S/N*. An LR-HSQMBC (optimized for *J* = 8 Hz) experiment revealed additional correlations not observed in CD_3_CN (37). Extensive analysis of the NMR data recorded in both solvents and MS/MS fragmentation (SI Appendix, Tables S2-S3, Figs. S10-S11) allowed us to establish the structure as a new bromo-chloro-polyketide that we have named cabrillospiral A (**2**) (Fig. 3).

Cabrillospiral A (**2**) exhibits rare features including an α-halogenated bicyclic [6,5]-spiroketal and an α-β unsaturated aldehyde appended to a C=C double bond that is exocyclicly fused to a tetrahydrofuran. The latter, a vinylogous formate ester, is to the best of our knowledge unprecedented among both the natural product and synthetic literature. Cabrillospiral A (**2**) has 11 chiral centers distributed across four independent stereogenic substructures making stereochemical elucidation a challenge. Interpretation of the NOESY data and ^1^H-homonuclear scalar coupling (^2,3^*J*) led to the assignment of the Δ^2^ double bond geometry and the relative stereochemistry of six asymmetric centers, albeit without stereocluster interconnectivity (SI Appendix, Figs. S12-S14). The full assignment of the relative and absolute configurations is ongoing. Cabrillospiral A (**2**) eluted with two minor compounds (SI Appendix, Fig. S11B) that shared the same chromophore and were indistinguishable from **2** by MS/MS. After repeated rounds of HPLC purification from several extracts, we obtained one analog in sufficient amount (27 µg) to measure ^1^H NMR, HSQC, and COSY data (SI Appendix, Table S4) and elucidated the structure as cabrillospiral B (**3**), which we propose as the C-6 epimer of **2**. In addition to compounds **1**-**3**, several other potentially new natural products were isolated from the CSMR deployments (Table 1, SI Appendix, Table S5 and Figs. S15-S25). In total, compounds belonging to two new compound classes (**1** and **2**-**3**) and at least 11 potentially new compounds were recovered from a single deployment site. Importantly, compounds **1, 2**, and unidentified compound *m/z* 757.4474 were consistently detected in three separate deployments spanning seven months, providing a path forward to obtain additional quantities for some lead compounds. Evidence that the greatest yields were obtained from the longest deployments (Fig. 4) justifies more detailed studies of the time course of compound recovery.

**Table 1.** Additional compounds isolated from CSMR deployment site.

| *m/z* | Mass speciation | Molecular formula | Purity | Related natural products: isobaric or isomeric |
| --- | --- | --- | --- | --- |
| 757.4474 | [M+H]<sup>+</sup> | C<sub>43</sub>H<sub>64</sub>O<sub>11</sub> | semi pure | gambierol, amphidinolide C2, glycosylated steroids |
| 456.1317 | [M+Na]<sup>+</sup> | C<sub>20</sub>H<sub>29</sub>Cl<sub>2</sub>NO<sub>5</sub> | 4 isomers in a HPLC fraction | none |
| 1211.6231 | [M+Na]<sup>+</sup> | C<sub>63</sub>H<sub>96</sub>O<sub>21</sub> | HPLC fraction | altohyrtin C, spongistatin 2, steroids |
| 1086.6190 | [M+NH<sub>4</sub>]<sup>+</sup> | C<sub>52</sub>H<sub>80</sub>N<sub>10</sub>O<sub>14</sub> | semi pure | none |
| 836.4025 | [M+H]<sup>+</sup> | C<sub>42</sub>H<sub>72</sub>BrCl<sub>2</sub>NO<sub>6</sub> | HPLC fraction | none |
| 237.1492 | [M+H]<sup>+</sup> | C<sub>14</sub>H<sub>20</sub>O<sub>3</sub> | HPLC fraction | >30 |
| 488.257 | [M+H]<sup>+</sup> | C<sub>23</sub>H<sub>34</sub>ClNO<sub>8</sub> | HPLC fraction | none |
| 606.3572 | [M+H]<sup>+</sup> | C<sub>34</sub>H<sub>52</sub>ClNO<sub>6</sub> | HPLC fraction | none |
| 644.3936 | [M+H]<sup>+</sup> | C<sub>34</sub>H<sub>58</sub>ClNO<sub>8</sub> | semi pure | none |
| 730.2286 | [M+Na]<sup>+</sup> | C<sub>33</sub>H<sub>48</sub>Cl<sub>3</sub>NO<sub>9</sub> | semi pure | none |
| 913.4561 | [M+Na]<sup>+</sup> | C<sub>47</sub>H<sub>70</sub>O<sub>16</sub> | semi pure | glycosylated steroids |

**Figure 4.**
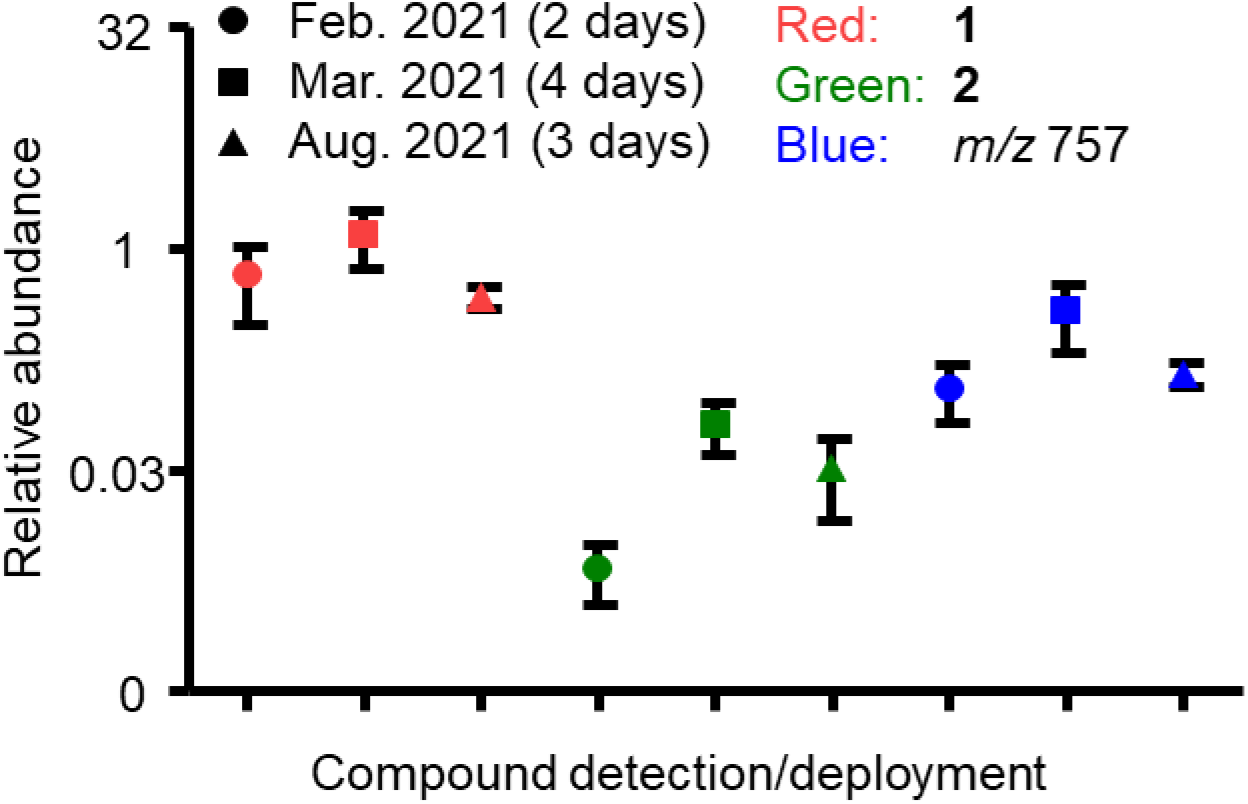
Reproducibility of three target compounds at the CSMR site. In all cases, compound yields were highest for the longest deployments. Cabrillostatin (**1**), cabrillospiral A (**2**), unknown compound (*m/z* 757).

### Bioactivity testing

Multiple SMIRC deployments at the CSMR site allowed us to isolate additional amounts of cabrillostatin (**1**, 26 µg) and cabrillospirals A (**2**, 75 µg) and B (**3**, 27 µg) for biological testing. Given that the amino acid statine is the pharmacophore for some protease inhibitors (38), we tested cabrillostatin (**1**) for activity against the aspartic acid protease cathepsin D. Surprisingly, **1** was not inhibitory but instead potentiated protease activity in a concentration dependent manner from 1.5-fold at 100 nM to almost 2-fold at 10 µM (SI Appendix, Fig. S26). Testing of **1** in an expanded panel of aspartic acid, serine, and cysteine proteases revealed no additional inhibitory activity. We further tested compounds (**1**-**3**) against a panel of nine diverse cancer cell lines (SI Appendix, Table S6) using cell painting (39) and high-content imaging. Phenotypic profiles generated using a customized image analysis pipeline provided in-depth quantification of cell morphology and biomarker staining in comparison with 28 reference compounds across six diverse drug categories with distinguishable phenotypes (40) (SI Appendix, Fig. S27A, Table S7). Principal component analysis (PCA) clearly separated **1** from the DMSO controls in most cell lines (Fig. 5A). Quantifying the separation showed that **1** was significantly bioactive (p<10^−6^) at 10 µM in six of the nine cell lines (Fig. 5B, SI Appendix Table S8). The phenotypic profiles of **1** relative to the reference compounds were unique (confidence score >0.1 (40), SI Appendix, Table S9) indicating that it possess a distinct mechanism of action. A dose-response experiment in A549 cells validated the bioactivity of **1** at 10 µM but failed to detect activity at lower concentrations (SI Appendix, Fig. S27B). In contrast to **1**, cabrillospirals A and B (**2**-**3**) induced no significant phenotypic changes in any of the cell lines.

**Figure 5.**
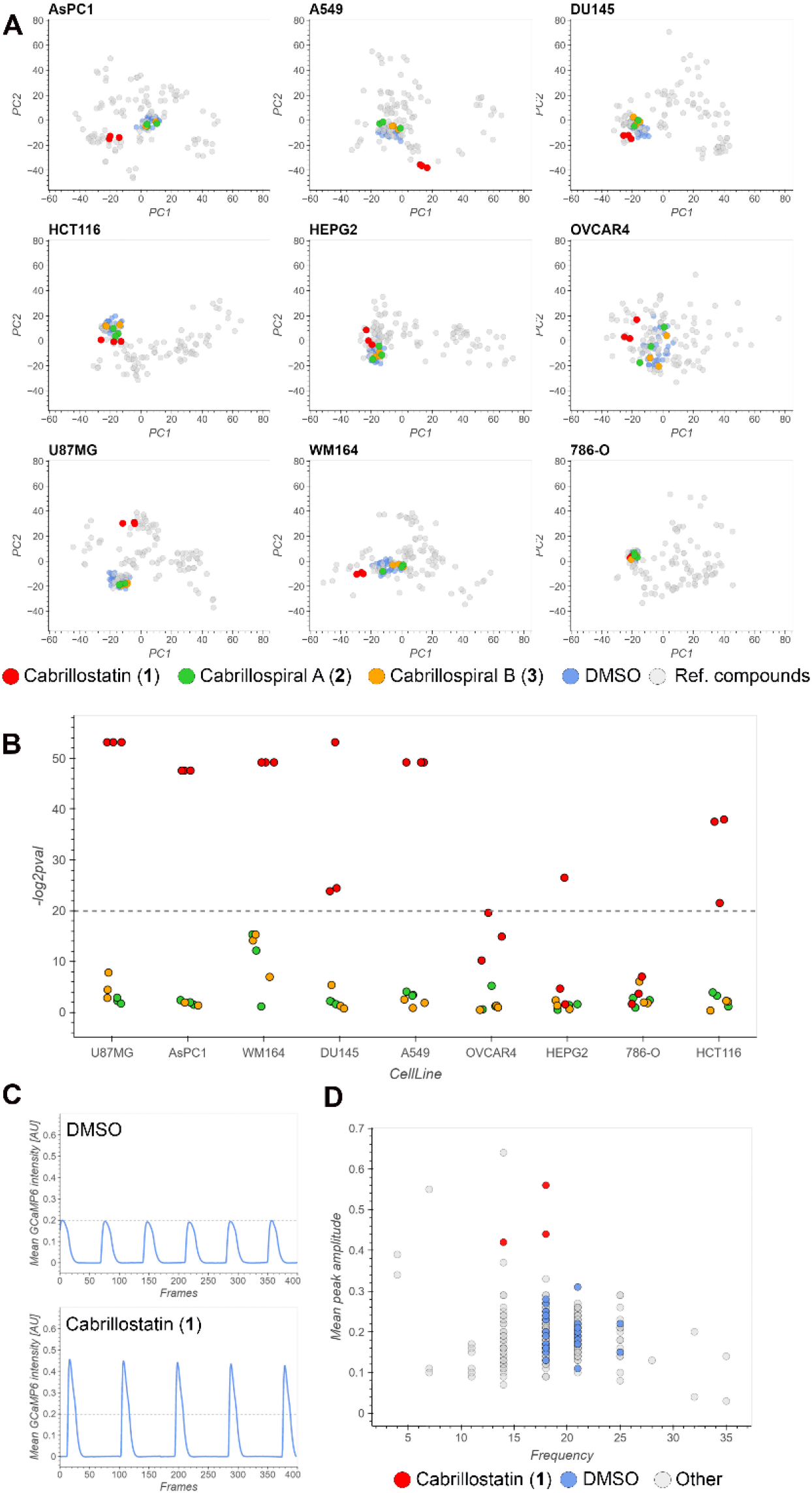
Bioactivity of cabrillostatin (**1**) and cabrillospirals A (**2**) and B (**3**). A. Principal component analysis of high dimensional phenotypic profiles of **1**-**3** (red, green, yellow, respectively), reference compounds (grey), and DMSO (blue) in nine diverse cell lines. B. Bioactivity expressed as significance (-log2pval) across all cell lines. C. Representative Ca^2+^ transient traces of DMSO (top) and **1** (bottom). D. Scatter plot showing beating frequency and mean peak amplitude of **1** (red), DMSO (blue) and anti-arrhythmic and anti-cancer drugs (grey).

Compounds **1-3** were further tested in an induced pluripotent stem cell derived cardiomyocyte (IPSC-CM) model (41) that allows the quantification of Ca^2+^ transients and as such can detect changes in the frequency and amplitude of cardiomyocyte beating induced by anti-arrhythmic or anti-cancer drugs (SI Appendix, Figs. S28A and S28B, Table S10). Concordant with the phenotypic profiling results, compound **1**, but not **2** or **3**, showed a clear alteration of IPSC-CM contractions (Fig. 5C, SI Appendix, Fig. S28C) in the form of increased peak amplitude and a small decrease of peak frequency (Fig. 5D), both of which were validated using an independent batch of IPSC-derived cardiomyocytes (SI Appendix, Fig. S28D). The observed changes in the IPSC-CM Ca^2+^ transients warrant further investigation given the suggested positive inotropic effect. The bioactivity results for cabrillostatin support the application of SMIRC for natural product drug discovery.

### Environmental distributions of compounds ***1*** *and* ***2***

The GNPS platform was recently expanded to include the web-enabled search engine MASST (Mass Spectrometry Search Tool) (42), the equivalent of NCBI BLAST for MS/MS datasets in the MassIVE repository. Using MASST, we parsed ∼2,700 untargeted, small-molecule datasets with the aim of identifying the environmental distributions and potential biotic origins of cabrillostatin (**1**), cabrillospirals A-B (**2**-**3**), aplysiopsene A, and 11 other yet to be characterized compounds (SI Appendix Tables S5 and S11) identified in the SMIRC extracts. Surprisingly, **1** was confidently identified in 11 datasets collected from 2019-2021 and largely associated with the study of dissolved organic matter (DOM) in nearshore (Imperial Beach, Scripps Pier CA) and deep ocean (California current) seawater samples. A match from the metabolomic analysis of a dinoflagellate culture led us to suspect *Ostreopsis* sp. as the biogenic source, however **1** was also detected in the medium controls, which were prepared using seawater collected from the Scripps pier (43) thus suggesting an alternative producer. Cabrillostatin clustered with several related compounds in one public dataset (44), allowing us to assign putative structures for four derivatives based on comparative MS/MS spectral analyses (SI Appendix, Fig. S29). Among these, cabrillostatin B is cyclized with 11-hydroxydodecanoic acid and cabrillostatin B1 bears an additional ketone functionality. In cabrillostatins C and D, the statine moiety is replaced with 4-amino-3-hydroxydeca-4-en-7,9-diynoic acid and with 4-amino-3-hydroxy-5-phenylpentanoic acid, respectively (SI Appendix, Figs. S30-S33). The proposed diynoic amino acid is to the best of our knowledge unprecedented among natural products. Determining the structure of **1** has improved our broader understanding of DOM, which has largely defied characterization (44, 45), and raises questions about the ecological roles of a widespread and biologically active natural product in seawater ecosystems. In contrast to cabrillostatin, the cabrillospirals were not detected in any of the MassIVE datasets, suggesting that this compound is highly site-specific thus demonstrating the potential value of deploying SMIRC across diverse habitats.

### Metagenomic analyses

In an attempt to link compounds identified in the CSMR extracts to the producing organisms, we generated untargeted shotgun metagenomes from marine sediment and bulk water samples (average 76.4±12M paired end reads per sample) collected at the deployment sites. Both the sediment and seawater communities were dominated by Proteobacteria and Bacteroidetes, with higher relative abundances of Actinobacteria and Planctomycetes in the sediment samples (SI Appendix, Fig S34A). In total, we recovered 1,843 BGCs (N50=18 kbp) with many (52%) of the larger BGCs (>10 genes) predicted to be complete (i.e., not on a contig edge). The predicted products of these BGCs are largely nonribosomal peptides (28%), terpenes (25.7%), and ribosomally synthesized and post-translationally modified peptides (RiPPs, 17.7%). They could be clustered into 631 gene cluster families (GCFs, SI Appendix, Fig. S34B), of which only four (those encoding the alkaloid osmolyte ectoine, the peptide natural product anabaenopeptin, and the pigments flexirubin and carotenoids like zeaxanthin) shared similarity to BGCs in the MIBiG reference database (46), emphasizing both the biosynthetic complexity and potential novelty of these environments.

Based on the structure of cabrillostatin (**1**), we used hybrid polyketide synthase–non-ribosomal peptide synthetase (PKS-NRPS) genes as hooks to identify the cognate BGC in the metagenomes. More specifically, we expected a leucine starter unit for the NRPS, a hybrid KS domain for C_2_ elongation of the amino acid to yield the statine moiety, followed by highly reducing PKS modules to account for the saturated fatty acyl component. However, among the 43 PKS-NRPS hybrid BGCs recovered from both sample types, none encoded the predicted enzymes. Interestingly, highly-reducing PKS domains are commonly associated with fungi (47) and the amino acid statine is primarily observed from bacteria (48), providing further ambiguity into the biological origin of **1**. Given the possibility that a low abundant organism (thus, not assembled) could account for production, we extracted and characterized with NaPDoS2 (49) all KS domains from the unassembled reads along with reference sequences from highly-reducing fungal PKSs (50-52) and the statine-related KSs from didemnin B and burkholdac A biosynthesis (29, 53). Unfortunately, none of the metagenome extracted KSs clustered with these reference sequences in a KS phylogeny, leaving the biosynthetic source of **1** unresolved and demonstrating the challenges associated with linking compounds to producers in complex microbial communities.

For cabrillospirals (**2**-**3**), potential biosynthetic ‘hooks’ included a 10+ module type I PKS (T1PKS), halogenases, a *p*-hydroxybenzoic acid (*p*-HBA) starter unit, and methyltransferases to account for the C-26 methyl and C-27 hydroxymethyl groups. Among the 68 T1PKS BGCs identified, our ability to confidently identify a candidate BGC was limited by the assembly challenges associated with highly repetitive modular PKSs (54). Nonetheless, we detected two candidate BGCs (SI Appendix, Fig. S34C) in a high-quality MAG (4.96 Mbp; 98.2% complete, 3.6% contamination) in the phylum Planctomycetes, which was recently recognized as a rich and poorly explored source of specialized metabolites (55). In an attempt to resolve the T1PKS modules in these BGCs, we identified >10 highly similar KS domains (96-98% amino acid similarity) in the unassembled data as would be expected for the cabrillospiral BGC. While inconclusive, the planctomycete MAG remains our top candidate for the production of compound **2**. Unfortunately, long-read Oxford Nanopore sequencing failed to cover this BGC, again likely due to the complexity of the sediment microbial communities.

## Discussion

Explorations of biotic diversity have proven key to the discovery of biologically active natural products including many of today’s most beneficial medicines. The discovery process has invariably started with the collection or cultivation of a specific organism followed by extraction and either chemical or bioassay-guided compound isolation. While effective for decades, it has become increasingly difficult to find structurally unique compounds that can be incorporated into screening programs using these traditional approaches. The realization that microbial genome sequences contain significant, unrealized biosynthetic potential launched the field of genome mining and efforts to activate silent biosynthetic gene clusters through co-cultivation, elicitor addition, or genetic manipulation (13). Despite these advances, current approaches to natural product discovery have failed to capture the vast extent of chemical diversity predicted to exist in Nature.

Here we report on the Small Molecule In Situ Resin Capture (SMIRC) technique, a compound-first approach to natural product discovery in which adsorbent resins are used to capture compounds directly from the environments in which they are produced. Using this technique, we captured both known and unknown natural products and purified sufficient quantities to elucidate the structures of cabrillostatin (**1**) and the halogenated polyketides cabrillospirals A and B (**2**-**3**), which represent two new compound classes. The latter possess a new carbon skeleton and a functional group (vinylogous formate ester) that is unprecedented in both the natural product and synthetic literature. The Tanimoto similarity coefficient calculated for cabrillospiral A (0.47), which describes the similarity of this compound to other structures, demonstrates its uniqueness among natural products and the potential of this technique to access new chemical space, a high priority for natural product research. The intriguing biological activities associated with cabrillostatin (**1**), including effects on cardiomyocyte beat frequency and amplitude, demonstrate the applications of this approach to natural product drug discovery. Cabrillostatin also induces phenotypic changes in cancer cell lines that are distinct from reference compounds suggesting a unique mode of action. Interestingly, cabrillostatin is widespread in seawater and represents a now characterized component of DOM that can be studied in an ecological context and used to facilitate the identification of related compounds. While our efforts to link these compounds to their biological origins remain ongoing, further advances in sequencing technologies coupled with the targeted isolation of specific taxa (e.g., planctomycetes) will facilitate new connections between chemical and biological diversity.

While the SMIRC technique will benefit from further refinements, we show that compounds can be recovered in sufficient quantities for NMR-based structure elucidation and bioactivity testing. Some compounds were detected at the same location over time, suggesting that scale-up deployments represent one approach to address expected supply issues. Incorporating the resin in agar to support in situ microbial growth represents a further advance that led to the isolation of aplysiopsene A, a compound previously reported from a marine sacoglossan (25). While synthesis represents a tractable supply route for compounds such as cabrillostatin, more complex structures such as the cabrillospirals will require a larger investment from the broader synthetic community. Ultimately, supply issues will need to be addressed on a compound-by-compound basis and will likely provide additional opportunities to develop new approaches to access chemical diversity.

In summary, the Small Molecule In situ Resin Capture Technique (SMIRC) provides a unique approach to access chemical space that has remained beyond reach using current methods. It is agnostic of the producing organism and can be used to isolate novel natural products directly from Nature in sufficient quantities for NMR based structure elucidation and biological testing. This compound first approach inverts the traditional paradigm of microbial natural product discovery thus circumventing many of the major bottlenecks associated with cultivation and genome mining. By incorporating up front bioassays, bioactive compounds can be directly targeted without the need to build large culture collections. Finally, SMIRC is relatively simple to assemble and deploy, requiring the type of analytical instrumentation readily available at most research-intensive universities. Moving forward, this approach can be expanded to other environments, provide opportunities to explore the ecology of chemically mediated interactions, and be used to build unique chemical libraries for drug discovery and other commercial efforts that benefit from natural product diversity.

## Materials and Methods

### HP-20 resin processing

HP-20 resin was placed in MeOH and gently shaken at 40 rpm for 1h then filtered (Büchner funnel-filter membrane, 200 µm) under reduced pressure and washed with MeOH (x3) then H_2_O (x3). HPLC grade MeOH and MilliQ grade H_2_O were used throughout, and all glassware acid washed and rinsed with MeOH before use. Bulk washed resin could be stored in water at room temperature for several weeks before deployment at which time 100 g was packaged between two layers of Nitex™ nylon mesh (120 µm, Genesee Scientific, El Cajon, CA) and framed within a 30 cm wooden embroidery hoop (Fig. 1A). Packaged resin was secured in the field using aluminum tent stakes or rocks and processed upon recovery by washing with deionized water then transferring the resin into a glass fritted Büchner funnel (200 µm pore size), rinsing with H_2_O, and eluting three times with 200 mL MeOH under reduced pressure. The solvent was removed on a rotary evaporator to yield a salt-free crude extract. Agar-resin matrix were prepared using the following protocol: 250 g of activated and washed HP-20 resin were added together with 33 g Instant Ocean salt mixture and 12 g agar to 1 L of deionized H_2_O and autoclaved at 121°C for 20 mins. After cooling down to 50°C the still liquid agar-resin suspension was shaken and distributed into ten 15 cm petri dishes. The solidified plates were aseptically removed from the dishes, enclosed between two layers of Nitex™ nylon mesh and framed within 30 cm wooden embroidery hoop (two plates each). To avoid microbial contamination Nitex™ mesh and embroidery hoops were sterilized with 70% isopropanol before use. Packaged containers were wrapped with sterile aluminum foil until deployment. Upon recovery, agar-resin from each container was extracted three times with 150 mL of dichloromethane-MeOH (1:1). The solvents were removed on a rotary evaporator to yield crude extracts.

Methods for chemical analyses, antibiotic screening, compound isolation, phenotypic screening, eDNA extraction, DNA sequencing, and metagenomic analyses are included in the Methods section of the SI Appendix. Detailed structural elucidation of compounds **1**-**3** is provided in the Supplementary Text section of the SI Appendix.

## Acknowledgments

We thank Linh Anh Cat and Lauren Pandori for their assistance with access to the Cabrillo State Marine Reserve (Permit #: CABR-2023-SCI-0006), Brendan M. Duggan (Skaggs School of Pharmacy, UC San Diego) for access to NMR equipment and expert guidance, Douglas A. Sweeney, Alyssa M. Demko, and Leesa J. Klau for their contributions to the early development of SMIRC. This research was supported by NIH grants R01GM085770 to P.R.J. and R21AT010493 to P.R.J. and T.F.M.

## Notes

### Competing Interest Statement

The authors have declared no competing interest.

### Summary of Updates

Several sections updated, authors updated, supplemental files updated and combined.

