## Supplementary Information for "Small Molecule *in situ* Resin Capture – A Compound First Approach to Natural Product Discovery"

**This PDF file includes:**

Supporting text

Figures S1 to S55

Tables S1 to S12

SI References

Supporting Information Text

**Detailed structure elucidation of cabrillostatin (1).** The ^1^H NMR spectrum exhibited three doublet resonances attributable to three methyl groups (δ_H_ 0.90, *J* = 6.5 Hz; 0.96, *J* = 6.7; 1.20, *J* = 6.3 Hz) positioned on methine carbons. In addition, three downfield resonances (δH 5.00, m; 4.11, ddd and 4.08, dt) were attributable to CH protons bonded to oxygen or nitrogen. The complete structure elucidation of the compound was accomplished by 2D NMR experiments (^1^H-^1^H COSY, HSQC, HMBC, Table S1). The ^1^H-^1^H COSY experiment established the two spin systems in the molecule covering major parts of the carbon skeleton (16 out of 18 carbons, Fig. S6). A spin system comprising two methyl doublets at 0.96 and 0.90 ppm, coupled to a CH at 1.64 ppm, was further coupled to geminal CH_2_ signals at 1.45 and 1.35 ppm, in turn sequentially correlated with a CH-N at 4.08 ppm then a CH-O at 4.11 ppm. The latter signal was finally correlated with another CH_2_; the geminal protons at 2.54 (dd *J* = 4.5, 17.0 Hz) and 2.36 (dd *J* = 7.7, 17.0) ppm. HMBC correlation of the latter CH_2_ to a carbonyl group at δ_C_ 173.3 ppm established the β-hydroxy-γ-amino acid residue, statine. The statine residue accounted for eight carbons, two oxygens and one nitrogen. Ten additional carbons and one oxygen remained to be assigned for completion of the backbone of the molecule. The second spin system contained one methyl doublet at 1.20 ppm that was coupled to a CH-O signal at 5.00 ppm. This downfield resonance was coupled to a CH_2_ at 1.46 and 1.60 ppm, in turn correlated with another CH_2_ at 1.32 and 1.44 ppm. The spin system extended further into the ‘methylene envelope’ at ~1.35 ppm. The distal end of this spin system comprised CH_2_ protons at 2.18 and 2.20 ppm, characteristic of a methylene α to a carbonyl. The latter signals sequentially correlated with vicinal methylene protons at 1.48 and 1.71 ppm and overlapped methylene signals at 1.35 ppm. The CH_2_ signal at 2.18 ppm exhibited an HMBC correlation to an amide carbonyl at 175.7 ppm establishing the connection of the spin system to the statine residue and a 9-hydroxy-decanonyl substructure. Both carbonyls accounted for two degrees of unsaturation (two C=O double bonds) therefore, the remaining unsaturation is a ring. Despite the absence of a long range HMBC correlation from the methine proton at 5.00 ppm to the carbonyl at 173.3 ppm (due to low sample amount), the downfield shift of the proton is supportive of an ester or lactone functionality. The MS-MS spectrum provided additional confirmation of the structure (Fig. S8). Elimination of H_2_O (fragment *m/z* 310.2377, M–18 Da) can only be explained with the presence of an unfunctionalized alcohol, i.e., the β-hydroxy group of the statine residue. Based on this structural feature and the provenance of the compound, we named the natural product cabrillostatin (**1**). The relative stereochemistry of the statine residue was tentatively assigned based on the NMR shifts in comparison with published values for statine or isostatine containing natural products (1, 2). Experimental ^1^H NMR shifts and the *J* values of the CH-3, CH-4, and CH_2_-2 resonances are consistent with a *syn* (e. g. 3*S*, 4*S* as present in pepstatin (3) and miraziridin A (1)) orientation of the NH and OH groups. The configuration of the C-13 distal stereocenter remains unassigned.

**Detailed structure elucidation of cabrillospiral A (2).** NMR experiments (^1^H, ^1^H-^1^H COSY, HSQC, HMBC, NOESY, 600 MHz, 1.7 mm microcryoprobe, CD_3_CN) were conducted with 50 µg of **2**. After pooling compound from two SMIRC extracts (combined ~100 µg of **2**) the NMR experiments (^1^H, COSY, and HSQC spectra) were repeated in CD_3_OD resulting in better peak shapes with resolved signals and *S/N*. An LR-HSQMBC (optimized for *J* = 8 Hz) experiment revealed additional correlations not observed in CD_3_CN (4). The ^13^C NMR chemical shifts were indirectly detected with heteronuclear coupled experiments, HSQC, and HMBC. Extensive analysis of the NMR data recorded in both solvents led to the structure (Tables S2-S3).

HSQC (CD_3_CN) of **2** revealed 14 methines, seven methylenes, two methyls, and five quaternary carbons. In addition, one formyl group was identified from a sharp proton resonance at 9.58 ppm. A *para*-substituted phenol was established from ^1^H, HSQC, and HMBC data with two coupled aromatic H_2_ doublets at 7.01 and 6.72 ppm (*J* = 8.0 Hz) that showed ^3^*J*_CH_ HMBC cross peaks to quaternary signal C-24 at δ_C_ 154.2 ppm (characteristic of a phenol) and to the *para* C-21 δ_C_ 133.7 indicating an aliphatic substituent at this position. Two extensive ^1^H-^1^H spin systems (*A* and *B*) were identified from COSY and HSQC NMR (Fig. S11A). The aromatic protons in the *meta* position to the phenol OH exhibited ^3^*J*_CH_ HMBC correlations to a benzylic 20-CH_2_, δ_C_ 37.3 and established a connection to the first spin system *A*. Diastereotopic CH_2_ signals (H-20a, δ 2.89; H-20b, δ 2.25) were coupled to H-19 (δ 1.89, m) and further correlated to a methyl doublet (δ_H_ 0.80, *J* = 6.8 Hz) and to 18-CH-O (δ_H_ 3.46, δ_C_ 78.3). From here, the spin system extended to a CH-O (4.20 ppm) that was vicinally coupled to a CH_2_ (H-16a, δ 2.02; H-16b, δ 1.88), which subsequently correlated to the terminal CH_2_ (H-15a, δ 2.37; H-15b, δ 1.92) of the spin system *A*.

The spin system *B* comprised 10 carbons and extended from a diastereotopic CH_2_ (H-4a 3.29 ppm, doublet *J* = 17.5 Hz; H-4b 3.10, dd *J* = 4.9, 17.5 Hz) to 12-CH-O. H-4b was vicinally coupled to a CH-O at H-5 (δ 4.44), which further correlated with CH-O (H-6, δ 4.48). This COSY cross-peak was more pronounced in CD_3_OD, where both proton resonances were better resolved (4.43 and 4.55 ppm, respectively). The system continued over CH_2_-7 to a CH-8 that was coupled to the second CH_3_ doublet of the molecule (H_3_-26, δ 1.04, *J* = 6.2 Hz) and to a deshielded CH (H-9, δ 4.13), which in turn correlated to a CH-O (H-10, δ 4.32). H-10 coupled with the shielded proton H-11b of a diastereotopic CH_2_ (H-11a, δ 2.39; H-11b, δ 1.73) and finally H-11a was correlated with a CH-O (H-12, δ 4.11).

The *A* and *B* spin systems were connected with the long-range heteronuclear correlations (HMBC). Crosspeaks of H_2_-15 and H-16a with a deshielded quaternary C-14 (δC 109.6) placed it vicinal to C-15 at the end of spin system *A*. A further three-bond HMBC correlation of H-15a to a deshielded tertiary C-13, δ 63.2) connecting it with the quaternary C-14. On the other side, H-13 exhibited HMBCs to C-14 and to C-12 providing evidence for the vicinity of the both CH-12 and CH-13 despite a missing COSY correlation and finally connected the spin systems *A* and *B*. The chemical shift of C-14 is characteristic for the spiro-center carbon in a 5,6 spiroketal. The substructure was established by connecting tertiary carbons C-10 (δC 67.3, 6-ring B) and C-17 (δC 86.2, 5-ring C) over oxygen bridges with the spiro-center C-14. To fully elucidate the planar structure, four remaining carbons (aldehyde, methylene at δC 54.7, and two quaternary carbons), three degrees of unsaturation, and the positions of Cl and Br needed to be assigned. Both methylene protons H_2_-4 exhibited strong HMBC to a quaternary carbon C-3 at 179.4 ppm and proton H-4b to another quaternary C-2 resonance at 116.2 ppm that almost overlapped with two aromatic carbons C-23/23′ (δC 116.1) in the *ortho* position of the phenol ring D making the interpretation of the NMR data difficult. The aldehyde proton showed HMBC cross peaks with the hydroxymethyl terminus (C-27 at 54.7 ppm and the quaternary C-2). The methylene protons appeared as a broad singlet at 4.24 ppm in CD_3_CN and lacked clear 2D correlations in CD_3_CN, except a very weak HMBC to C-3. However, in CD_3_OD they appeared as an ‘AB quartet’ (δ H-27a 4.33, H-27b 4.30, *J* = 11.3 Hz) and exhibited clear HMBC correlations to quaternary C-2, C-3, and to the aldehyde C-1 with an unusual shift of 192.1 ppm indicative of an α-β unsaturated aldehyde thus locating the CH=O and the -CH_2_OH groups at C-2. The tetrahydrofuran (ring A) via oxygen bridge from C-6 to C-3 explained the deshielded resonances of both carbons (δC 87.7 and 179.4 respectively) and accounted for the last remaining degree of unsaturation. The positions of the halogens were determined based on the NMR shifts of the respective carbons, with inclusion of Cl within the spirobicyclic ring, and the analysis of the MS/MS fragmentation. In the high-resolution MS/MS spectrum we observed two major fragments with characteristic isotopic patterns characteristic for the chlorine containing *m/z* 277.0991 (calcd for C_16_H_18_ClO_2_, 277.0990, 0.36 ppm) and bromine containing *m/z* 287.0268 (calcd for C_12_H_16_BrO_3_, *m/z* 287.0277, -3.13 ppm) parts of the compound (Fig. S10). No fragments containing both halogens (other than those resulting from multiple losses of H_2_O) were observed, likely indicating a considerable spatial distance between the locations of the two halogens within the molecule. Even though it is challenging to predict how such a complex molecule would fragment in the ESI-MS, the calculated formulas and the associated degree of unsaturation allowed us to propose the structures of the major MS/MS fragments. The Br was assigned to the CH-9 (δC 65.7) and the Cl to CH-13 (δC 63.2); thus, securing the entire planar structure of **2**.

**Stereochemical assignments of cabrillospiral A (2).** The stereochemical elucidation of structure **2**, with 11 chiral centers distributed across four independent stereogenic substructures, is a major challenge. Interpretation of the NOESY data and *J*-*J* coupling constants provided key insights. The first stereogenic system consists of the vinylogous formate ester and associated tetrahydrofuran (ring A, Fig. S11A). NOE between the formyl proton H-1 and the methylene protons H_2_-4 demonstrated the *E* configuration of the Δ^2,3^ double bond. Clear NOE correlations from the proton H-4b to both oxygenated methine protons H-5 and H-6 provided evidence of their same facial orientation and established the relative configurations in ring A given the energetically favorable equatorial orientation of the aliphatic chain. The analysis of *J*-*J* coupling constants and NOESY data provided some information on the relative configuration associated with the [5,6]-spiroketal moiety. The most energetically stable conformation of the spiroketal benefits from two anomeric effects. The small coupling constant of 3.2 Hz between H-12 and H-13 indicates a dihedral angle of nearly >60 degrees and thus axial orientations of the OH and Cl substituents. The axial H-11b exhibits a large vicinal coupling (*J* = 11 Hz) to H-10 supporting their di-axial disposition and the energetically favorable equatorial orientation of the bulky aliphatic substituent at C-10.

NMR analysis, especially scalar coupling analysis, resolved the partial stereochemistry of several structural elements in **2** that were not immediately obvious from the NOE data. Instead, we clarified our understanding of the relative conformations within each sub-structure by analysis of vicinal coupling constants (^3^*J*) in ^1^H NMR and further refined by DFT calculations of the simplified model compounds **4a**-**4b** (Fig. S12). The *cis*-1,2-disubstituted furanylidene group (ring A, vinylogous formate ester) was suggested by observation of a ^3^*J* ~ 3.5 Hz vicinal coupling constant of H-5 to H-6, and supported by DFT calculations of both *cis* and *trans* isomers of model compound **4** (Fig. S13) and Monte-Carlo searching of optimized geometries in the lowest-energy conformers (>90% of Boltzmann population). Only the *cis*-isomer, with an H-5–H-6 dihedral angles (*θ* ~38˚), satisfies the Karplus relationship requirement (electronegative substituents are considered) for the ^3^*J*_H5-H6_ ~3.5 Hz coupling constant (5). NOE correlations from both H-5 and H-6 to H-4a further support the *cis* orientation, even though no NOE between H-5 and H-6 were observed due to signal overlap.

Investigation of the spiroketal ring B-C system of **2** led to a surprising interpretation: the C-12, C-13, and C-14 (spiro-C-O) substituents must each be axial with successively alternating α,β-orientations. This counter-intuitive conclusion assumes that the thermodynamically equilibrated spiro-system, like those of almost all known [6,5]- and [6,6]-spiroketal polyketides, is stabilized by the anomeric effect: the C-O bond external to the pyran ring is invariably axial (6, 7). To test this ‘triaxial hypothesis’, model compound **5** (Fig. S12) was minimized (DFT) in a manner similar to that described above. The three lowest-energy conformers (>99% Boltzmann populations, Fig. S14) each benefit energetically from the expected anomeric effect, but they also exhibit C-12–C-14 triaxial disposition of substituents in each case. We rationalize this result by an understanding that the three polar substituents OH, Cl, and C-O at C-12–C-14 prefer conformations that minimize the net dipole of **5**, and that **5** is anchored by the anomeric effect at C-14. Unlike in some sugars, where intramolecular H-bonding plays a large role, this strong axial tendency is not compensated by vicinal hydrogen bond donor-acceptor effects (Cl is a weaker H-bond acceptor). So tenacious is this triaxial effect in [6,5]-spiroketal **5**, that the next highest energy conformer (~0.2% of Boltzman population) is the twist-boat (Fig. S14), not the tri-equatorial chair conformer. In **2**, C-12–C-14 triaxial rigidity would be reinforced by a preference of the bulky branched substituent at C-10 to lay equatorial.

Two stereo-substructures remain to be assigned: the relative configurations at C-8–C-9 and C-17–C-19, but all stereoelements must finally be united by defining the inter-stereoelement relationships to secure the complete stereostructure of **2**. This challenging task, requiring a multipronged approach that includes a combination of ^1^H and ^13^C NMR chemical shift calculation guided by synthesis of suitable models, is underway.

**Cabrillospiral B (3).** Cabrillospiral A (**2**) eluted with two minor compounds (Fig. S11B) that shared the same chromophore and were indistinguishable from **2** by MS/MS. After repeated rounds of isolation, we obtained one of the analogs in sufficient amount (27 µg) to record ^1^H NMR, HSQC, and COSY experiments in CD_3_OD. We assigned the structure of this minor isomer, cabrillospiral B (**3**), using the following reasoning: The few differences in the NMR spectra to that of **2** were found around ring A (Table S4) and the most significant were the CH-O-5 (δ_C_ 73.6; δ_H_ 4.28, dd (2.9, 5.9 Hz) vs. δ_C_ 70.5; δ_H_ 4.43 dd (3.5, 4.9)), the CH-O-6 (δ_C_ 90.9; δ_H_ 4.55 m vs. δ_C_ 88.5; δ_H_ 4.55 m), the CH_2_-7 (δ_C_ 38.9; δ_H_ a 1.55 m, b 1.76 m vs. δ_C_ 34.9; δ_H_ a 1.85 m, b 1.92 m), and minor differences observed at CH-8, CH_3_-26, and CBr-H-9. CH_2_-4 resonances were obscured by overlap with the broad residual CHD_2_OH solvent signal (low sample amount) and because of a possible deuterium exchange. Finally, the hydroxymethyl H_2_-27 appeared as a broad singlet instead of an AB quartet in **2** while the chemical shifts remained unchanged. The electronic circular dichroism (ECD) spectra of **2** and **3** showed opposite Cotton effects at *λ* 260 nm (Fig. S11C) assigned to the vinylogous ester (ring A). Taken together, the NMR data (changes in chemical shifts in ring A and at CH_2_-7) and ECD evidence prompted us to propose **3** as a diastereomer of **2**; specifically, the C-6 epimer (Fig. S11D).

**Supplementary Materials and Methods**

**LCMS analyses.** LC-HRMS was performed on an Agilent 6530 Accurate-Mass QToF with ESI-source coupled with an Agilent 1260 Infinity HPLC equipped with a degasser, binary pump, autosampler, DAD detector, and a 150x4.6 mm Kinetex C_18_ 5 µm column (Phenomenex, Torrance, CA) calibrated using the Agilent Reference Calibration Mix. Crude extracts (5 µL, 1 mg/mL) were admitted from the injector loop to the LC-HRMS and eluted (1 mL/min) using an isocratic mobile phase (20:80 acetonitrile (ACN):H_2_O and 0.1 % formic acid (FA) for 2 min followed by a gradient elution to 5:95 ACN:H_2_O over 18 min, isocratic for 2 min, then increasing to 100% ACN over 1 min, and finishing with 100% ACN for 2 min. MS data were acquired over the range 135-1700 *m/z* in positive mode using the following parameters: MS scan rate, 2/s; MS/MS scan rate, 3/s; gradient collision energy (slope 2.6, offset 15 eV); source gas temperature, 300°C; gas flow, 11 L/min. All solvents were LCMS grade. The data was manually analyzed using the Masshunter Qualitative Analysis Software B.05.00. Molecular networking and automated library searches were performed using the online workﬂow (https://ccms-ucsd.github.io/GNPSDocumentation/) on the GNPS website (http://gnps.ucsd.edu) (8) and were visualized using Cytoscape 3.8.2 (9) and the GNPS in-browser network visualizer. Spectra of interest were queried in MASST via the GNPS platform (10) and publicly available metabolomic datasets were searched with default parameters (<https://masst.ucsd.edu/>). MS/MS data can be found on the Mass spectrometry Interactive Virtual Environment (MassIVE) at <https://massive.ucsd.edu> under the identifiers MSV000091904 and MSV000091909.

**Antibiotic screening.** Crude extracts were screened for antibiotic activity using an HPLC microfractionation assay. In brief, 1 mg (i.e., 33 µL at 30 mg/mL) of crude extract was injected into an analytical HPLC (reversed-phase 150x4.6 mm Kinetex C_18_ 5 µm column, flowrate 0.8 mL/min) using a gradient from 10-100% ACN (with 0.1 % FA) over 16 min then isocratic for 4 min. Fractions (200 µL) were collected every 15 seconds starting at 1.8 minutes (system dead time) into flat bottom 96-well micro-titer plates using a Gilson FC204 fraction collector yielding 80 fractions per extract. The fractions were dried under vacuum (4 h) and the content of each well dissolved in 10 µL DMSO and 190 µL of an overnight culture of outer membrane defective *E. coli* *LptD*4213 (11) (12). Positive (10 µM chloramphenicol) and negative (10 µM streptomycin and solvent/media) controls were tested for each plate. The plates were incubated for 16 h with shaking (120 rpm, 30°C) and turbidity (OD_650_) measured before and after incubation on a Molecular Devices Emax Precision Plate Reader (Molecular Devices, San Jose, CA). Growths inhibition was calculated as the reduction in OD relative to solvent/media controls.

**Compound isolation.** For chrysoeriol, crude extracts were separated into five fractions using 2 g C_18_ reversed phase silica gel 90 Å pore size (Sigma-Aldrich, St. Louis, MO) and a 20 mL MeOH:H_2_O step gradient (25:75, 50:50, 75:25, 90:10, and 100:0 MeOH:H_2_O). HPLC was performed using Agilent’s 1100 G1312A binary pump, 1100 G1315A DAD UV/Vis detector, 1100 G1313A autosampler, and 1100 G1322A degasser (Agilent Technologies, Santa Clara, CA) coupled to a Shimadzu (Kyoto, Japan) low temperature evaporative light scattering detector (ELSD) through a Hewlett Packard 35900E (Spring, TX) interface. Active fractions were further separated by HPLC (10x250 mm C_8_ column, isocratic MeOH:H_2_O (69:31) with 0.1% formic acid (FA) and 2.5 mL/min flow. For aplysiopsene A, crude agar-resin extracts were fractionated following the same procedure with isolation guided by UV HPLC (C_18_ column, 4.6x150 mm, isocratic ACN:H_2_O (50:50) with 0.1% FA, 1 mL/min). For cabrillostatin (**1**) and cabrillospirals A-B (**2**-**3**), crude extracts were fractionated as described above and further separated by HPLC (4.6x150 mm C_18_ column, isocratic ACN:H_2_O (35:65) with 0.05% FA, 1 mL/min flow).

**NMR and ECD spectroscopy.** NMR spectra (1D and 2D) were measured at 23°C on a JEOL ECZ spectrometer (500 MHz), equipped with a 5 mm ^1^H{^13^C} room temperature probe (JEOL, Akishima, Tokyo, Japan) or on a Bruker Avance III (600 MHz) NMR spectrometer with a 1.7 mm ^1^H{^13^C/^15^N} microcryoprobe (Billerica, MA). Quantitation of sub-milligram samples was achieved through comparison with an appropriate external standard, or the ‘quantitative solvent ^13^C-satellite’ method (13). NMR spectra were referenced to the solvent signals (CHD_2_OD, δ_H_ 3.31, CD_3_OD, δ_C_ 49.00 ppm; CHD_2_CN, δ_H_ 1.94, CD_3_CN, δ_C_ 1.74 ppm). The NMR spectra were processed using MestReNova (Mnova 12.0, Mestrelab Research). Electronic circular dichroism (ECD) spectra were measured on a JASCO J-810 spectropolarimeter (JASCO, Tokyo, Japan) at 23°C in CH_3_OH and 5 mm path length quartz cells.

**Quantification of flavonoids in seawater.** One liter of seawater was collected from the seagrass meadow, passed through a Waters Oasis HLB SPE column (Waters Corporation, Milford, MA), eluted with 3 mL MeOH, and analyzed by HPLC (UV detection, 360 nm) using chromatography conditions described above for antibiotic screening (microfractionation). A methanolic solution of purified chrysoeriol (1 mg/mL) was used as a standard. AUC values were used to calculate flavonoid concentrations.

**Cell lines.** Lung (A549), ovarian (OVCAR4), prostatic (DU145), renal (786-O), and pancreatic (AsPC1) carcinoma cells were purchased from ATCC (Manassas, VA). Hepatocellular (HEPG2), glioblastoma (U87M), and colorectal (HCT116) carcinoma cells were purchased from the UCSF cell line core facility. WM164 was a gift from Dr. Helmut Schaider. A549, OVCAR4, 786-O, WM164, and AsPC1 were maintained in RPMI 1640 (Invitrogen, Waltham, MA). DU145, HEPG2, U87MG, and HCT116 were maintained in DMEM with 4.5 g/dL glucose (Invitrogen, Waltham, MA). Both media were supplemented with 10% fetal bovine serum (Gemini Bio, West Sacramento, CA) and 1% penicillin/streptomycin (Thermo Fisher Scientific, Waltham, MA). Cell lines were maintained in culture in a humidified 37˚C incubator with 5% CO_2_ for a maximum of 6 weeks. Frozen cell lines were stored in liquid nitrogen with 10% DMSO. All cell lines were routinely tested for mycoplasma as described previously (14, 15).

**Cell painting.** Reference compounds were purchased from Selleck (Houston, TX) and/or MedChemExpress (Monmouth Junction, NJ) and tested at 2 doses (1:1000 and 1:10000 dilutions). Cells were seeding into 384 well Cell Carrier Ultra plates (Perkin Elmer, Waltham, MA) at an empirically determined density in 75 µL of the respective media per well. Compounds were added directly to the well using the ECHO 650 liquid handling system (Beckman Coulter, Brea, CA) integrated into a Perkin Elmer EXPLORER G3 WORKSTATION. Transfers and plate handling were established using PerkinElmer's plate::works™ (v6.2) software.

Staining with cell painting dyes were adjusted from prior studies (16). Briefly, 10 µL of an 8x Mitotracker deep red (Invitrogen, Waltham, MA) master mix (in DMEM at a concentration of 5 µM) was added to each well, incubated for 30 min, then fixed by adding 30 µL 16% paraformaldehyde (final concentration 4%) for 30 min at room temperature (RT). Plates were then washed with 1x HBSS (Invitrogen, Waltham, MA), permeabilized with 1x HBSS+0.5% (vol/vol) Triton X-100 solution (30 min), and washed twice with 1x HBSS. Thirty µl of a staining master mix in 1x HBSS+1% (wt/vol) BSA solution (Table S1) was added and incubated in the dark at RT for 30 min. All dyes were purchased from Life Technologies. Finally, cells were washed three times with 1x HBSS, covered with aluminum foil seals, and light-protected until imaging. All pipetting steps were automated using the Perkin Elmer EXPLORER G3 WORKSTATION and performed by MultiFloFX and 405 TS washer (BioTek, Agilent Technologies).

**High-content imaging and data analysis.** Imaging was performed on the Operetta CLS confocal spinning-disk high content analysis system (Perkin Elmer, Waltham, MA) using a 20x water-immersion objective. Each well was imaged at 5 fields of view, in 5 channels (filters according to wavelength information in Table S12) in 3 z-planes. Cell segmentation and single cell feature extraction was performed using the Harmony™ software (Perkin Elmer, Waltham, MA) from maximum intensity projections of each field of view. In total ~1100 features were calculated for each cell including intensity, morphology, and texture features. Phenotypic profiles were calculated using the Kolmogorov-Smirnov (KS) statistic as previously described (17). Reference compound classification and novel compound class prediction were calculated per well based on phenotypic profiles as previously described (17). All computations were performed using Python (v3.9).

**IPSC derived cardiomyocyte differentiation.** Induced pluripotent stem cell (IPSC) derived cardiomyocytes were generated from human IPSCs (WTC background genetically modified to inducibly express dCas9-KRAB fusion protein and constitutively express GCaMP6, purchased from the Gladstone institute stem cell core facility) using previously published differentiation, expansion, and maturation protocols with modifications (18-20). Briefly, human iPSCs were dissociated into a single cell suspension using TrypLE™ Express (Thermo Fisher Scientific, Waltham, MA) and seeded into Geltrex™ LDEV-Free Reduced Growth Factor Basement Membrane Matrix-coated 12 well-plates (Thermo Fisher Scientific, Waltham, MA) at 75k cells per well in E8 media supplemented with 10 μM Y-27632. Cells were fed daily with E8 media and differentiation was initiated once the hiPSCs reached 80-90% confluence (~48h) by addition of RPMI-1640 supplemented with B27 without insulin and 7.5 μM CHIR99021. After 48h, the media was replaced with RPMI-1640 supplemented with B27 without insulin and 7.5 μM IWP2. After an additional 48h, the media was replaced with RPMI-1640 supplemented with B27-without insulin. Two days later the media was changed to RPMI-1640 supplemented with B27 containing insulin and 1% P/S that was replenished every 2-3 days. Spontaneous beating was generally observed 8-10 days after addition of CHIR99021. Induced cardiomyocytes (iCMs) from wells that showed spontaneous beating in >80% of the surface area were exposed to lactate enrichment medium and re-platted into 384 well Cell Carrier Ultra plates (Perkin Elmer, Waltham, MA) after 5 days of metabolic selection. Finally, cells were switched to maturation medium for 10 days before initiating the compound screen.

**IPSC-derived cardiomyocyte Ca^2+^ transient imaging and analysis.** IPSC derived cardiomyocyte Ca^2+^ transients were monitored using a EVOS M7000 Imaging System (Thermo Fisher Scientific, Waltham, MA) for 20 seconds at a frame rate of 30hz in the GFP channel (Excitation max 488 nm). Mean fluorescence intensities for each field of view were extracted using a custom image analysis pipeline in Python. The signal was baseline corrected using the Python package BaselineRemoval (v0.1.3) and the mean signal intensity of the last 400 frames were plotted for each compound.

**Protease inhibition studies.** SARS-CoV-2 Mpro was expressed as outlined previously (21). SARS-CoV-2 PLpro was purchased from Acro Biosystems (PAE-C518), and human TMPRSS2 was purchased from Cusabio Technology (CSB-YP023924HU). Other recombinant proteases were purchased from R&D Systems which included SARS-CoV-1 3CL/Mpro (E-718), MERS-CoV 3CL/Mpro (E-719), human cathepsin B (953-CY), human cathepsin L (952-CY), human cathepsin D (1014-AS), and human cathepsin E (1294-AS). The final enzyme concentration in each assay was 50 nM of SARS-CoV-1, SARS-CoV-2 Mpro, MERS-CoV Mpro, and SARS-CoV-2 PLpro; 0.5 nM of human cathepsin L and B, 1 nM human cathepsin D and E, and 3 nM TMPRSS2 (22). Fluorogenic peptide substrates were purchased from a variety of vendors and used at the following concentration, 10 µM MCA-Ala-Val-Leu-Gln-Ser-Gly-Phe-Arg-Lys(DNP)-Lys-NH_2_ (a gift from Charles Craik, University of California San Francisco) (23), 50 µM Mu-His-Ser-Ser-Lys-Leu-Gln-AMC (Sigma SCP0224), 50 µM z-Arg-Leu-Arg-Gly-Gly-AMC (Bachem I1690), 5 µM Mca-Gly-Lys-Pro-Ile-Leu-Phe-Phe-Arg-Leu-Lys(DNP)-DArg-NH2 (CPC Scientific, SUBS-017A), and 85.5 µM Boc-Gln-Ala-Arg-AMC (Biosynth, MQR-3135-v), 5 μM z-Phe-Arg-AMC (R&D Systems, ES009) for cathepsin L and 100 μM of the same substrate for cathepsin B. Assay buffers for protease assays were as follows, Assay Buffer 1 for SARS-CoV-1 Mpro and MERS-CoV Mpro: 50 mM HEPES pH 7.5, 150 mM NaCl, 1 mM EDTA, 1 mM DTT, 0.01% Tween-20; Assay Buffer 2 for SARS-CoV-2 Mpro: 50 mM Tris-HCl, pH 7.5, 150 mM NaCl, 1 mM EDTA, 0.01% Tween-20; Assay Buffer 3 for SARS-CoV-2 PLpro: 50 mM HEPES pH 6.5, 150 mM NaCl, 0.01% Tween-20, 0.1 mM DTT; Assay Buffer 4 for Cathepsin B: 50 mM Na-Acetate pH 5.5, 5 mM DTT, 1 mM EDTA, 0.01 % BSA; Assay Buffer 5 for Cathepsin L: 50 mM Na-Acetate pH 5.5, 5 mM DTT, 1 mM EDTA, 0.01 % BSA, 100 mM NaCl; Assay Buffer 6 for Cathepsin D and E: 50 mM Citrate phosphate buffer, and Assay Buffer 7 for TMPRSS2: 25 mM Tris-HCl, pH 8.0, 150 mM NaCl, 5 mM CaCl_2_, 0.01% Triton X-100. Cabrillostatin (**1**) was diluted into the appropriate assay buffer to a concentration of 40 μM and then pre-incubated with each protease for 15 minutes at 25 °C. Separately, the fluorogenic substrates were diluted in the appropriate assay buffers at twice the target concentration and then added to the enzyme/inhibitor mixture. All assays were performed at 25°C in triplicate wells, and DMSO was used as vehicle control. The final concentration of cabrillostatin was 10 μM. Control inhibition assays were performed with 10 µM GC373 for all 3CL/Mpro enzymes (a gift from John Vederas, University of Alberta), 10 µM of compound 159, an in house discovered PLpro inhibitor (23), 10 µM E-64 for cathepsin L and B (Sigma E3132), 10 µM Pepstatin for cathepsin D and E (Sigma P5318), and 1 µM Camostat (a gift from James Janetka, Washington University St. Louis). The final volume of each reaction was 30 µL in a 384-well plate, and fluorescence was measured at 360/460 nm (ex/em) for peptide-AMC substrates or 320/400 nm (ex/em) for internally quenched MCA-peptide-DNP substrates in a Biotek Synergy HTX fluorescence plate reader. The reaction velocity was calculated as relative fluorescent units per second. The activity was normalized to wells lacking inhibitor but containing the vehicle (DMSO) in assay buffer.

**Metagenomic analyses.** Sediment (February and March 2021) and bulk seawater samples (August 2021) were collected in parallel to the SMIRC deployments. Sediment samples (~80g) were collected in sterile WhirlPak™ bags and transported on ice to Scripps Institution of Oceanography for storage. Bulk seawater samples (2 L each) were immediately filtered (0.2 μm Sterivex). All samples were stored (-20°C) until further processing. DNA was extracted using a phenol-chloroform protocol (24) and assessed using a Nanodrop for quality and concentration estimates. When needed, low concentration extracts from the same sediment/water samples were pooled prior to sequencing. Libraries for all samples were prepared using the Nextera XT DNA library preparation kit (Illumina, San Diego, CA) for sequencing on an Illumina NovaSeq 6000 system with 150-bp paired end reads.

Raw reads were quality trimmed and adapters removed using the BBMap package (25). For taxonomic and biosynthetic analyses, samples were processed using a read-based approach to limit assembly biases (26). Briefly, single-copy phylogenetic marker genes were extracted with hidden Markov models (HMMs) (27) from each sample and classified against a reference genomic database using phylogenetic inferences with pplacer (v1.1.alpha17) (28). To identify biosynthetic domains, translated reads were searched against the NaPDoS2 (29) database for ketosynthase (KS) and condensation (C) domains. Filtered reads were normalized with BBNorm (target=40, mindepth=5) and assembled with the IDBA-UD assembler using the pre-correction flag (mink=30, maxk=200, step=10) (30). Assembled contigs (>2000bp) were searched for biosynthetic gene clusters (BGCs) with antiSMASH (v5.1.2; both fungal and bacterial versions) (31). BGCs were clustered into gene cluster families (GCFs) based on relatedness using BiG-SCAPE (v1.1.2) (32). Contigs were binned into metagenome assembled genomes (MAGs) by calculating coverage profiles with bowtie2 (33) followed by an unsupervised binning approach with Metabat2 across individual samples (34). All MAGs were checked for quality (i.e., completeness and contamination) with checkM (35), dereplicated with dRep (36), and assigned taxonomy with GTDB-Tk (37). Raw metagenomic data is publicly available under NCBI Bio Project ID PRJNA967692 and biosamples SAMN34718313-SAMN34718326.

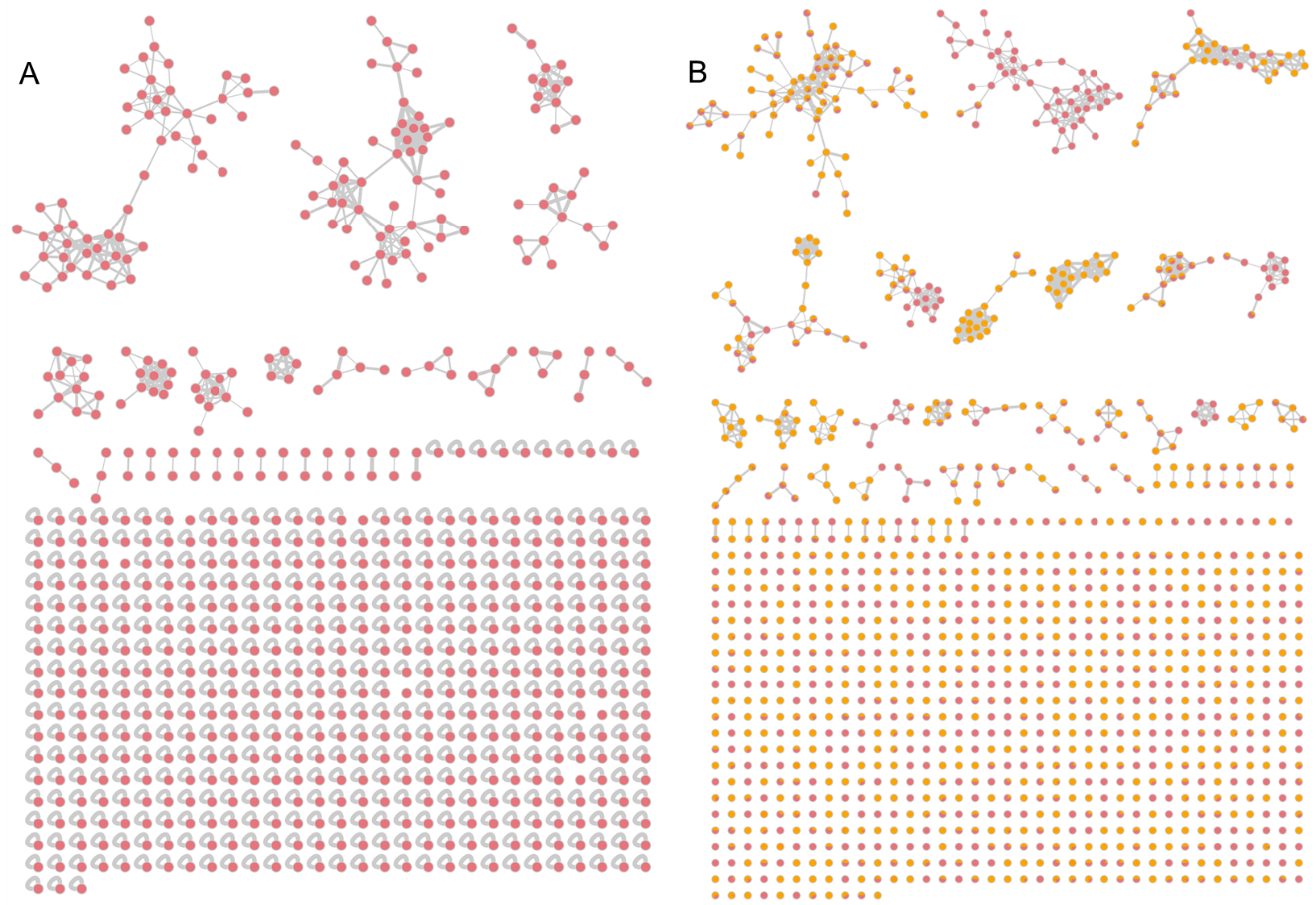

Fig. S1. Comparison of crude extract and fraction complexity using GNPS molecular networking. A. Network created with LCMS data from 15 crude SMIRC extracts (732 nodes) after rigorous subtraction of controls. B. Four of the same 15 extracts were separated into five fractions each and the results added to the network increasing the number of nodes to 1,231. Nodes are displayed as pie charts depicting relative peak intensity of ions detected in the crude extracts (red) and the fractions (orange).

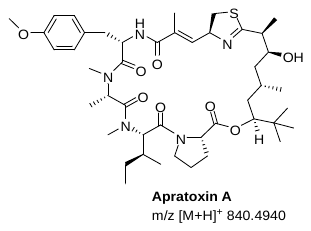

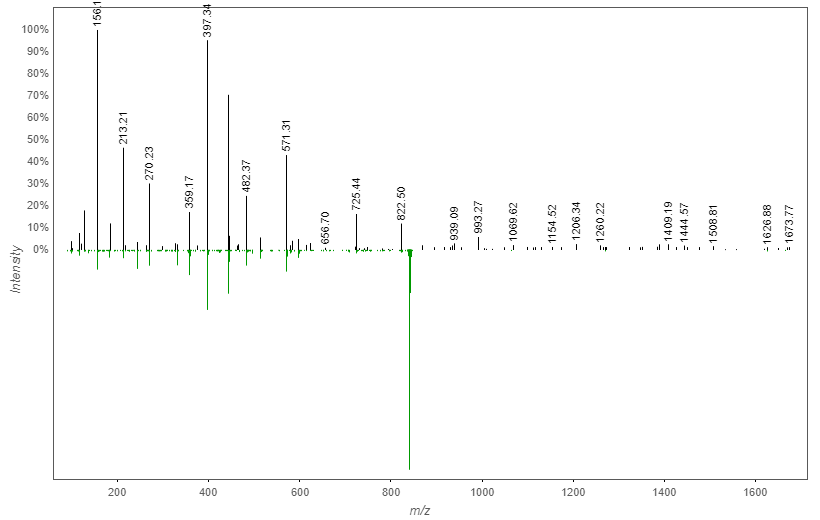

Fig. S2. A GNPS mirror-plot showing the identification of apratoxin A in a SMIRC extract from the Fiesta Island deployment site.

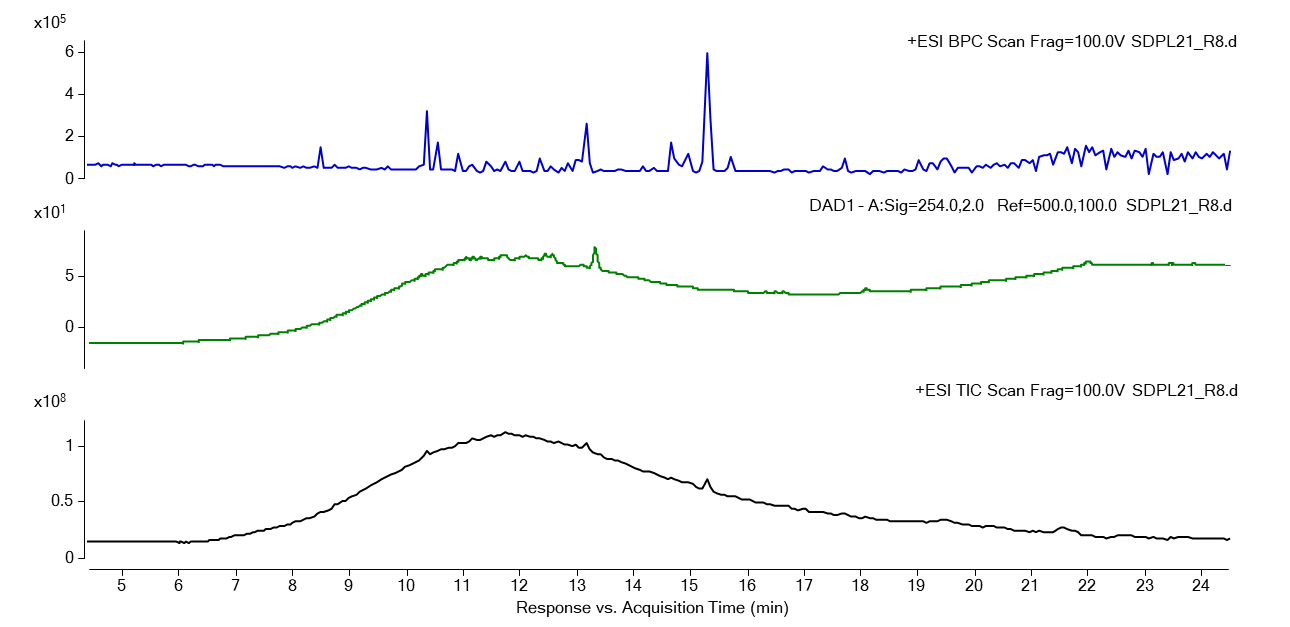

Fig. S3. LCMS analysis of a crude SMIRC extract from the CSMR deployment site. Upper chromatogram: base peak chromatogram (BPC), middle: diode array detection (DAD) UV 254 nm, lower: total ion count (TIC).

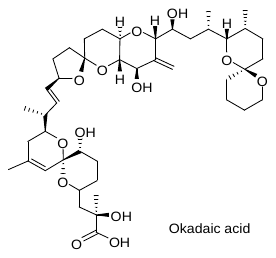

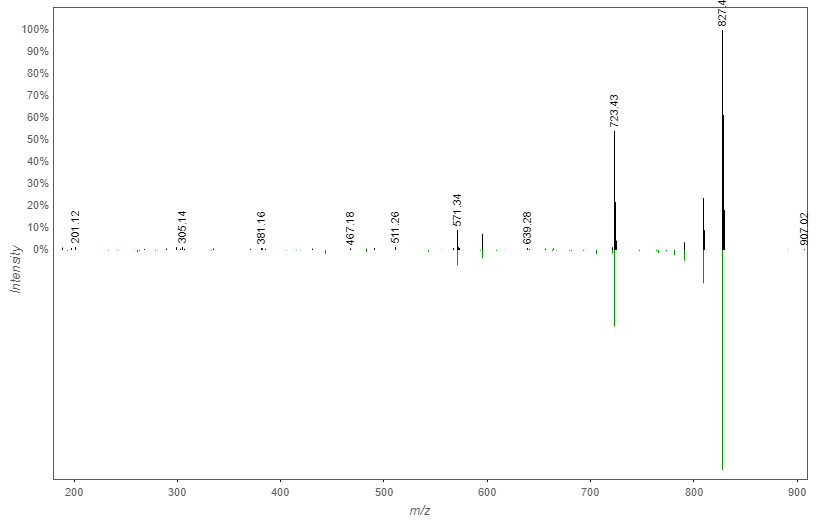

Fig. S4. A GNPS mirror-plot showing the identification of okadaic acid in a SMIRC extract from the CSMR deployment site.

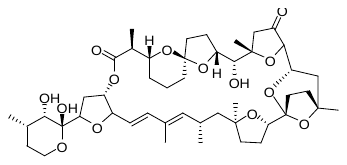

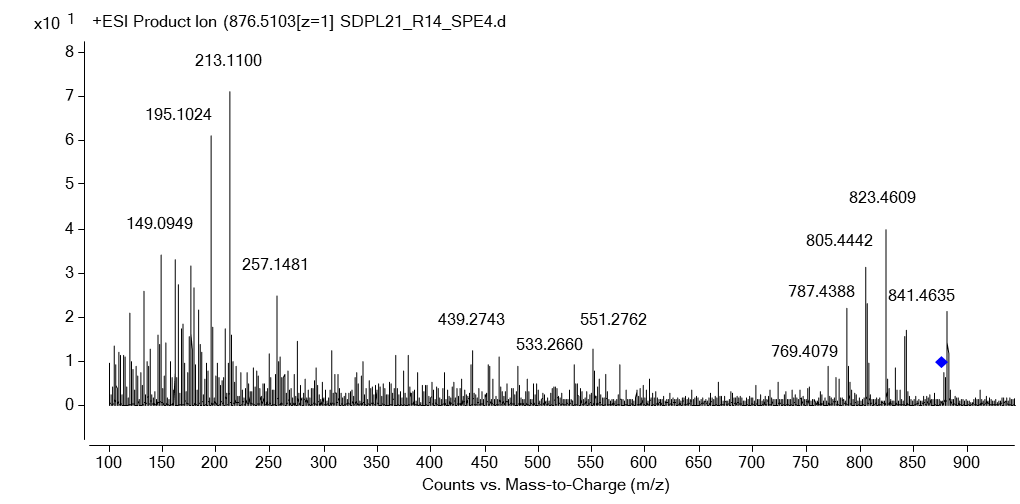

Fig. S5. MS/MS spectrum and structure of pectenotoxin-2 detected in a SMIRC extract from the CSMR deployment site. Reference spectrum can be found in Suzuki et al. 2003 (38)

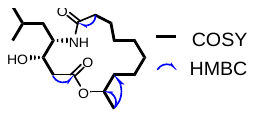

Fig. S6. Structure of cabrillostatin (1). Bold bonds indicate ^1^H-^1^H COSY correlations, blue arrows indicate key HMBC correlations.

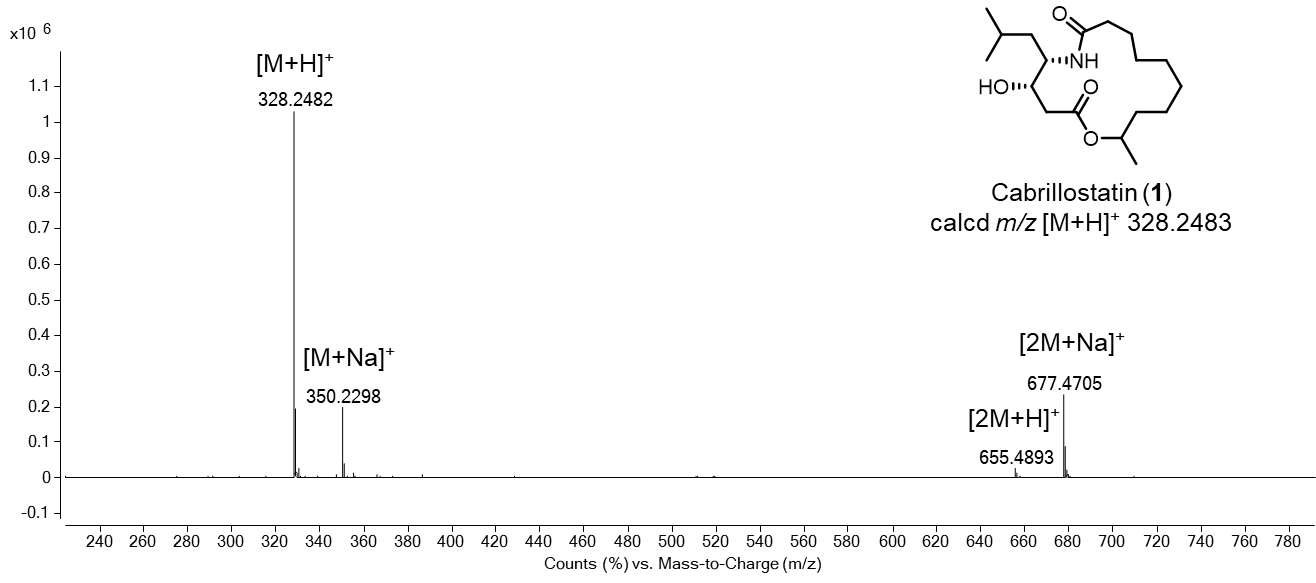

Fig. S7. HR-ESIMS of cabrillostatin (1).

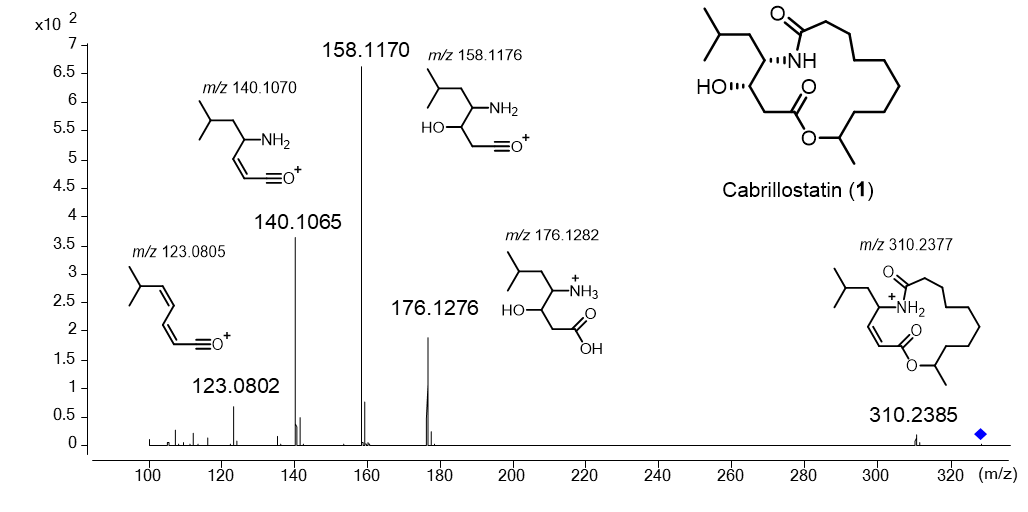

Fig. S8. HR-ESIMS/MS of cabrillostatin (1). Fragments annotated based on predicted fragmentation pattern.

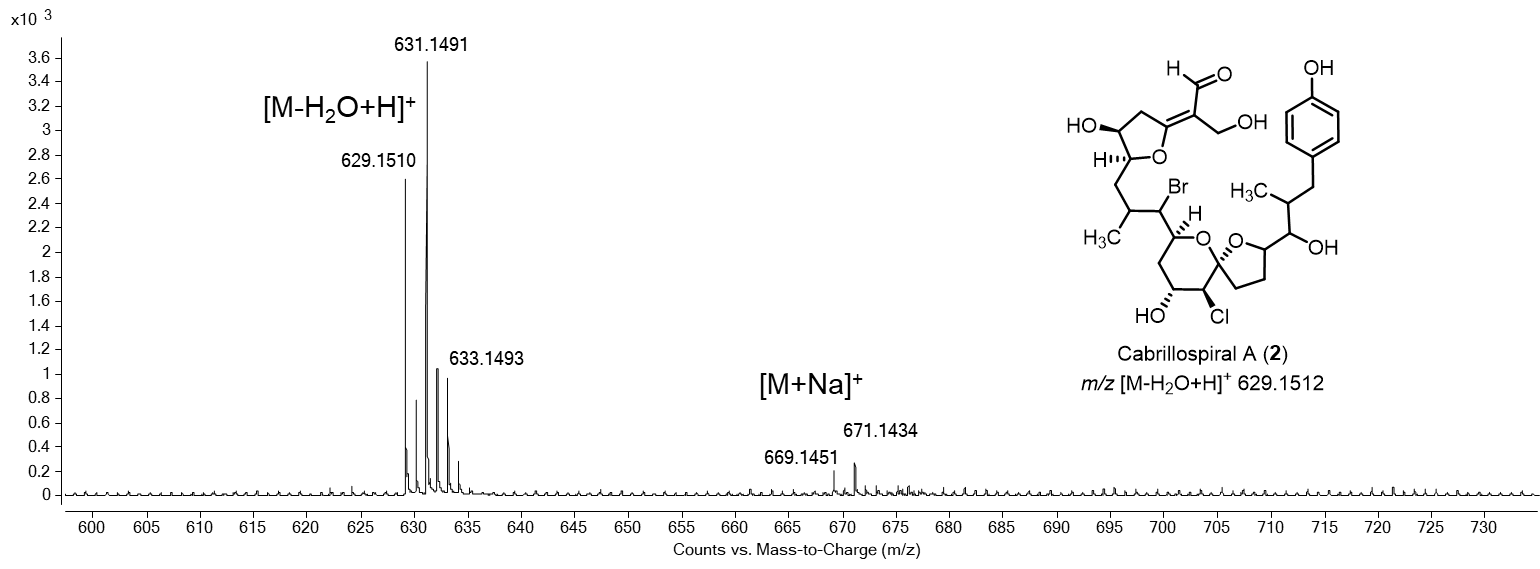

Fig. S9. HR-ESIMS spectrum and structure of cabrillospiral A (2).

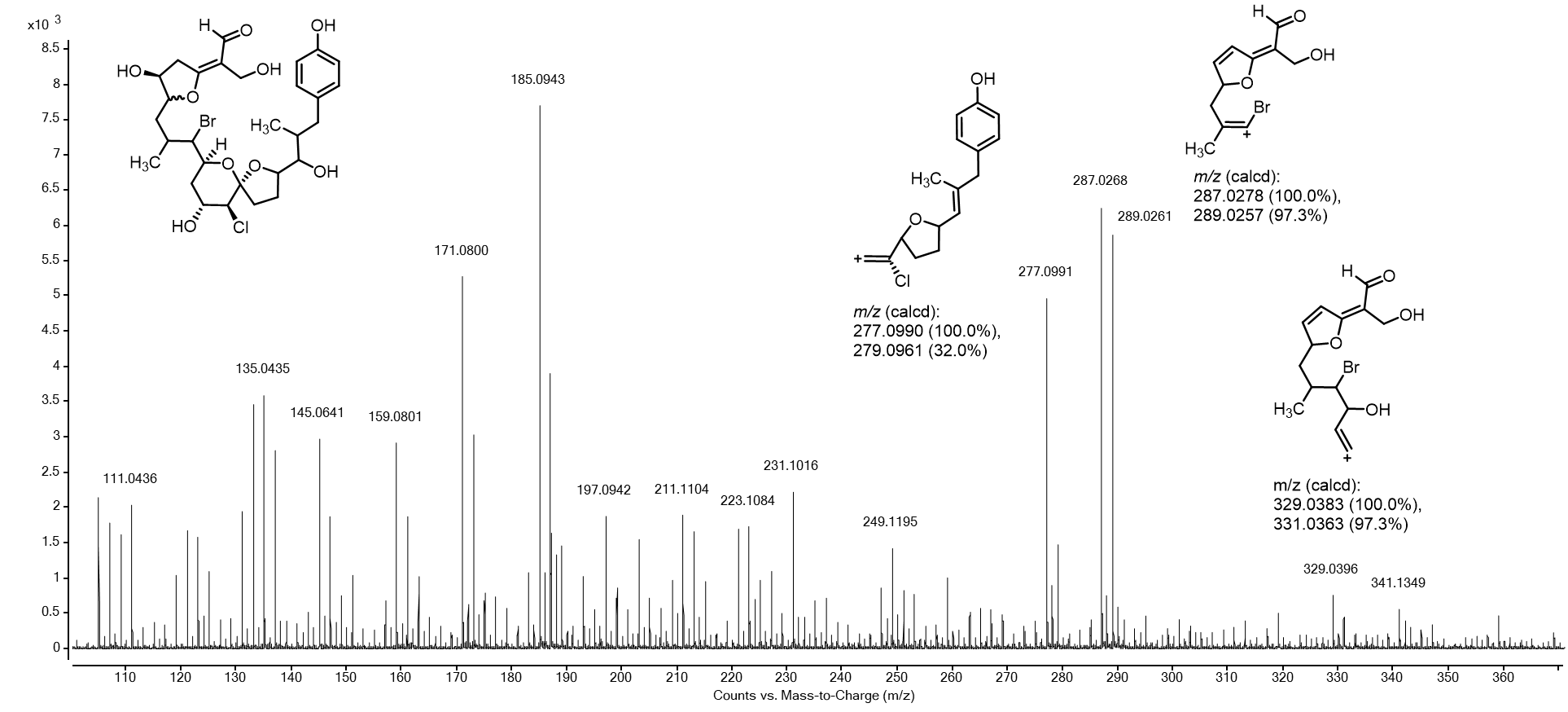

Fig. S10. MS/MS spectrum of 2 with proposed halogen containing fragments.

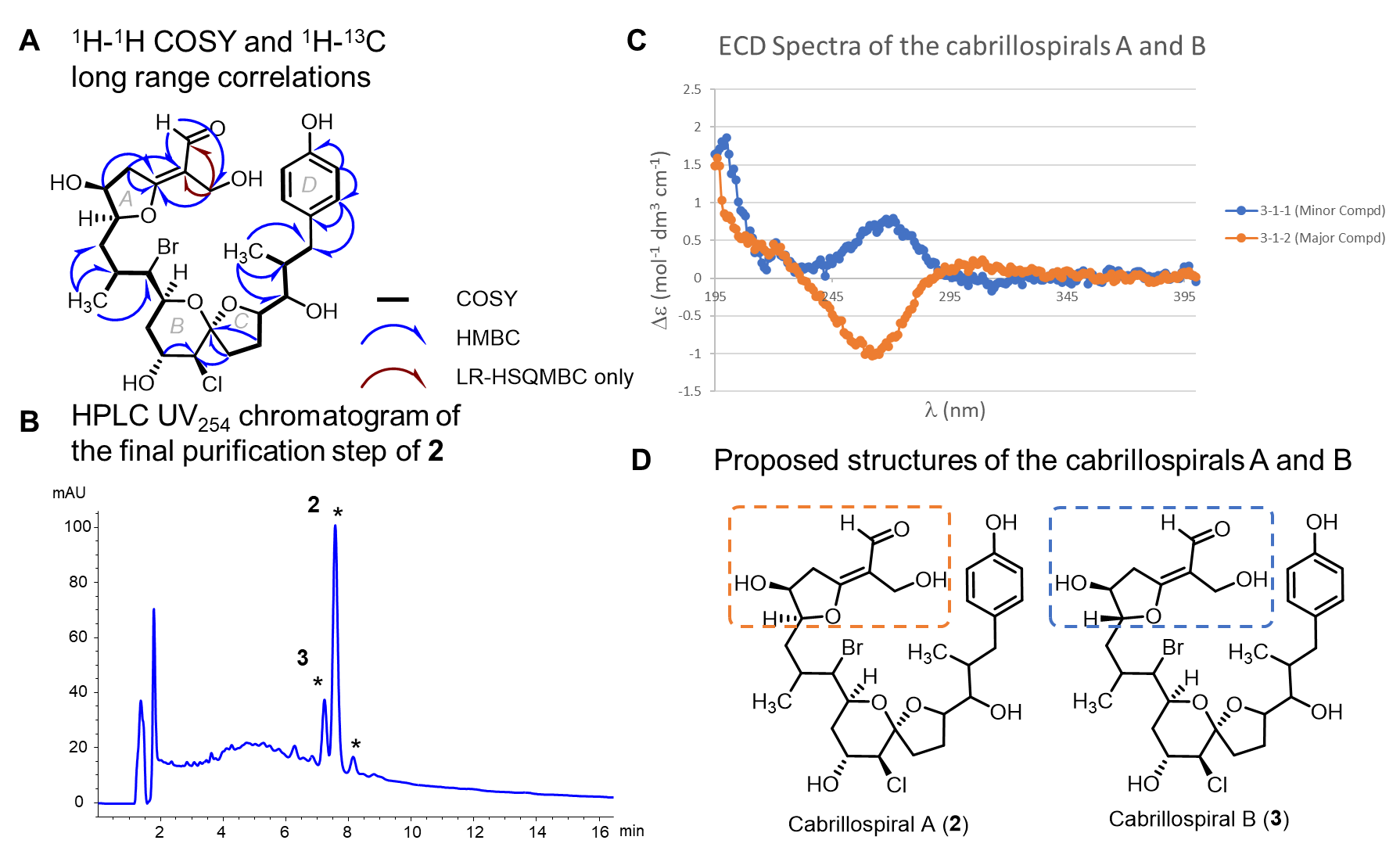

Fig. S11. Data for cabrillospirals A (2) and B (3). A. HMBC and COSY correlations. B. HPLC chromatogram from final purification step. Asterisk indicate collected peaks. C. ECD spectrum of 2 and 3. D. Proposed structures, boxes indicate the parts of the molecules that differ*.*

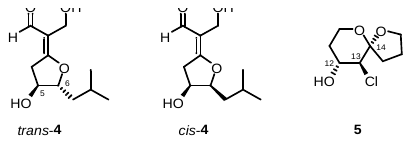

**Fig. S12.** Model compounds **4**-**5** for stereo-determination (^3^*J* and DFT calculations)
of ring A and spiroketal rings B and C in cabrillospiral A (**2**), respectively.

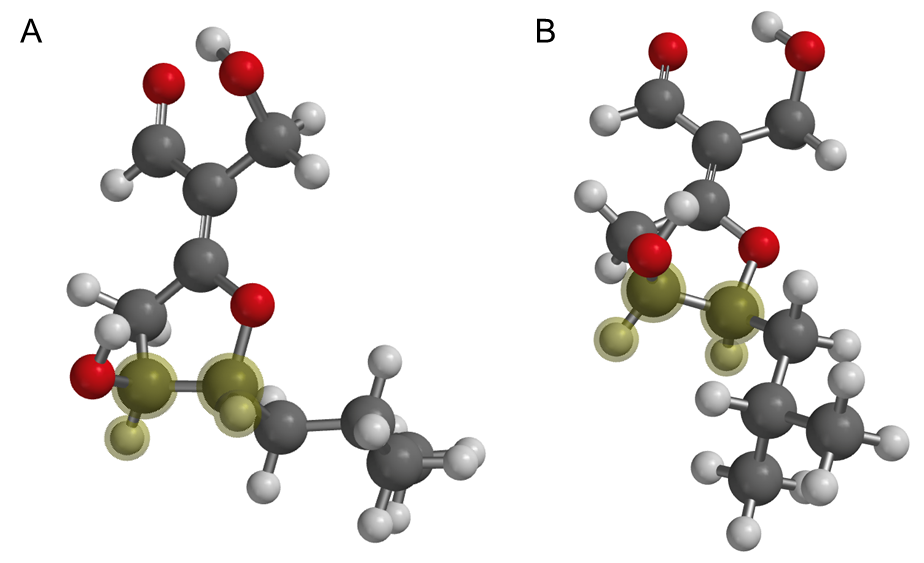

Fig. S13. DFT energy-minimized geometries of the lowest energy conformers (ωB97X-D, 6-31G*) for A *trans*-4 and B *cis*-4. Combined Boltzmann populations of the two lowest *trans* conformers are 82% and dihedral angles (*θ*_H5-H6_, highlighted) are –88.4˚ and 87.8˚. For *cis*-4, the two lowest conformers (94%) have values of *θ*_H5-H6_ = 38.1˚ and 44.3˚, respectively.

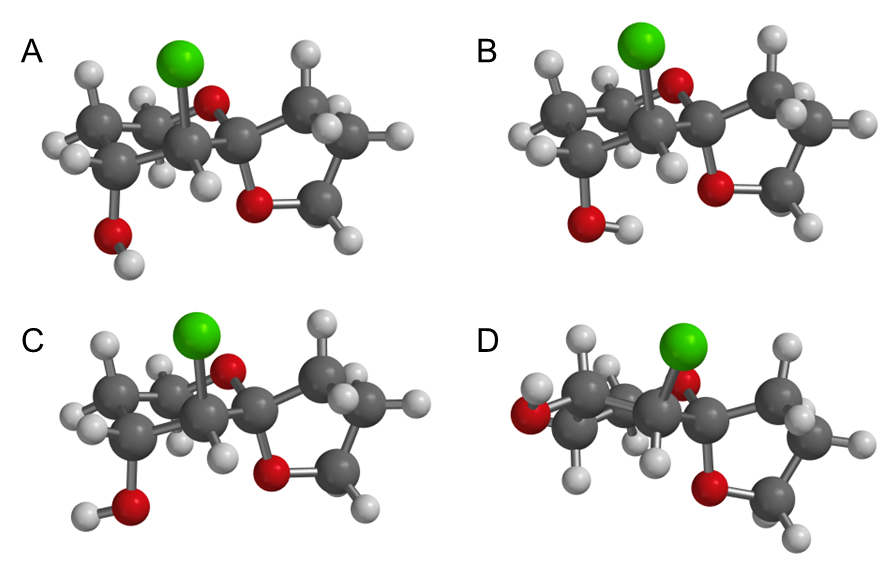

Fig. S14. Optimized geometries, minimized energies (DFT), and Boltzmann populations (%) of the four lowest energy structures of model compound 5. A. 5a E = 0 kcal.mol^–1^, (99.2%). B. 5b 3.36 (0.2). C. 5c 3.57 (0.2). D. 5d 3.75 (0.2).

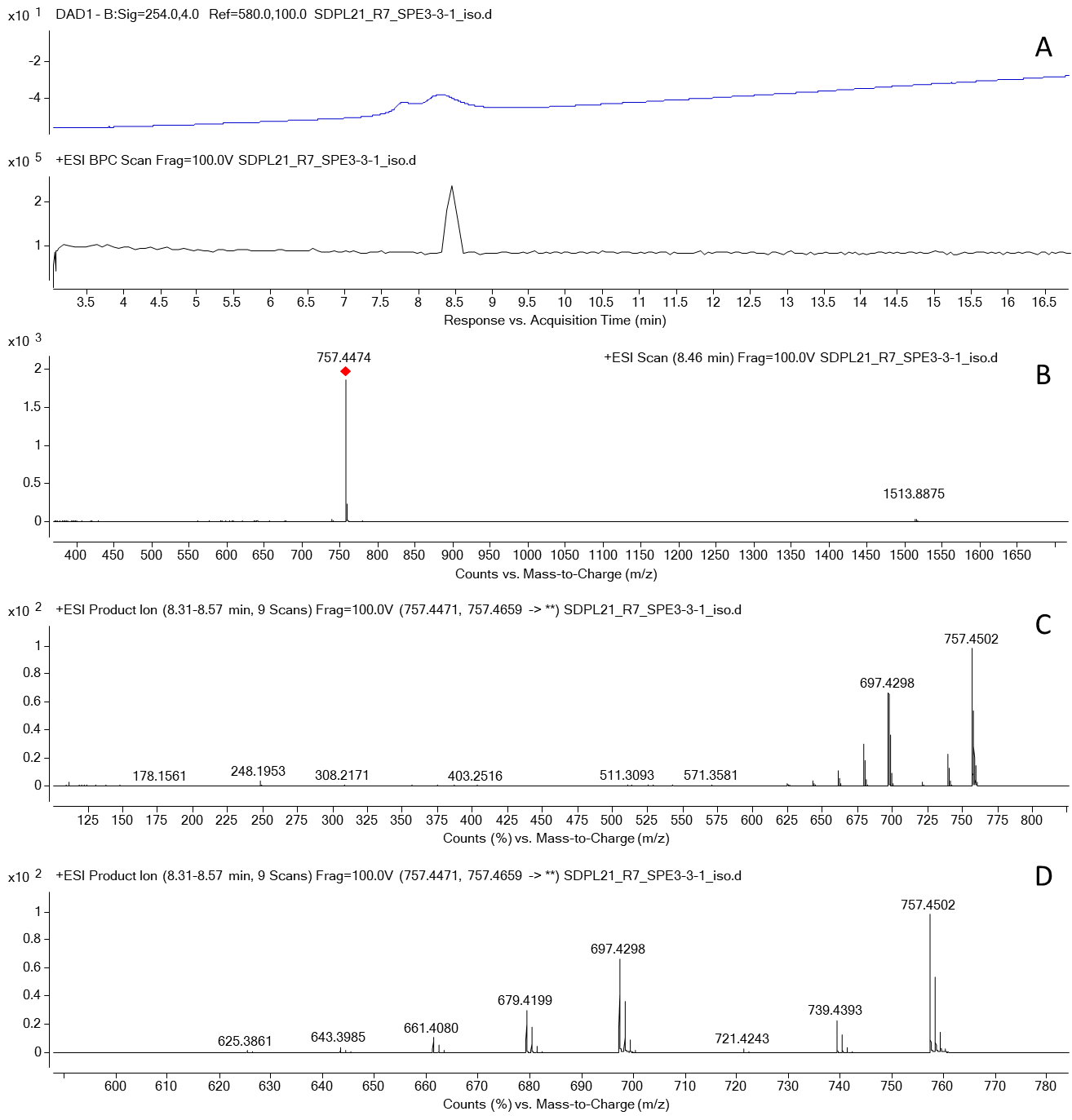

Fig. S15. LCMS of compound *m/z* 757.4474 [M+H]^+^ (calcd. formula C_43_H_64_O_11_; -6.26 ppm). A. UV 254 nm and base peak chromatograms. B. MS spectrum. C. MS/MS spectrum full. D. MS/MS spectrum expanded.

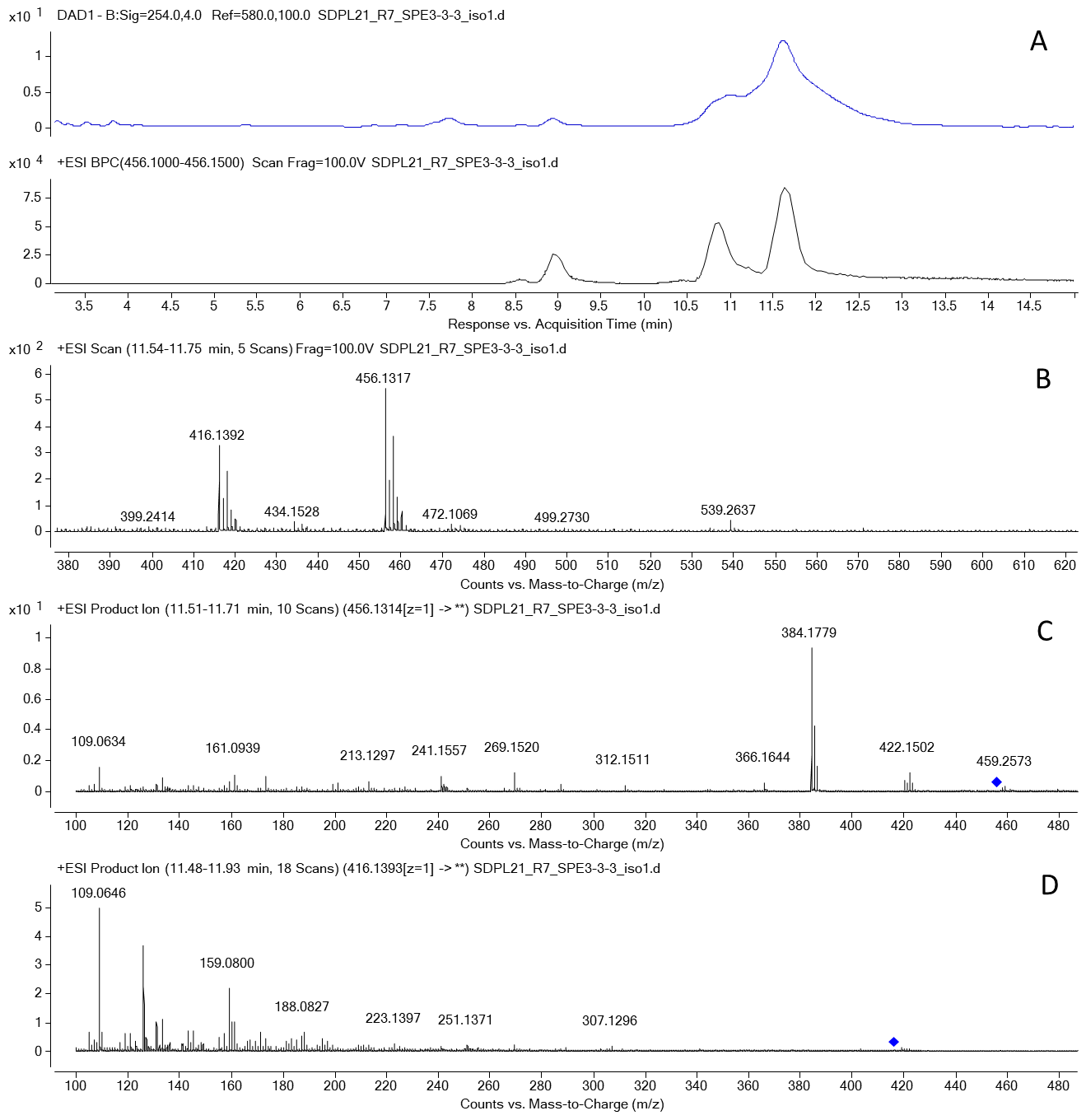

Fig. S16. LCMS of an HPLC fraction containing four *m/z* 456.1317 [M+Na]^+^ (calcd formula C_20_H_29_Cl_2_NO_5_Na; 0.44 ppm) isomers. A. UV (254 nm) and extracted ion chromatograms. B. MS spectrum of the major isomer (RT 11.7 min). C. MS/MS spectrum of *m/z* 456.1314 [M+Na]^+^. D. MS/MS spectrum of *m/z* 416.1393 [M-H_2_O+H]^+^.

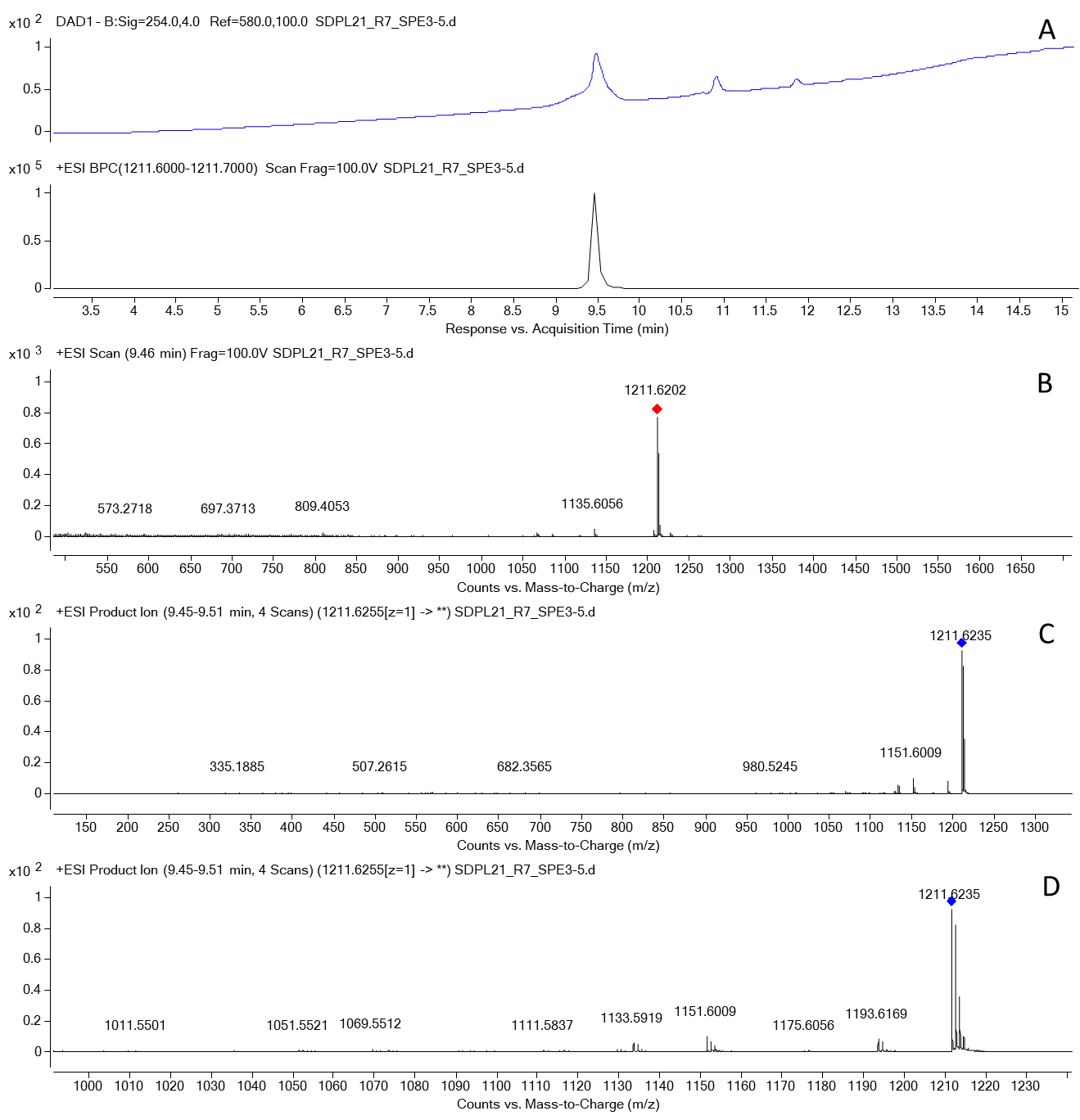

Fig. S17. LCMS of an HPLC fraction containing semi pure compound *m/z* 1211.6202 [M+Na]^+^. A. UV (254 nm) and extracted ion chromatograms. B. MS spectrum. C. MS/MS spectrum of the [M+Na]^+^ precursor. D. MS/MS spectrum expanded.

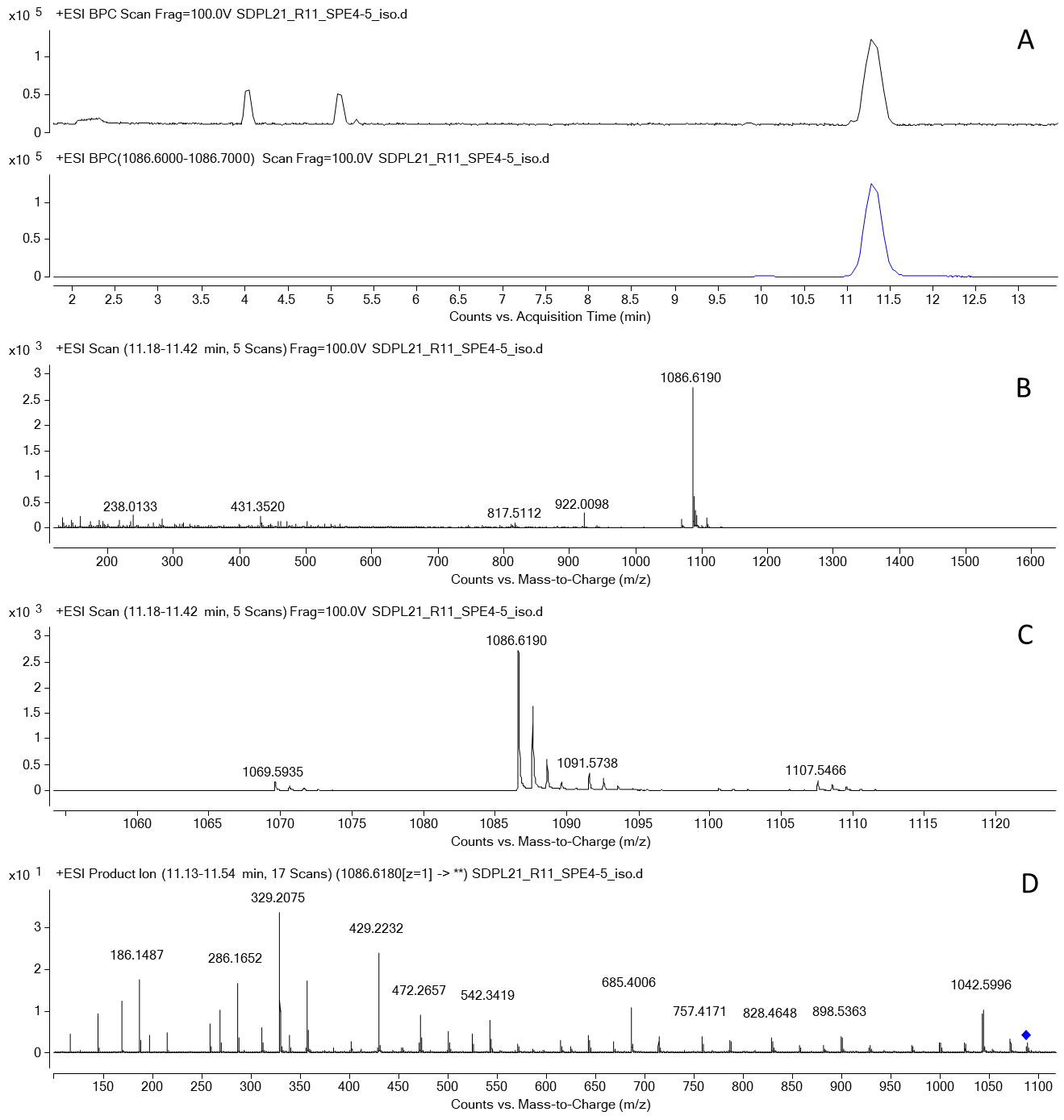

Fig. S18. LCMS of the semi pure compound *m/z* 1086.6190 [M+NH_4_]^+^ (calcd formula C_52_H_80_N_10_O_14_, -0.34 ppm). A. BPC and extracted ion (*m/z* 1089.6-1086.7) chromatograms. B. MS spectrum. C. MS spectrum enlarged; additional ions, [M+H]^+^, [M+Na]^+^ and [M+K]^+^ detected. D. MS/MS spectrum of the [M+NH_4_]^+^ precursor.

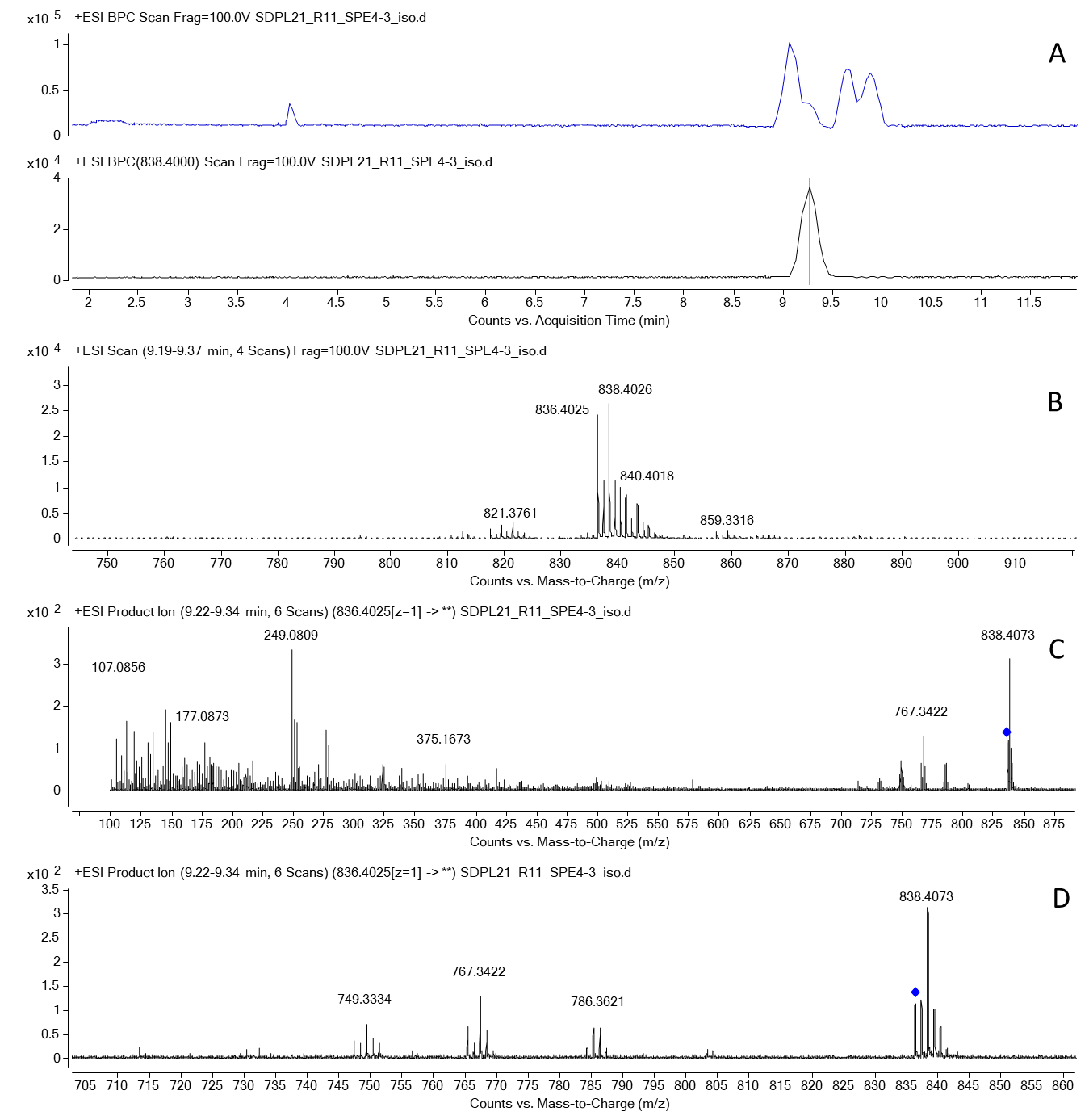

Fig. S19. LCMS of an HPLC fraction containing compounds *m/z* 1086.6190 (Fig. S18) and *m/z* 836.4025 [M+H]^+^ (calcd formula C_42_H_72_BrCl_2_NO_6_, 3.85 ppm, or C_42_H_69_Cl_4_N_3_O_5_, -4.67 ppm). A. Base peak and extracted ion (*m/z* 838.40) chromatograms. B. MS spectrum. C. MS/MS spectrum of the *m/z* 836.4025 precursor. D. Expanded MS/MS spectrum of the *m/z* 836.4025 precursor.

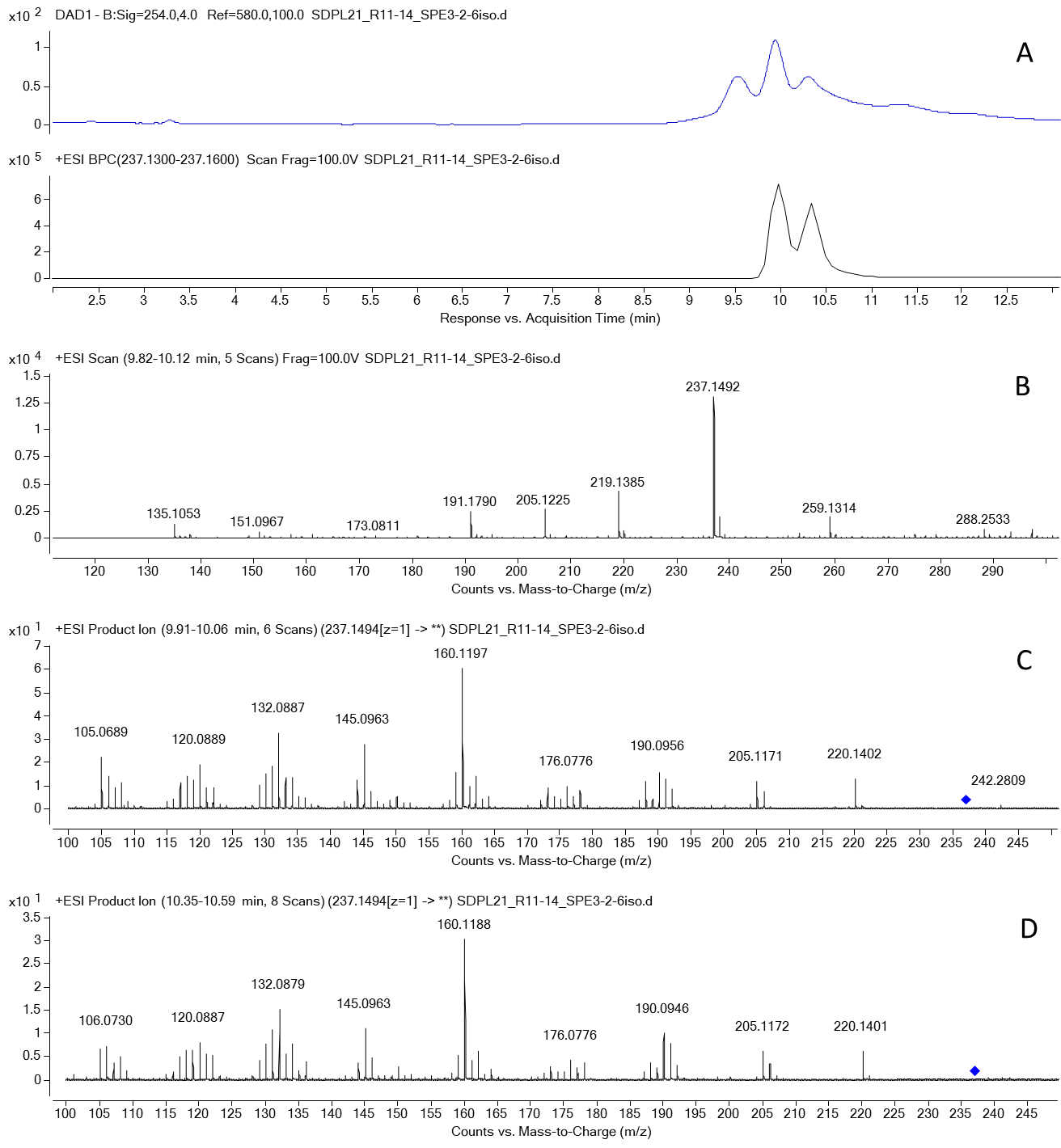

Fig. S20. LCMS of an HPLC fraction containing two compound isomers *m/z* 237.1494 [M+H]^+^ (calcd formula C_14_H_20_O_3_; 2.86 ppm). A. UV (254 nm) and extracted ion (*m/z* 237.13-237.16) chromatograms. B. MS spectrum. C. MS/MS spectrum of RT 10 min isomer. D. MS/MS spectrum of RT 10.3 min isomer.

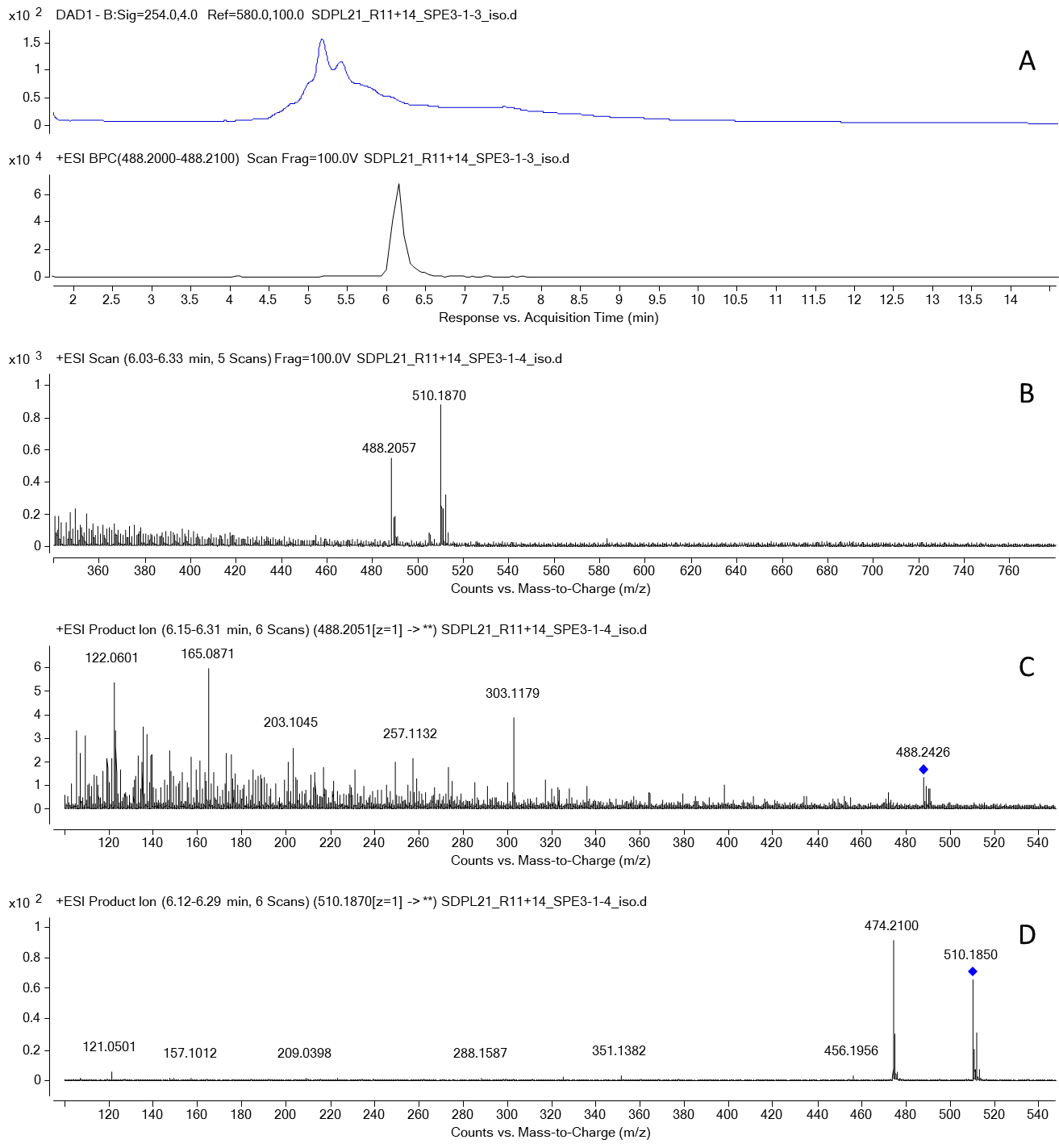

Fig. S21. LCMS of an HPLC fraction containing compound *m/z* 488.2057 [M+H]^+^ (calcd. formula C_23_H_34_ClNO_8_; -2.18 ppm) A. UV 254 nm and extracted ion (*m/z* 488.2-488.21) chromatograms. B. MS spectrum. C. MS/MS spectrum of [M+H]^+^ precursor ion. D. MS/MS spectrum of [M+Na]^+^ precursor ion.

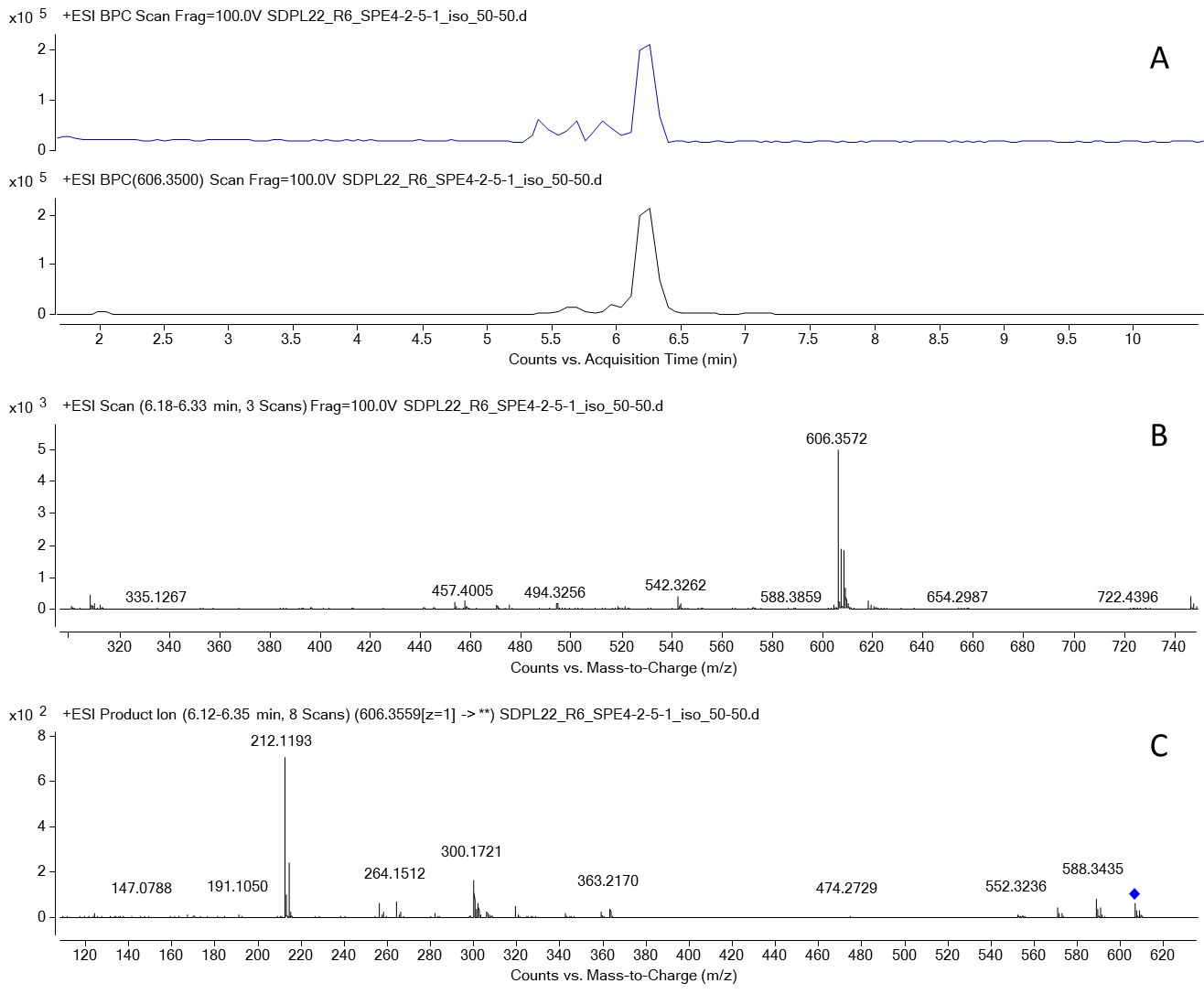

Fig. S22. LCMS of an HPLC fraction containing compound *m/z* 606.3572 [M+H]^+^ (calcd. formula C_34_H_52_ClNO_6_; 0.67 ppm) A. Base peak and extracted ion (*m/z* 606.35) chromatograms. B. MS spectrum. C. MS/MS spectrum.

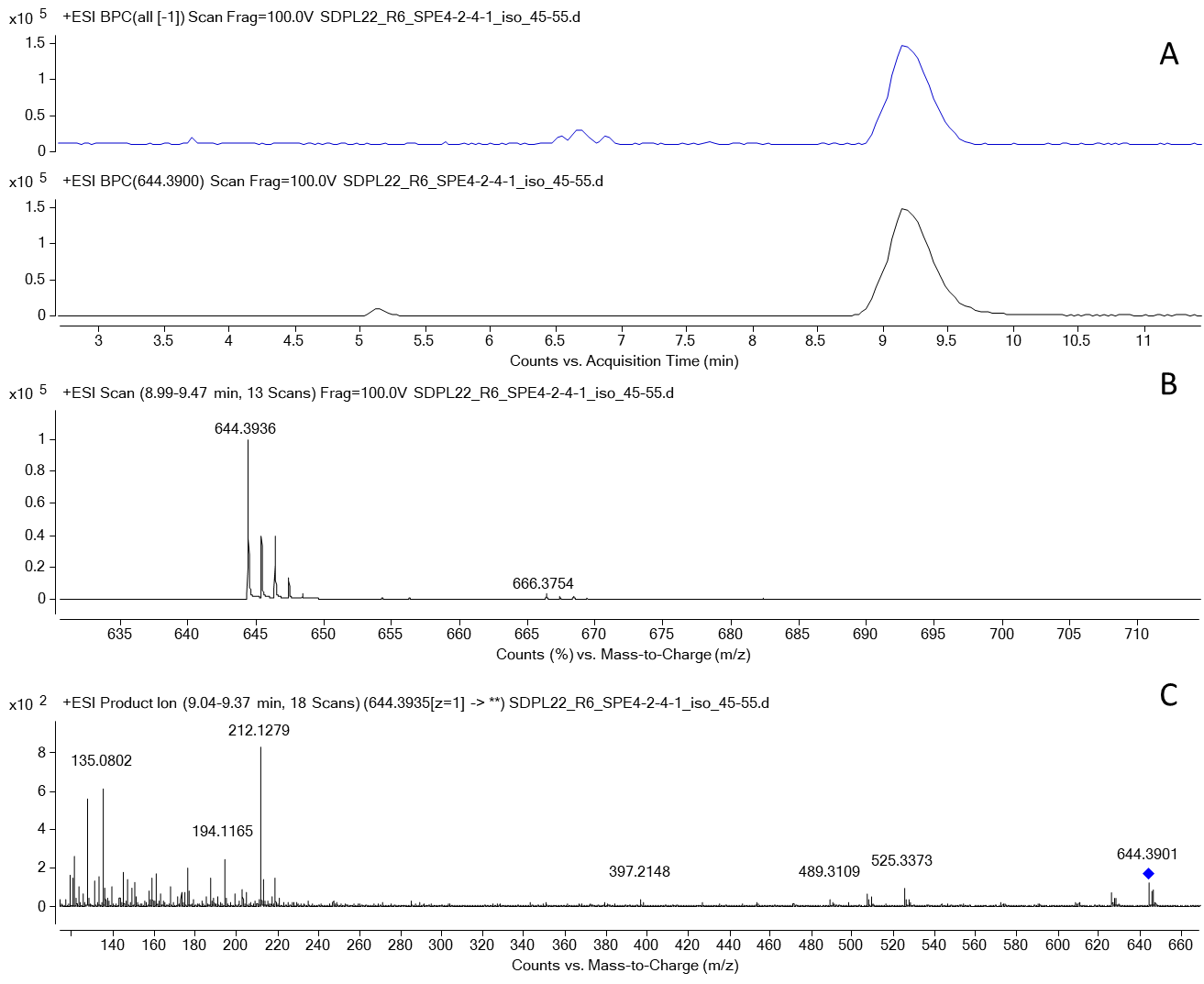

Fig. S23. LCMS of semi-pure compound *m/z* 644.3936 [M+H]^+^ (calcd. formula C_34_H_58_ClNO_8_, 1.91 ppm) A. Base peak and extracted ion (*m/z* 644.39) chromatograms. B. MS spectrum. C. MS/MS spectrum.

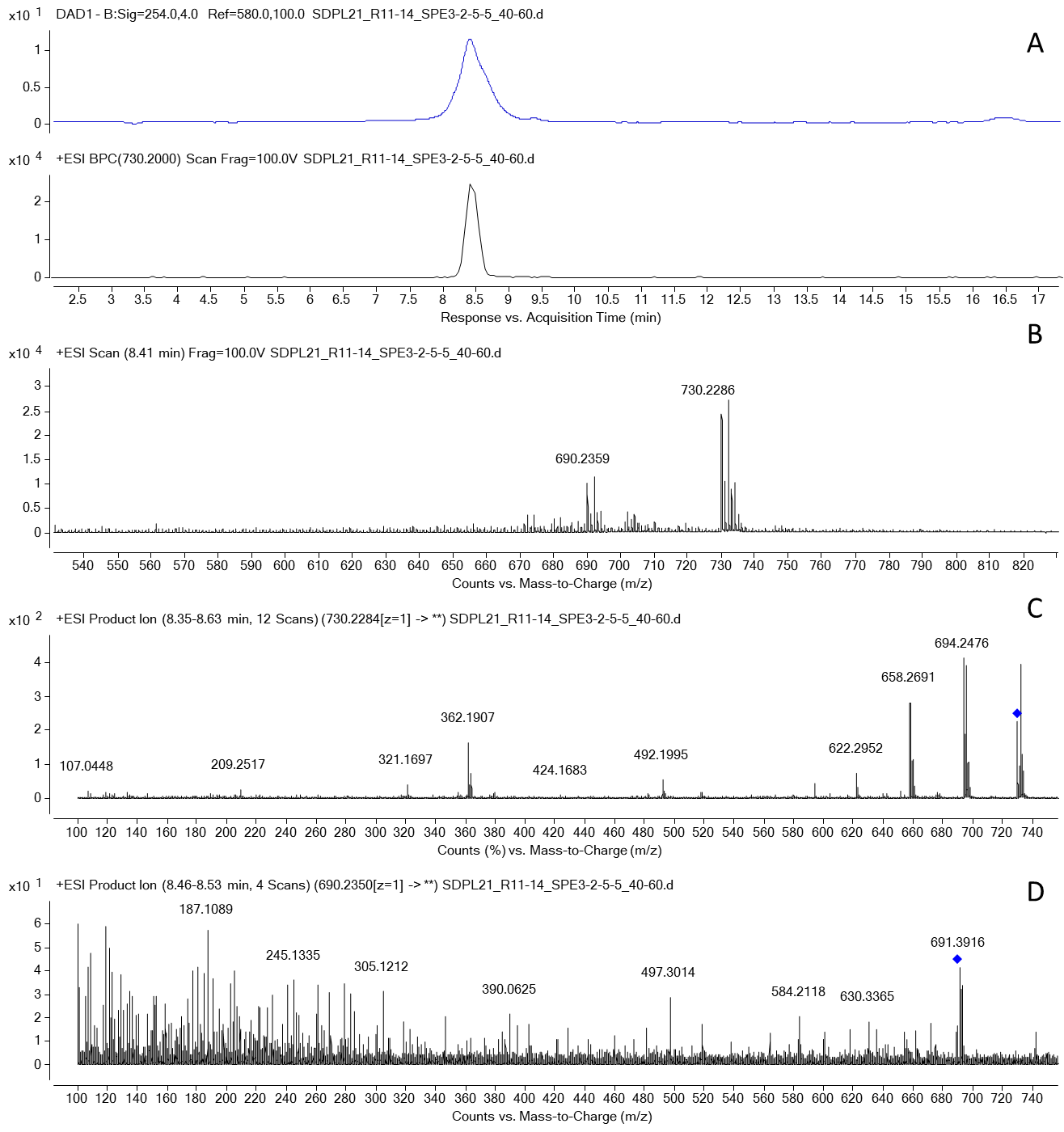

Fig. S24. LCMS of semi pure compound *m/z* 730.2286 [M+Na]^+^ (calcd formula C_33_H_48_Cl_3_NO_9_Na; -0.12 ppm). A. UV (254 nm) and extracted ion chromatograms. B. MS spectrum. *C.* MS/MS spectrum of *m/z* 730.2286 [M+Na]^+^. D. MS/MS spectrum of *m/z* 690.2359 [M-H_2_O+H]^+^.

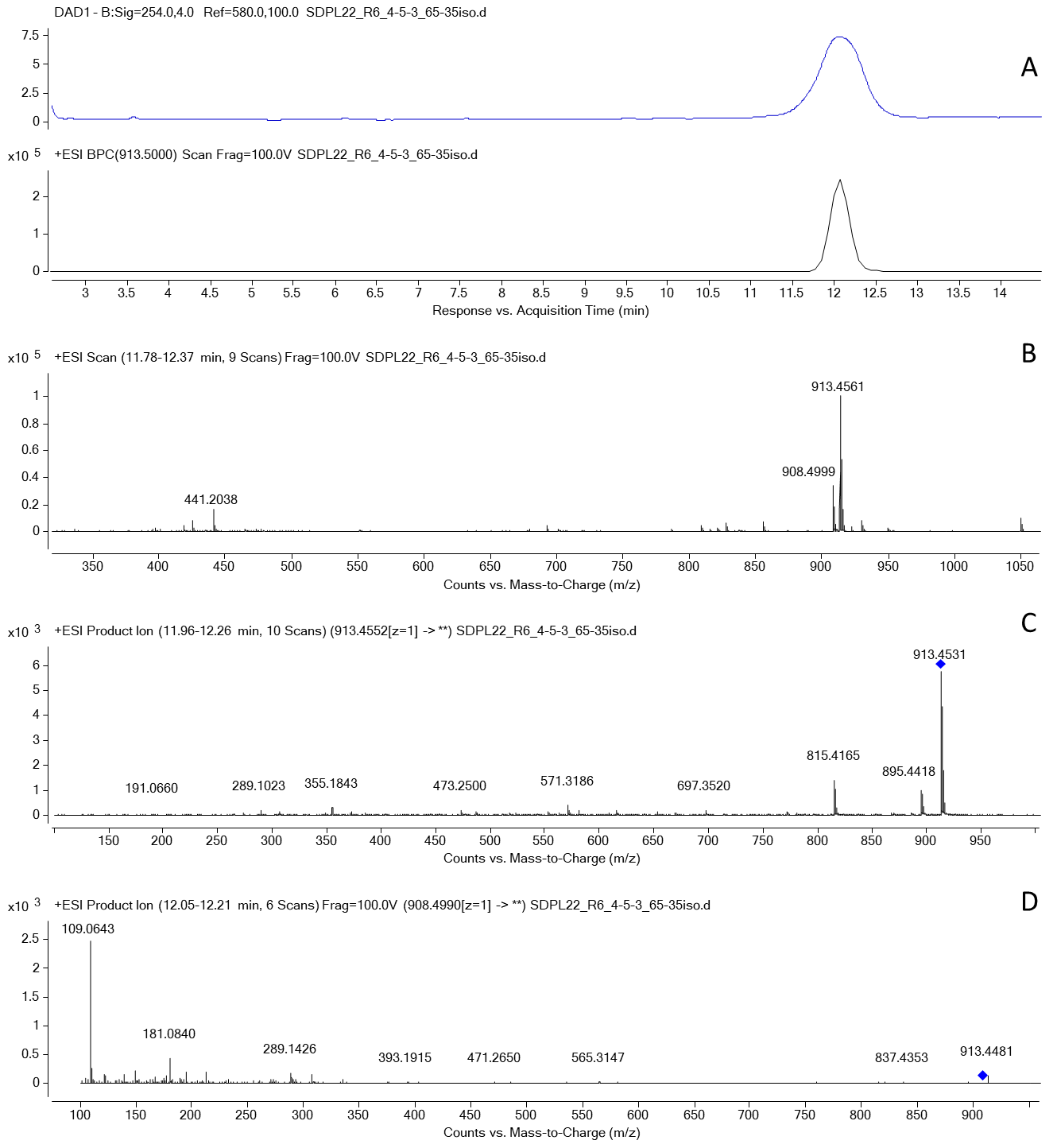

Fig. S25. LCMS of semi pure compound *m/z* 913.4561 [M+Na]^+^ (calcd formula C_47_H_70_O_16_Na; 0.54 ppm). A. UV (254 nm) and extracted ion chromatograms. B. MS spectrum. C. MS/MS spectrum of *m/z* 913.4561 [M+Na]^+^. D. MS/MS spectrum of *m/z* 908.4990 [M+NH_4_]^+^.

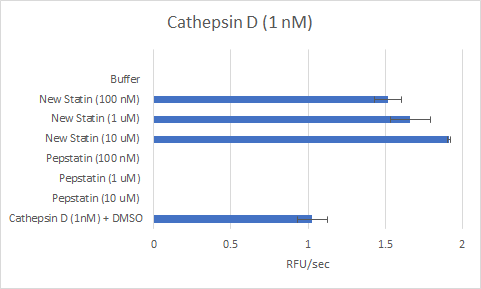

Fig. S26. Effect of cabrillostatin (1) on protease activity of cathepsin D. 1 potentiates the enzyme activity in a concentration dependent manner.

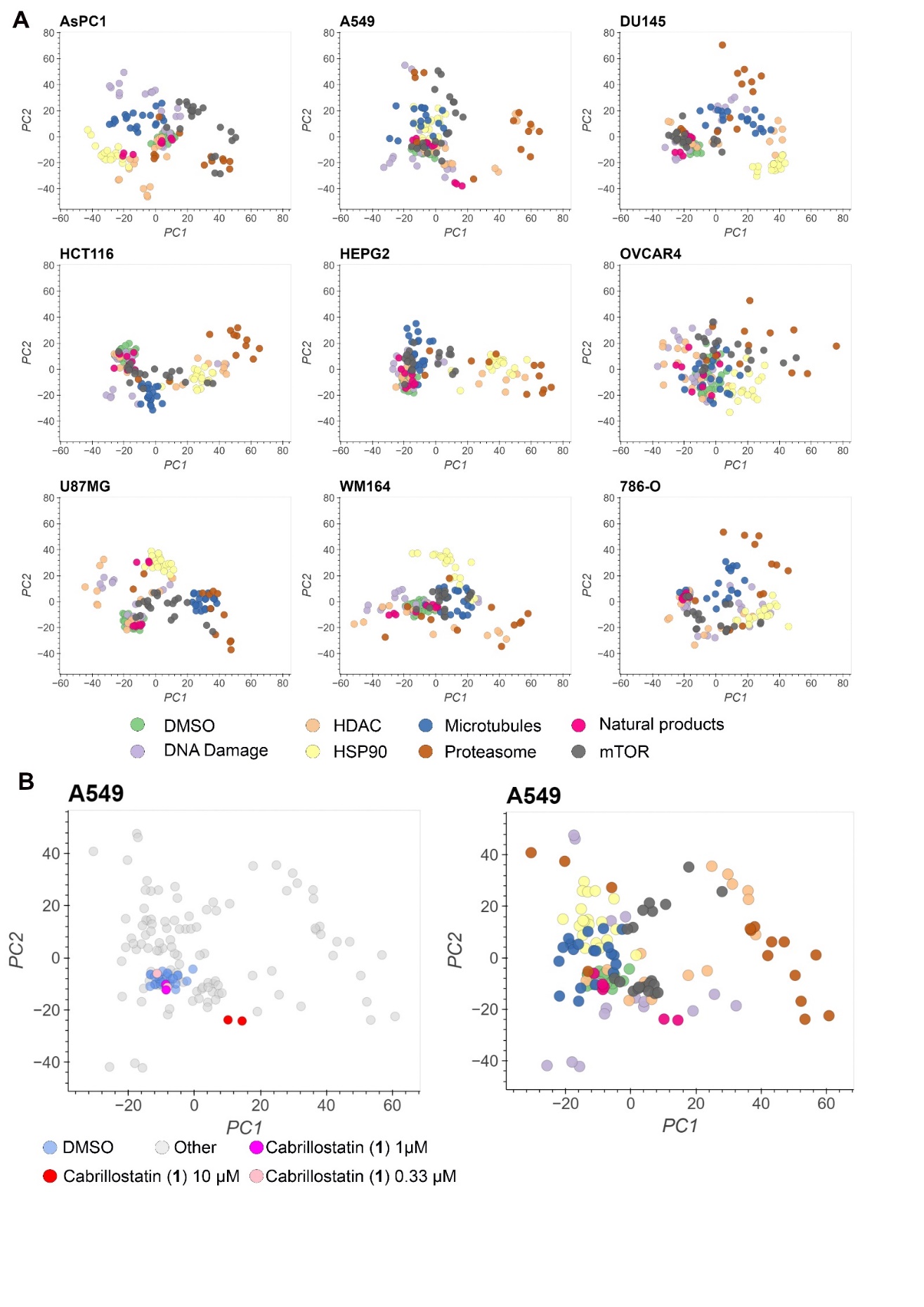

Fig. S27. Bioactivity of cabrillostatin (1) and cabrillospirals (2-3). A. Principal component analyses (PCA) of high dimensional phenotypic profiles of new natural products, reference compounds/targets, and DMSO in nine diverse cell lines (data as shown in Figure 5A). Colors indicate the different compound/target categories. B. PCA of high dimensional phenotypic profiles of a cabrillostatin dilution series. Cabrillostatin doses (left) and compound/target categories (right) are highlighted by color (color legend same as in A).

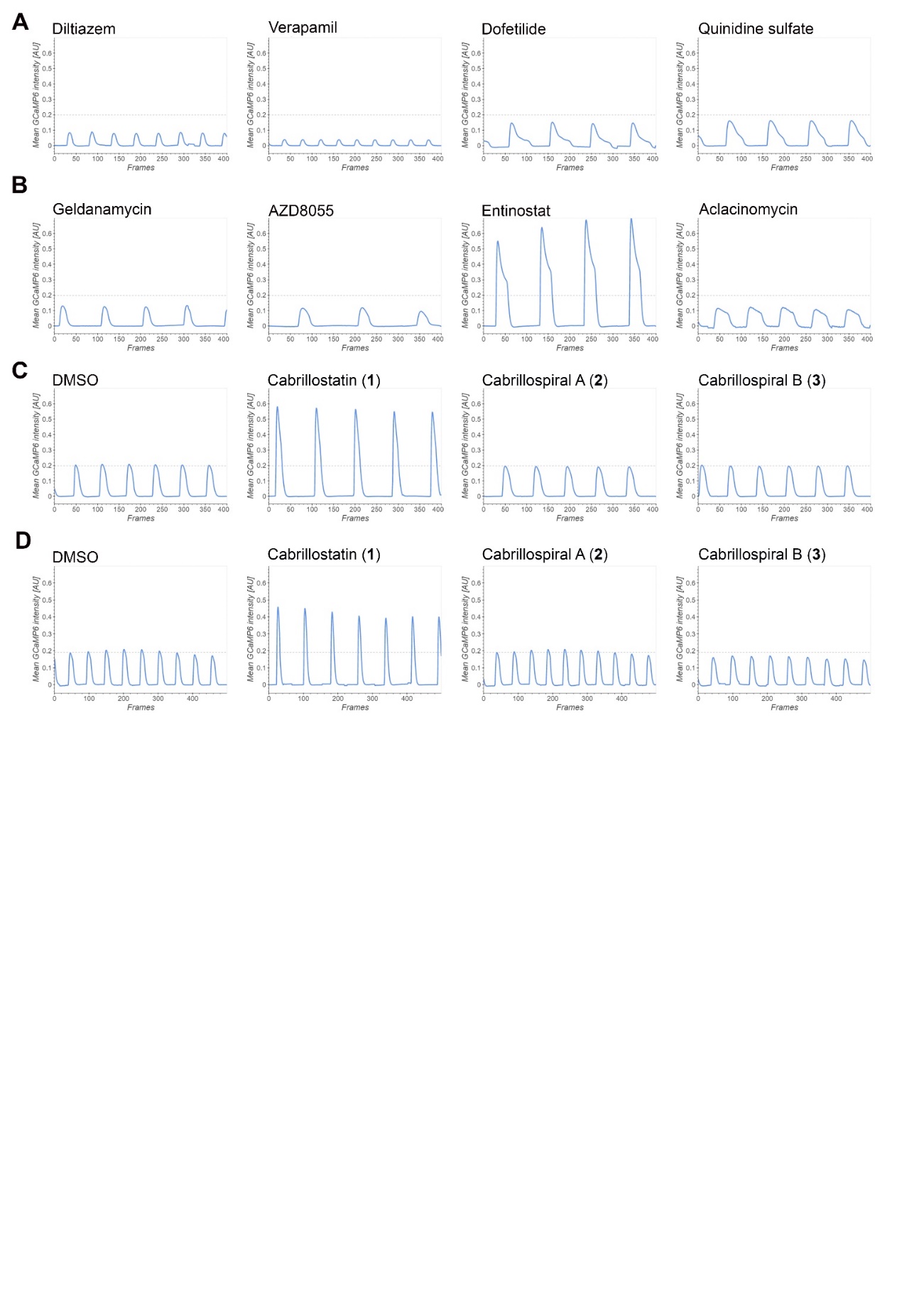

Fig. S28. Activity of cabrillostatin (1) on induced pluripotent stem cell derived cardiomyocytes (IPSC-CM). Representative Ca^2+^ transient traces of anti-arrhythmic compounds (A), anti-cancer drugs (B) or new natural products in different IPSC-CM differentiation batches (C and D).

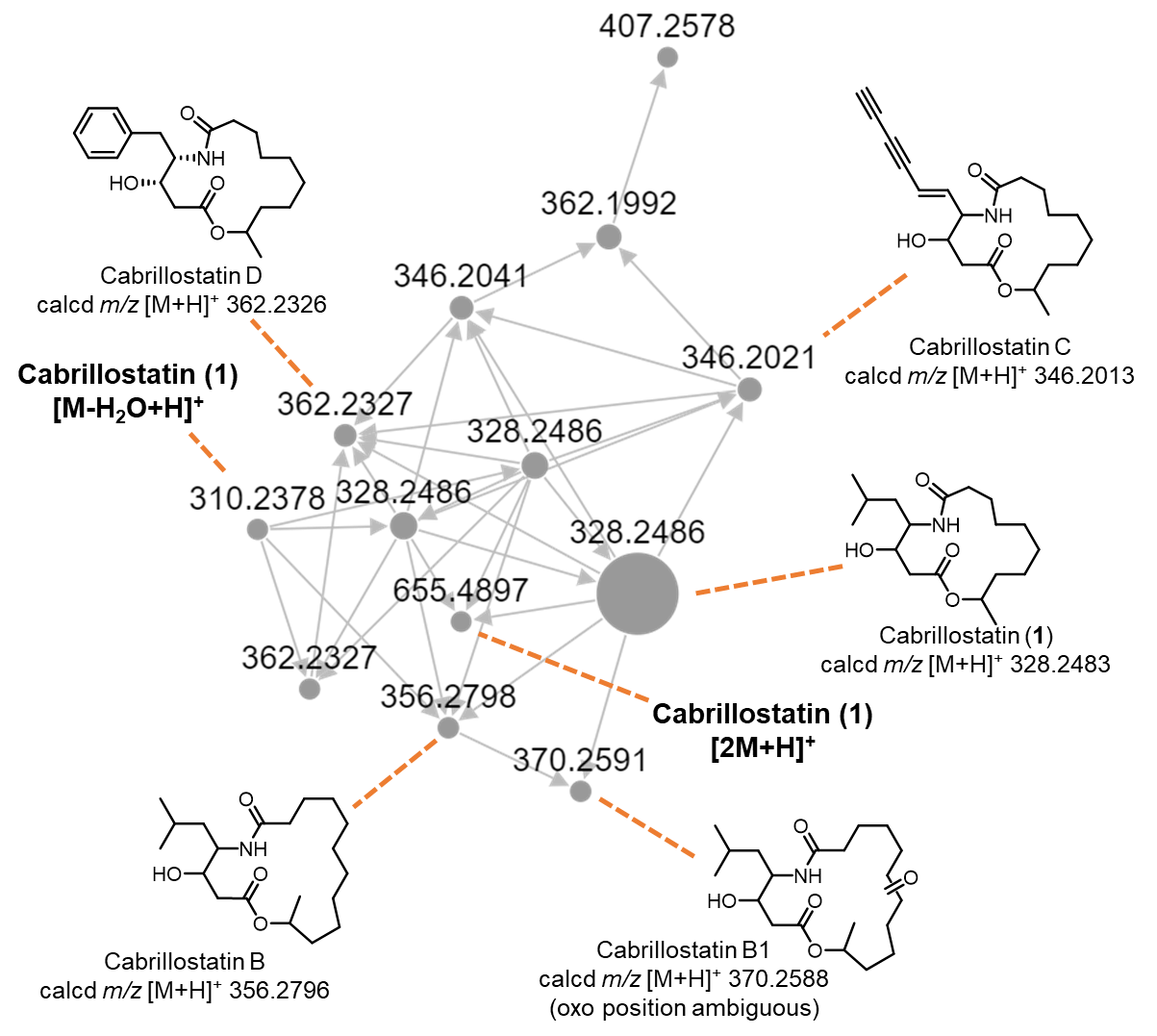

Fig. S29. Cabrillostatin (1) was detected in a molecular network constructed from an LCMS study of DOM in water samples collected at Imperial Beach, the Tijuana River estuary, and the Scripps Pier *(39)*. Nodes represent ions and are connected with edges (arrows) based on similarities in MS/MS fragmentation spectra. The structures of new compounds (cabrillostatins B, B1, C, and D) are proposed based on manual analysis of the MS and MS/MS spectra and comparison with those of 1. Node size represents the sum precursor intensity and correlates with the relative abundance of the compounds.

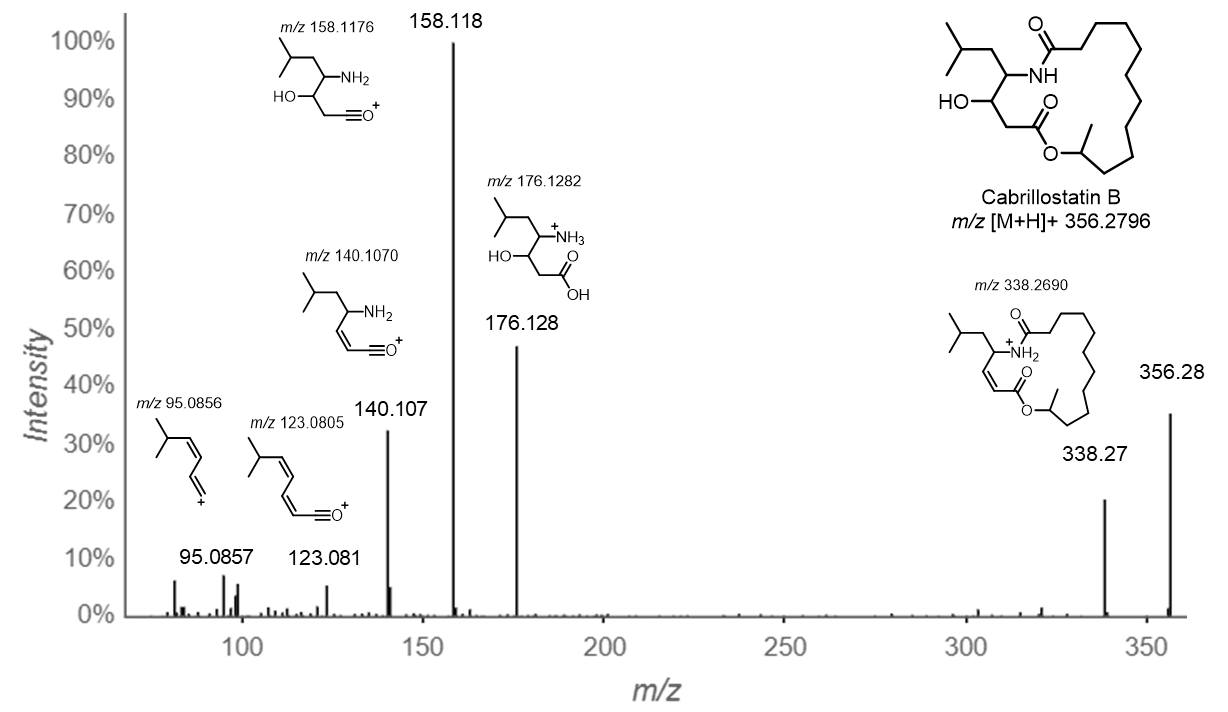

Fig. S30. Annotated MS/MS spectrum of cabrillostatin B.

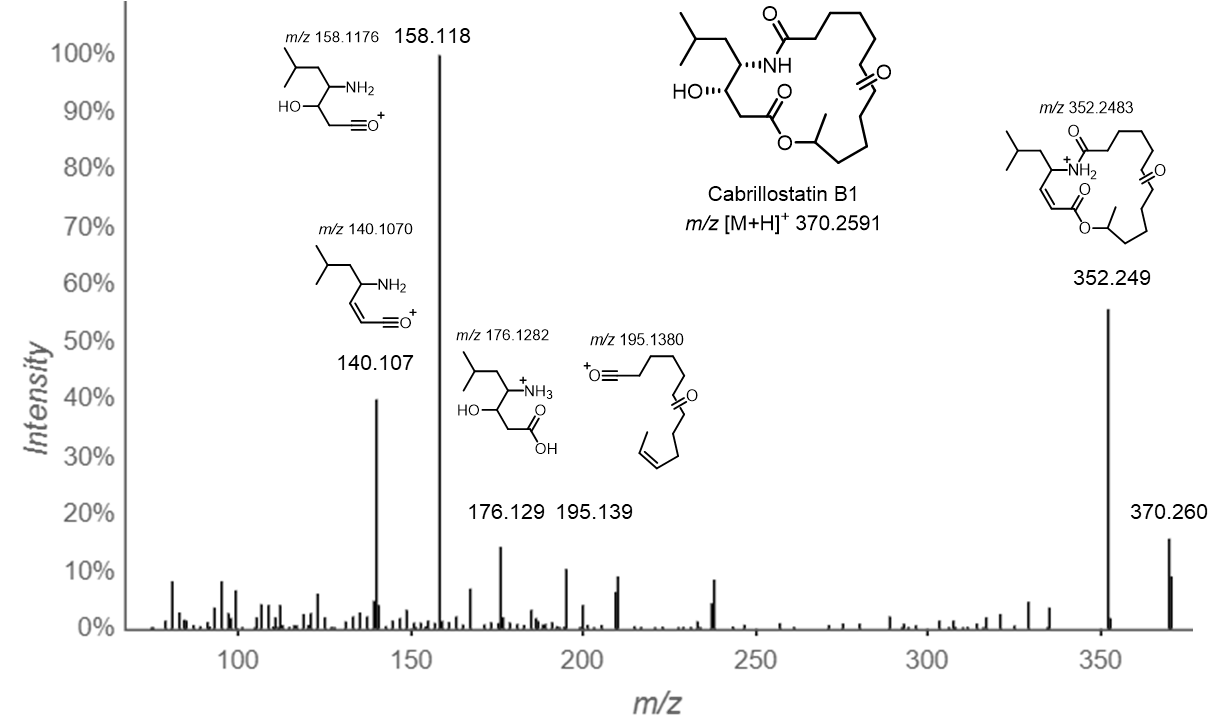

Fig. S31. Annotated MS/MS spectrum of cabrillostatin B1.

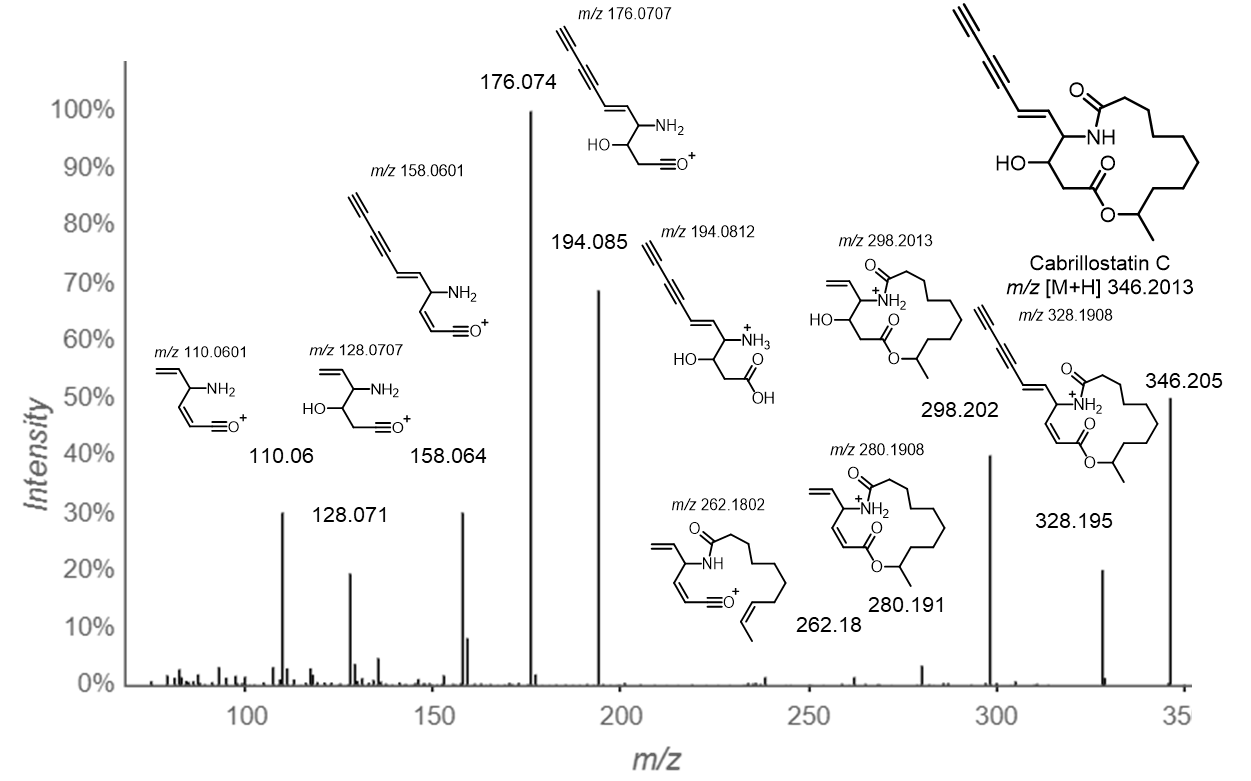

Fig. S32. Annotated MS/MS spectrum of cabrillostatin C.

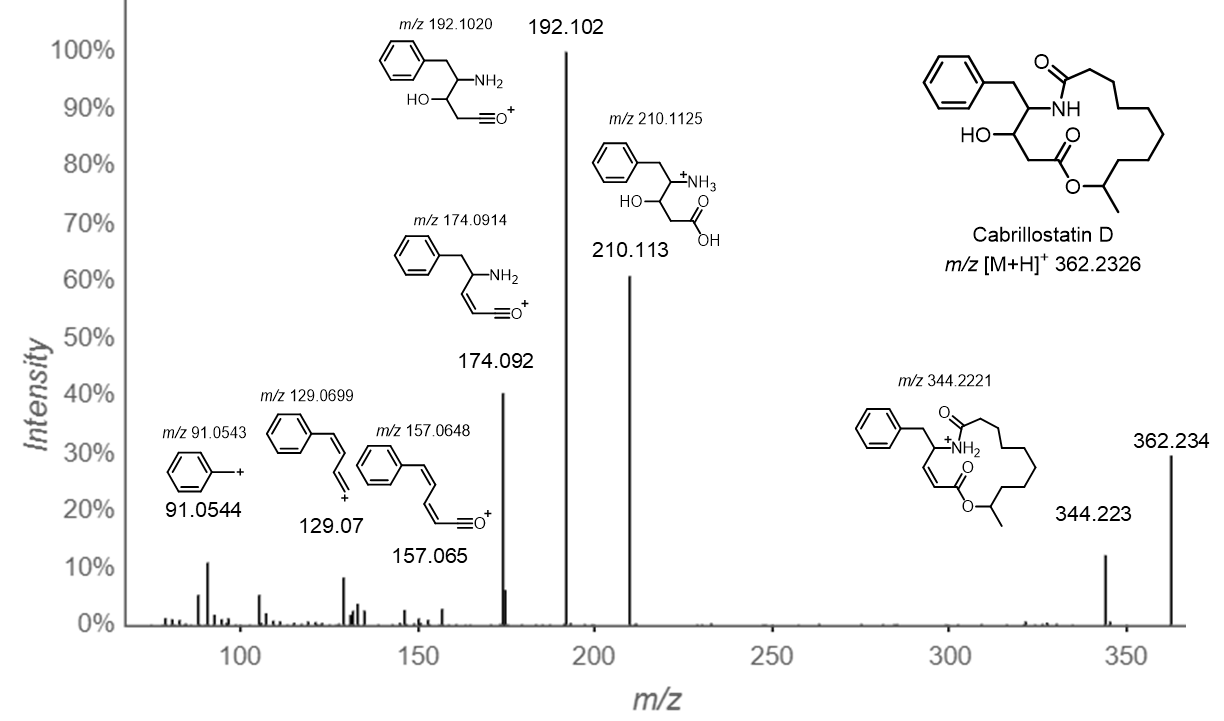

Fig. S33. Annotated MS/MS spectrum of cabrillostatin D.

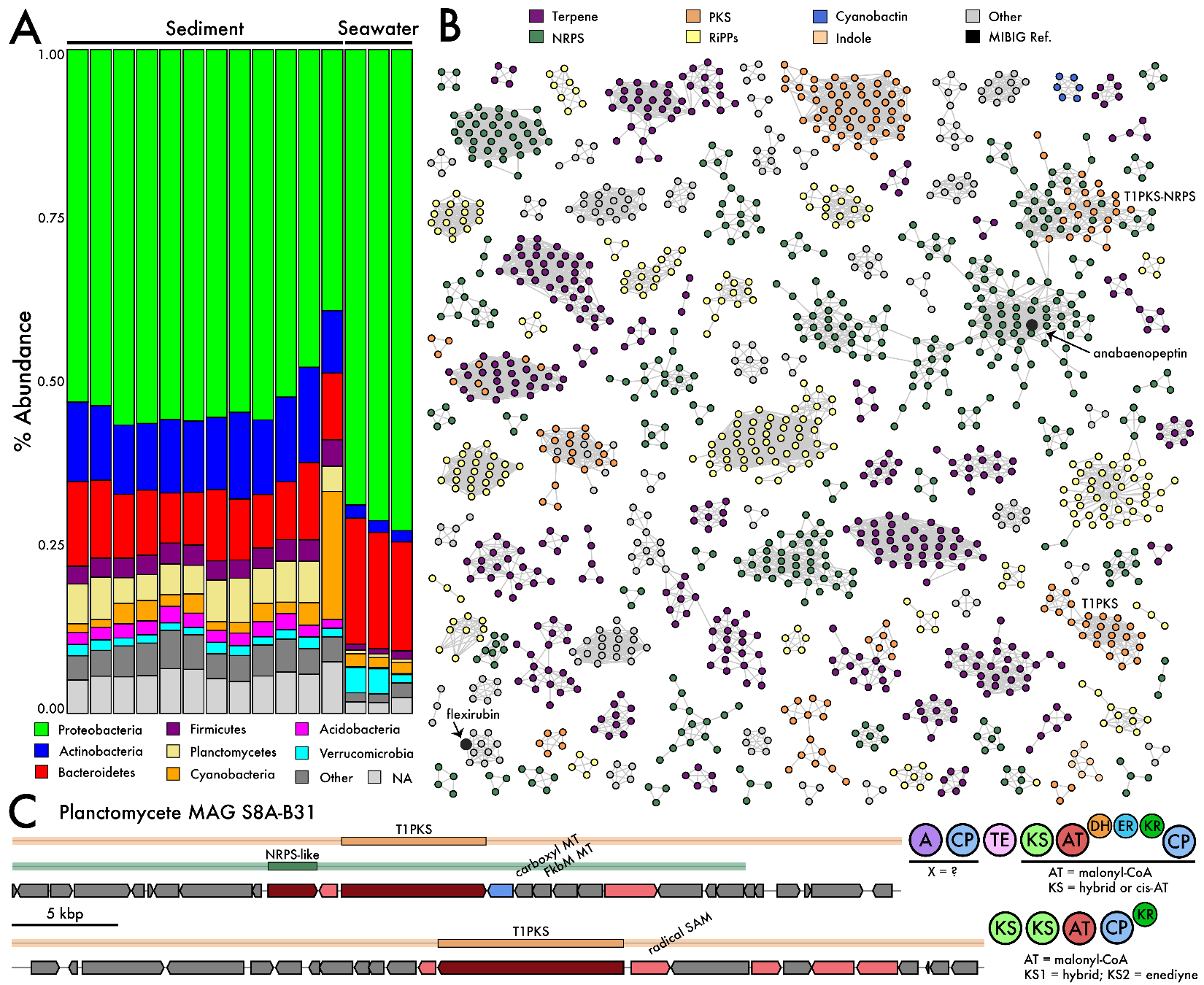

Fig. S34. CSMR microbial community composition and biosynthetic potential. A. Relative abundances of the eight most abundant bacterial phyla across sediment and seawater samples. B. Biosynthetic potential of bacterial communities clustered into gene cluster families (GCFs). Each node represents a predicted biosynthetic gene cluster (BGC) colored by its predicted product (reference MIBiG nodes in black). Singleton and doubleton GCFs not shown. C. Planctomycete MAG encodes two type 1 polyketide synthetase BGCs (PKS domains shown) that are candidates for the biosynthesis of compounds 2-3.

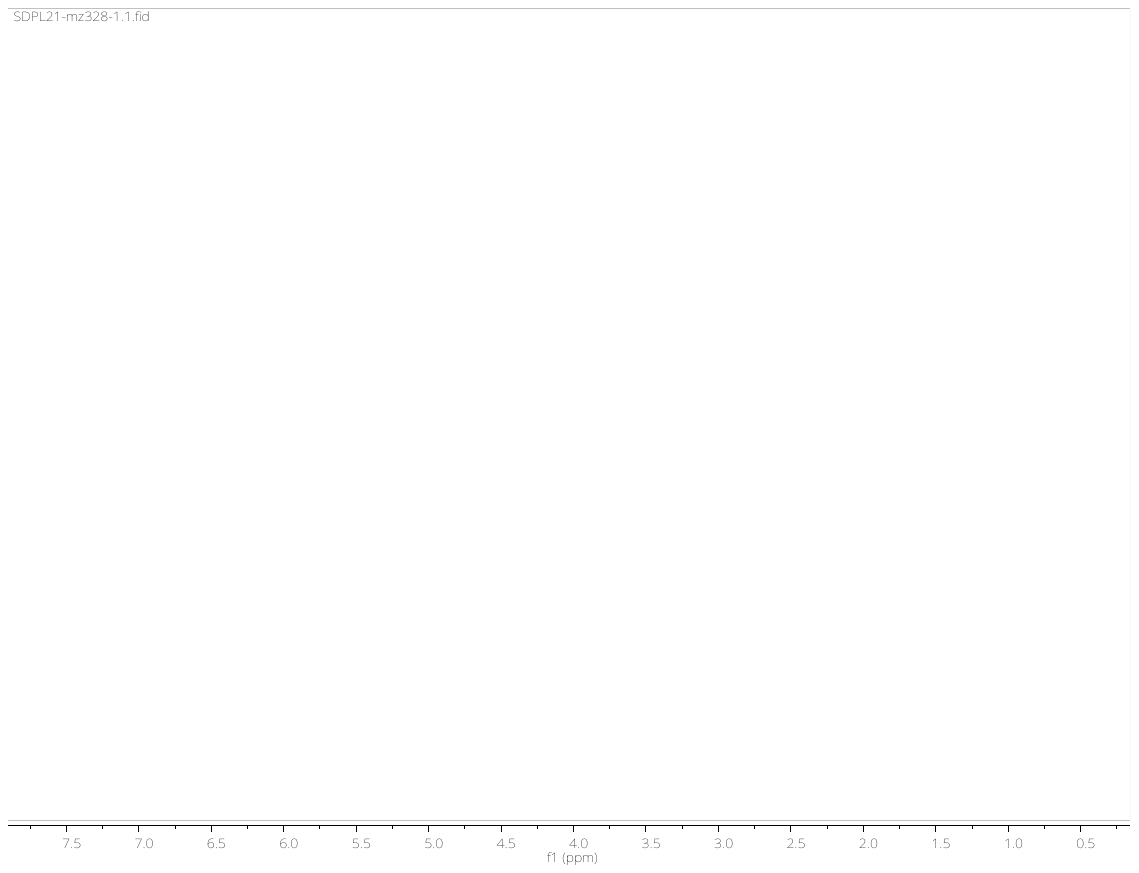

Fig. S35. ^1^H NMR spectrum of cabrillostatin (1) in CD_3_OD.

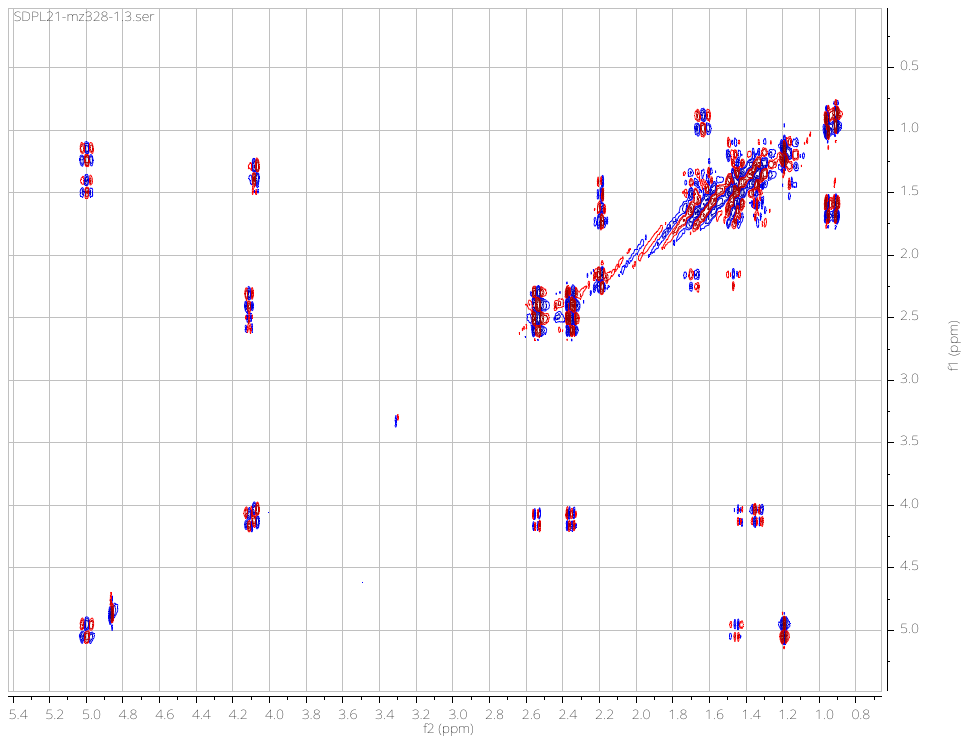

Fig. S36. DFT-COSY spectrum of 1 in CD_3_OD.

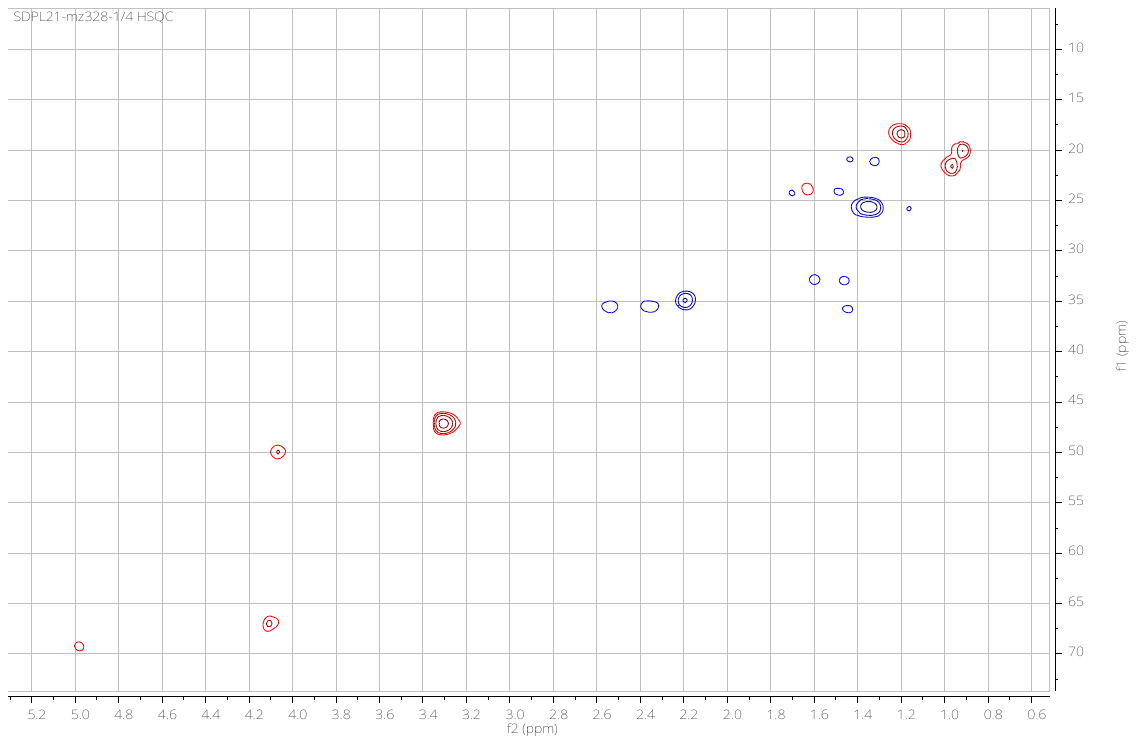

Fig. S37. HSQC spectrum of 1 CD_3_OD.

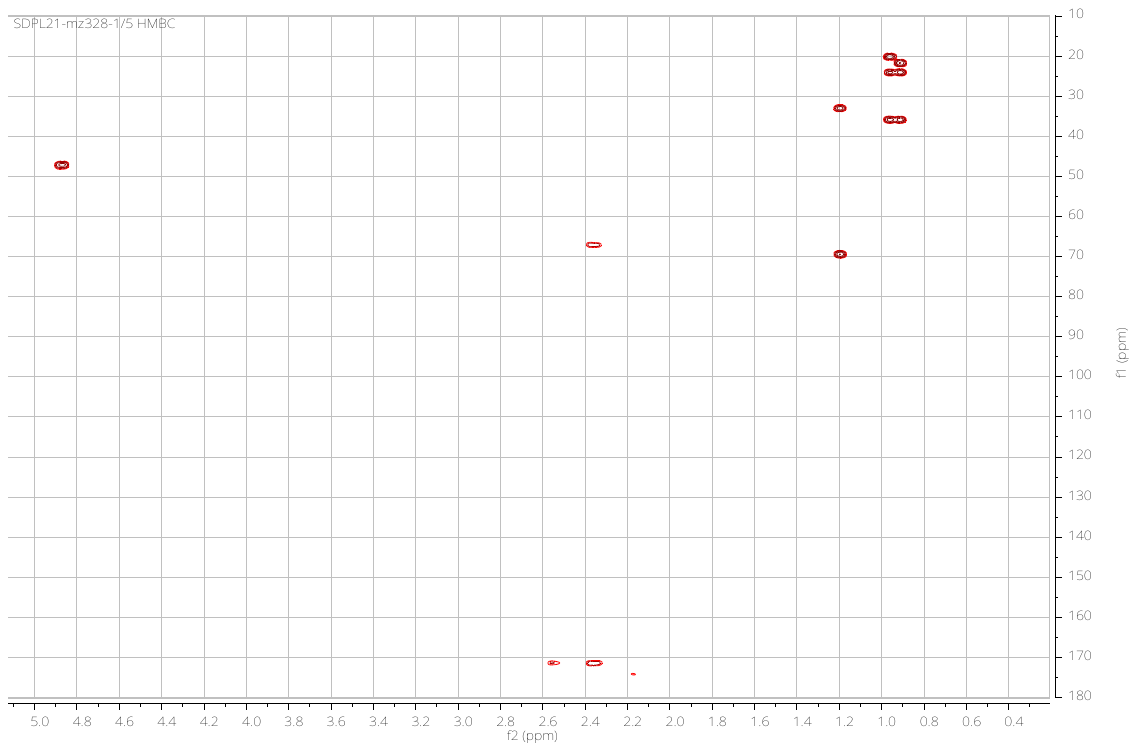

Fig. S38. HMBC spectrum of cabrillostatin (1) in CD_3_OD.

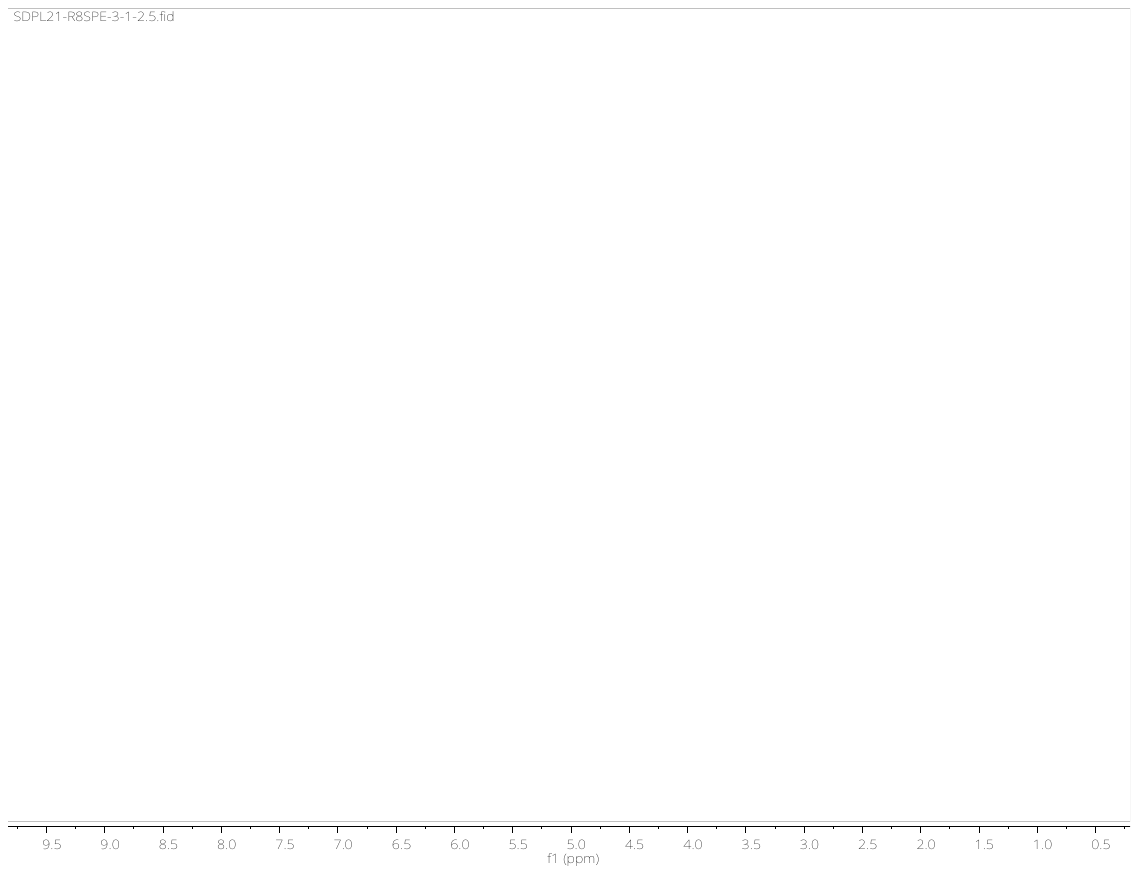

Fig. S39. ^1^H NMR spectrum of cabrillospiral A (2) in CD_3_CN.

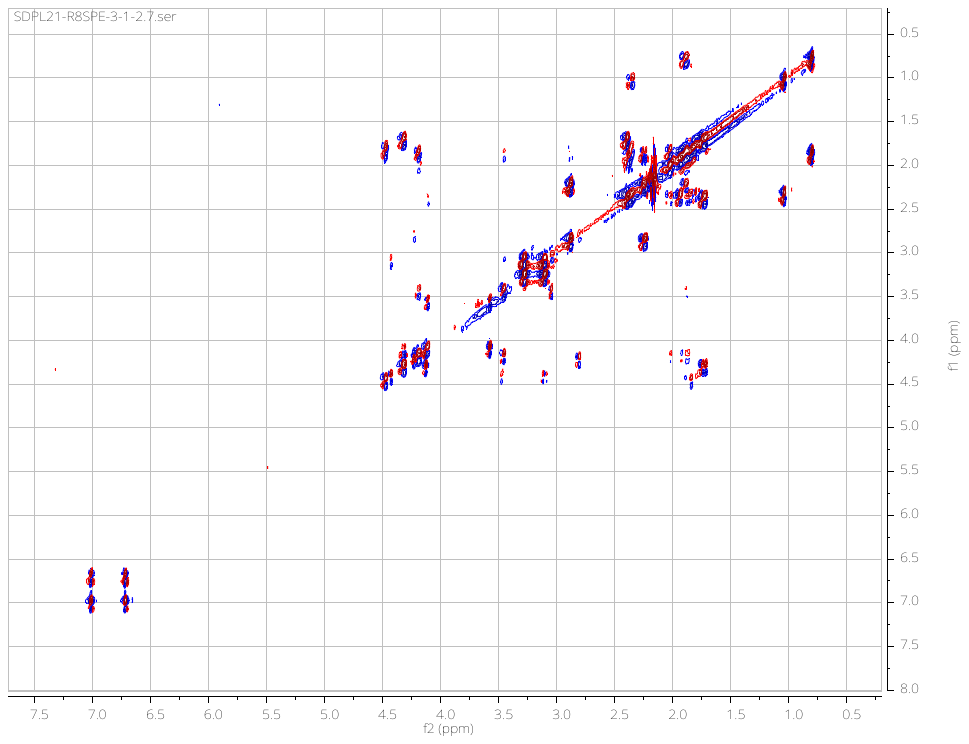

Fig. S40. COSY spectrum of 2 in CD_3_CN.

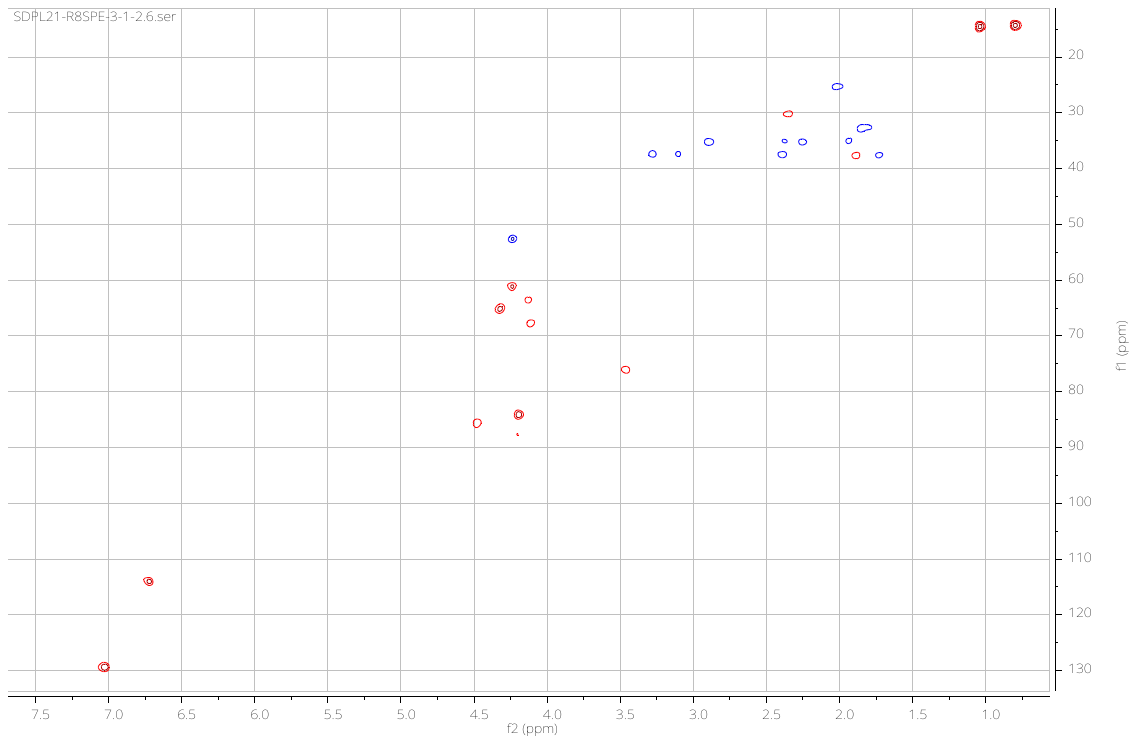

Fig. S41. HSQC spectrum of 2 in CD_3_CN.

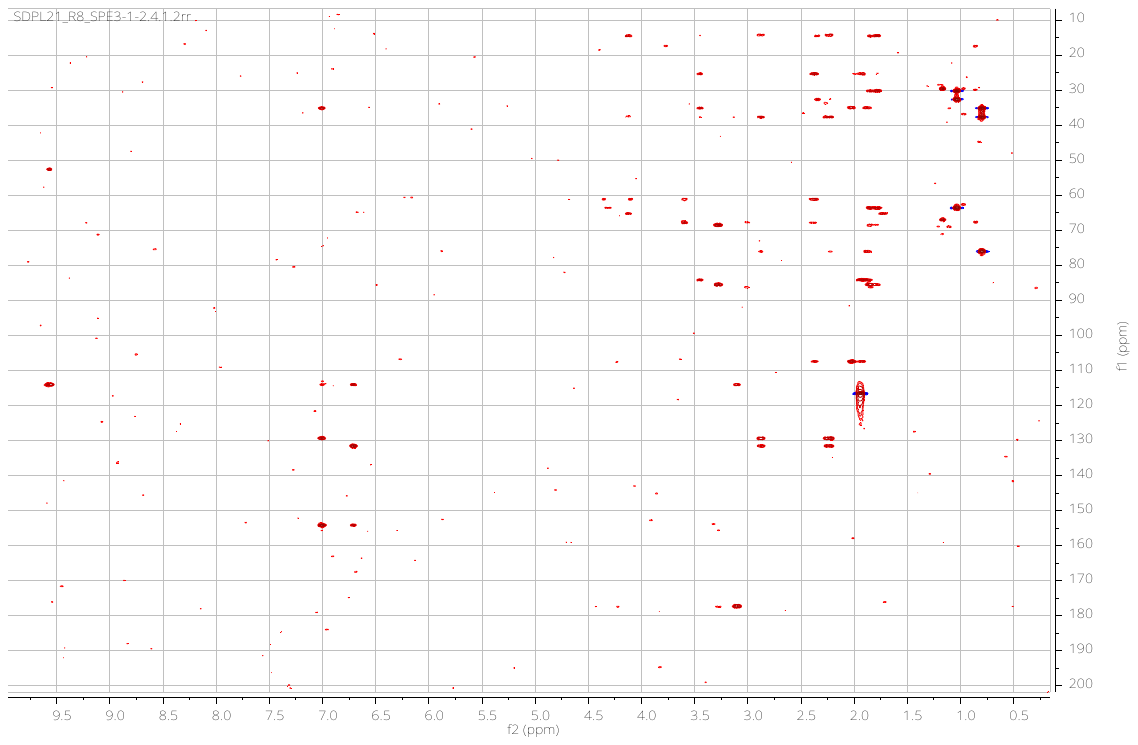

Fig. S42. HMBC spectrum of 2 in CD_3_CN.

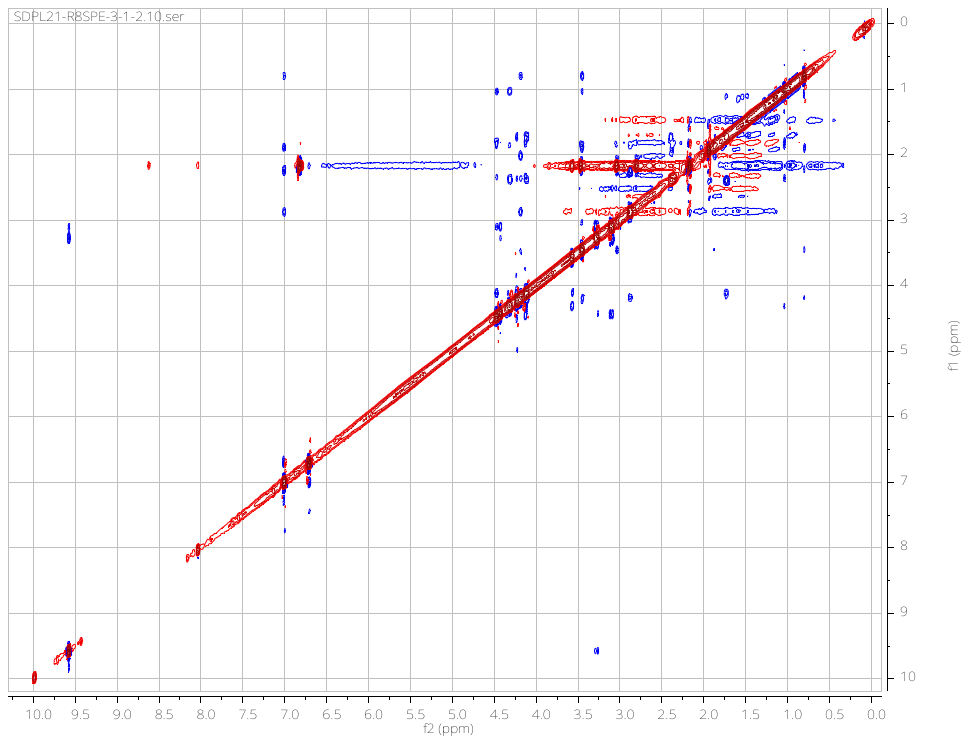

Fig. S43. NOESY spectrum of 2 in CD_3_CN.

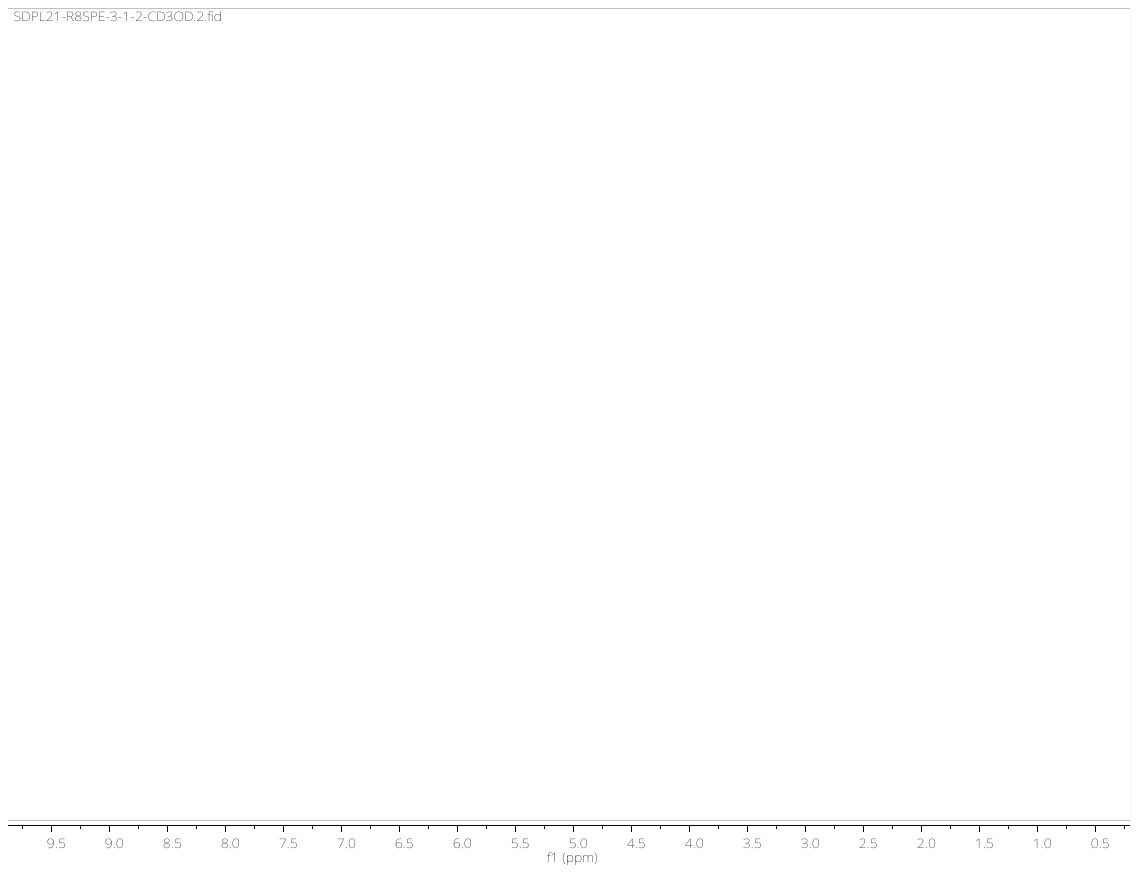

Fig. S44. ^1^H NMR spectrum of 2 in CD_3_OD.

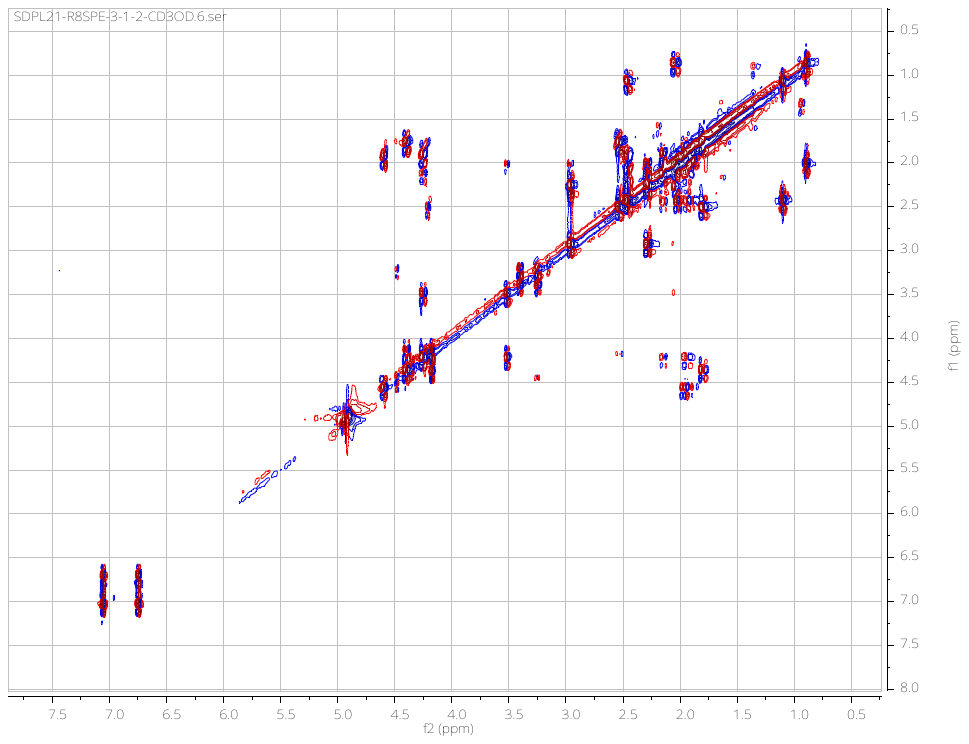

Fig. S45. COSY spectrum of 2 in CD_3_OD.

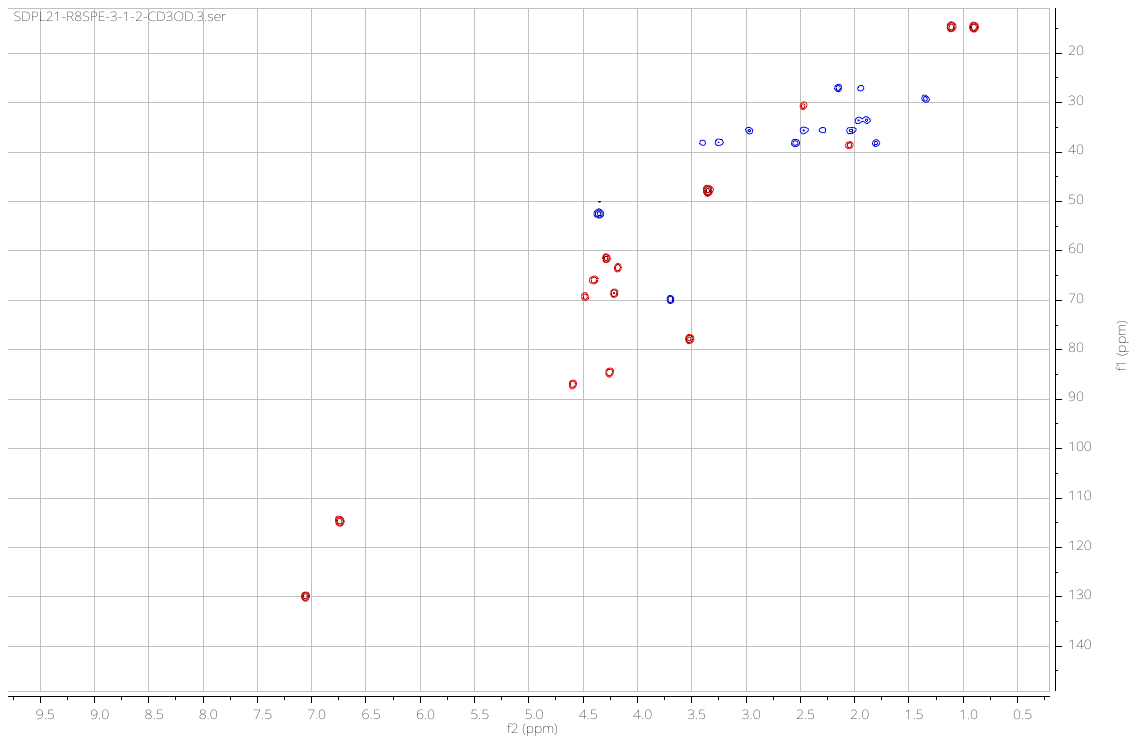

Fig. S46. HSQC spectrum of 2 in CD_3_OD.

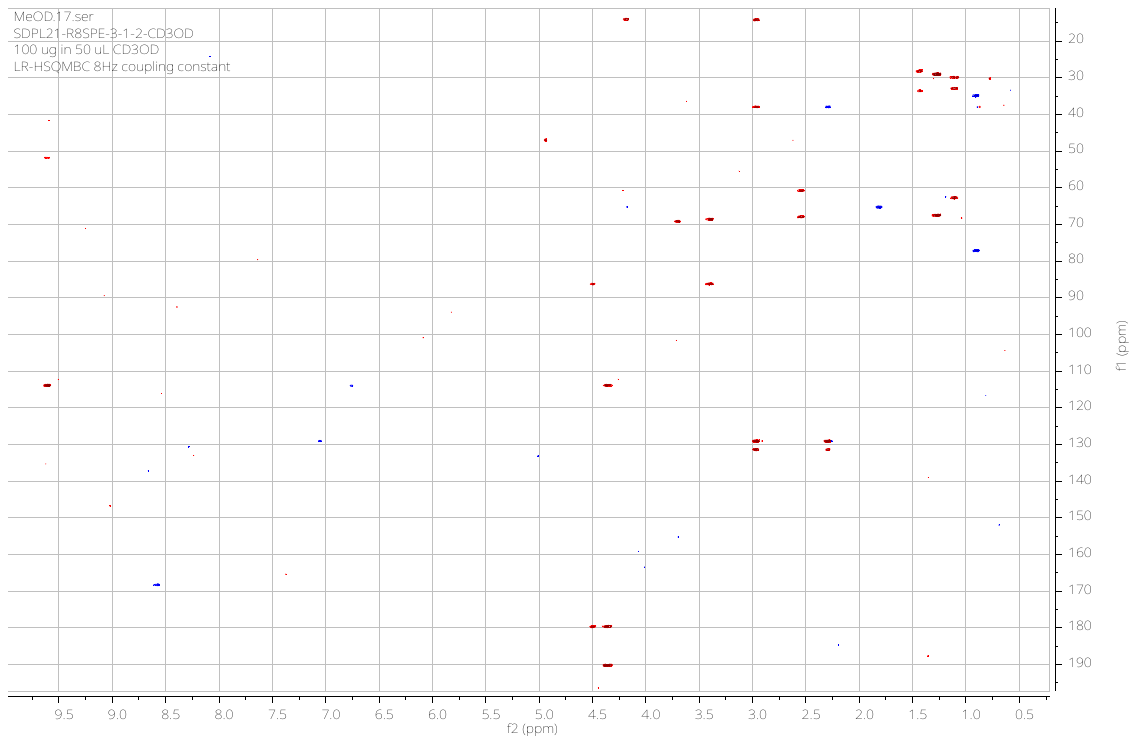

Fig. S47. LR-HSQMBC spectrum of 2 in CD_3_OD.

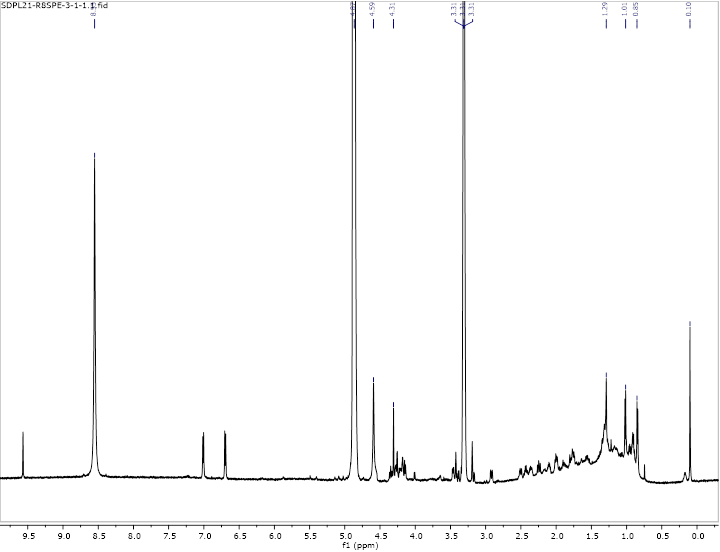

Fig. S48. ^1^H NMR spectrum of cabrillospiral B (3) in CD_3_OD.

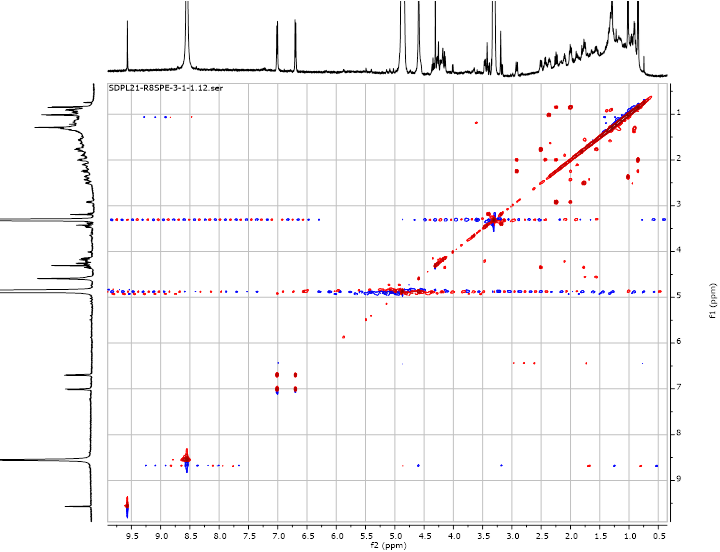

Fig. S49. COSY spectrum of 3 in CD_3_OD.

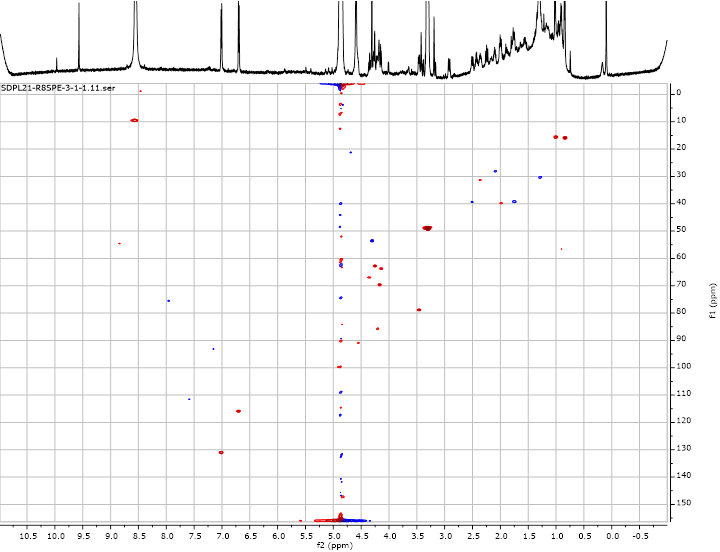

Fig. S50. HSQC spectrum of 3 in CD_3_OD.

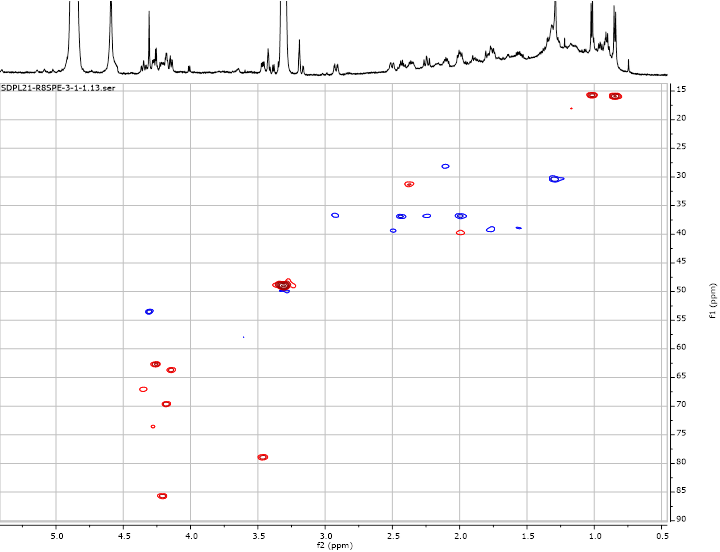

Fig. S51. HSQC spectrum of 3 in CD_3_OD (upfield region).

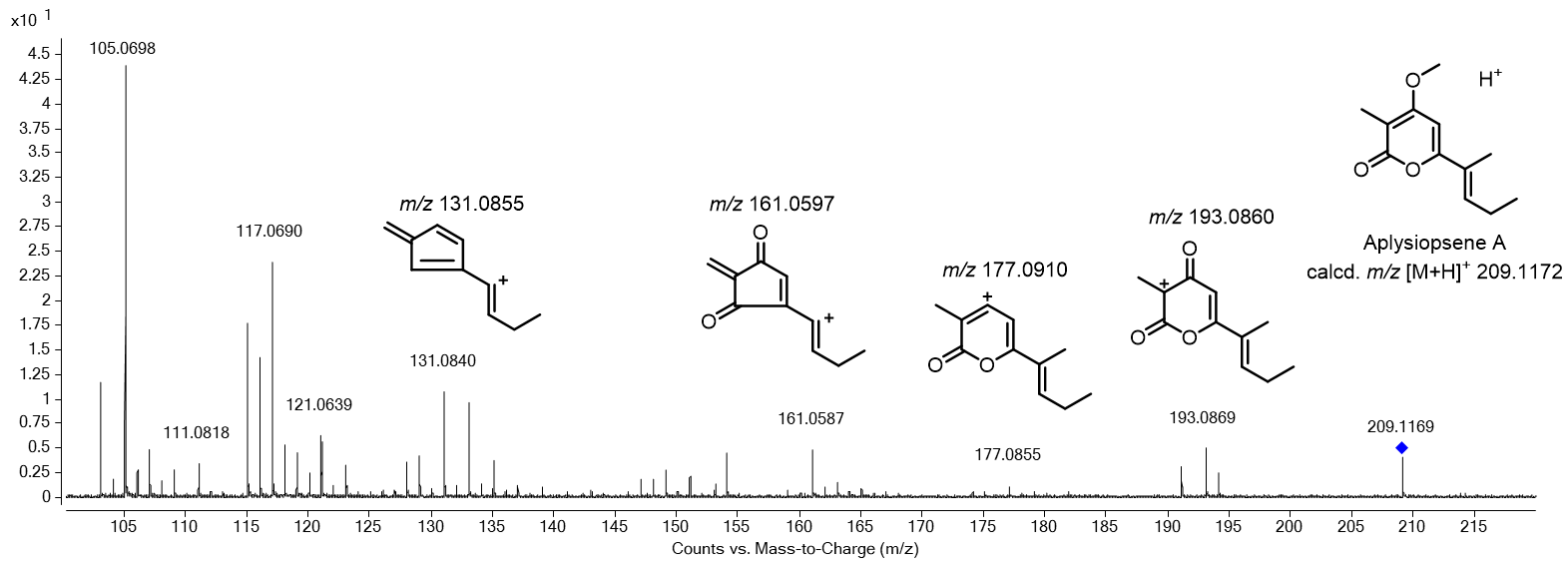

Fig. S52. Partially annotated ESI-MS/MS spectrum of aplysiopsene A.

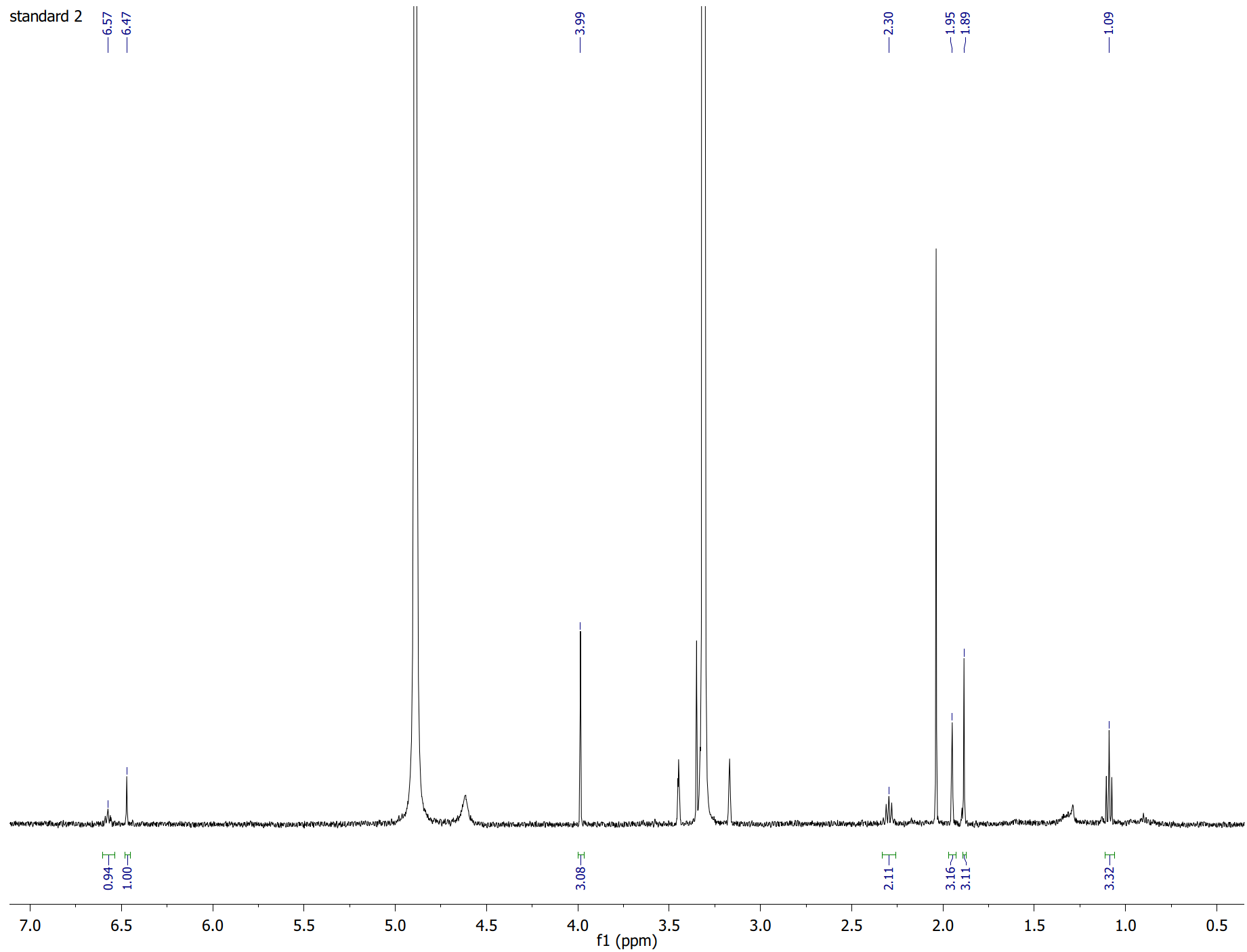

Fig. S53. ^1^H NMR of aplysiopsene A in CD_3_OD (500 MHz).

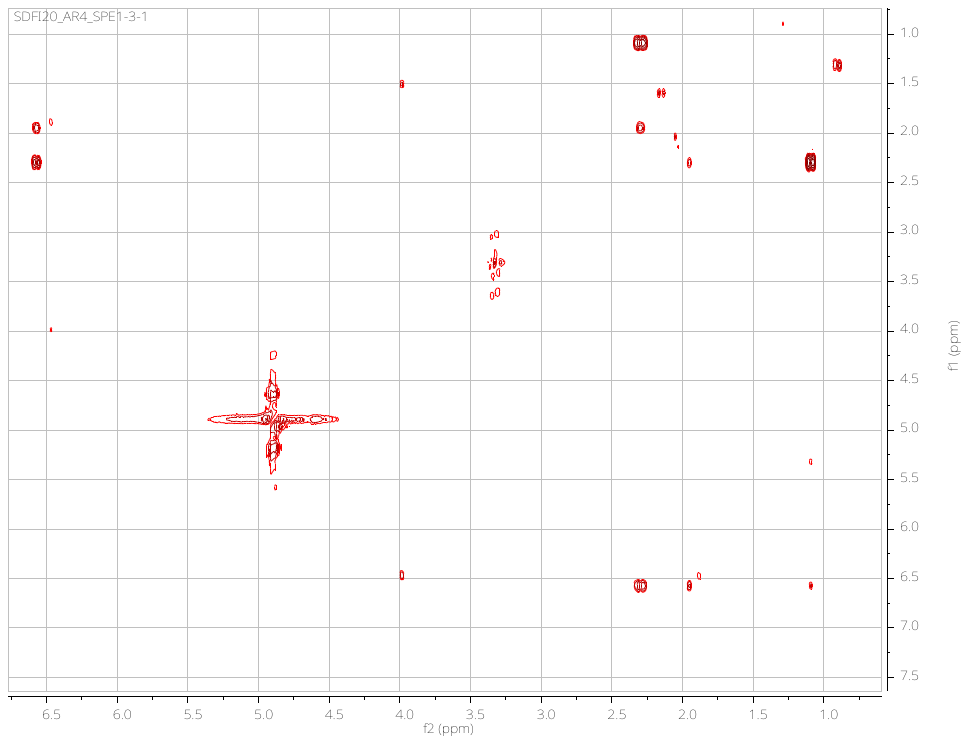

Fig. S54. COSY spectrum of aplysiopsene A in CD_3_OD (500 MHz).

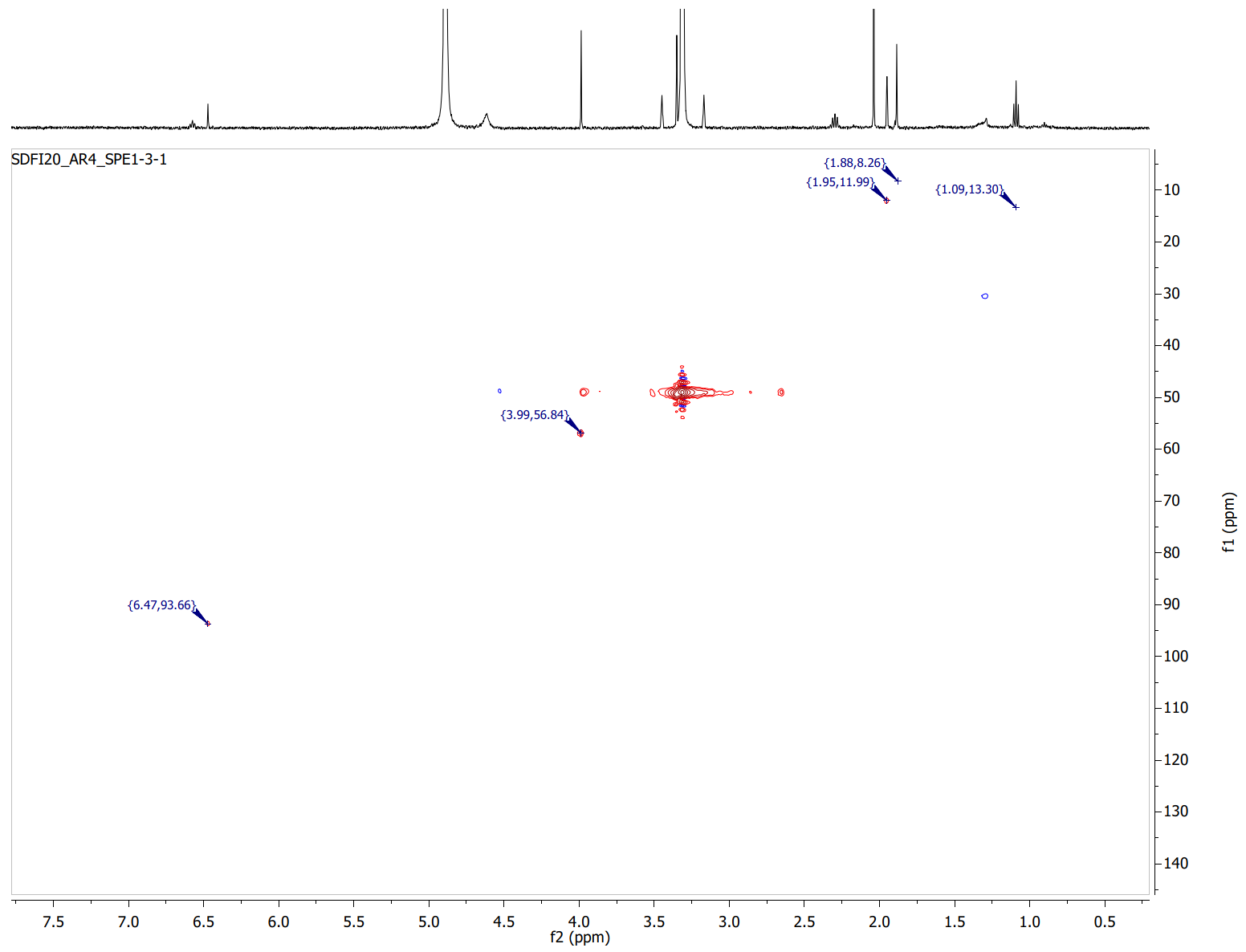

Fig. S55. HSQC spectrum of aplysiopsene A in CD_3_OD (500 MHz).

Table S1. NMR data for cabrillostatin (1) in CD_3_OD.

| 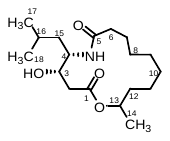 | | | | | |
| --- | --- | --- | --- | --- | --- |
| *^a^* | δ_C_ *^b^* | | δ_H_ (*J* in Hz) | COSY | HMBC |
| 1 | 173.3, C |  | |  |  |
| 2 | 37.5, CH_2_ | a 2.36, dd (7.7, 17.0)  b 2.54, dd (4.5, 17.0) | | H-3  H-3 | C-3, C-1  C-3 (w), C-1 |
| 3 | 68.9, CH | 4.11, ddd (3.6, 4.5, 7.7) | | H_2_-2, H-4 |  |
| 4 | 51.8, CH | 4.08, dt (3.6, 10.4) | | H-3, H_2_-15 |  |
| 5 | 175.7, C |  | |  |  |
| 6 | 36.8, CH_2_ | a 2.18, m  b 2.20, m | | H_2_-7 | C-5 |
| 7 | 26.1, CH_2_ | a 1.48, m  b 1.71, m | | H_2_-6, H_2_-8 |  |
| 8 | 27.5, CH_2_ | ab 1.35, m | | H_2_-7, |  |
| 9 | 27.5, CH_2_ | ab 1.35, m | | H-10b |  |
| 10 | 27.7, CH_2_ | a 1.16, m  b 1.35, m | | H_2_-9 |  |
| 11 | 22.9, CH_2_ | a 1.32, m  b 1.44, m | | H_2_-10  H_2_-10 |  |
| 12 | 34.9, CH_2_ | a 1.46, m  b 1.60, m | | H-13 |  |
| 13 | 71.2, CH | 5.00, m | | H-12b, H_3_-14 |  |
|  | 20.4, CH_3_ | 1.20, d (6.3) | | H-13 | C-12, C-13 |
| 15 | 37.8, CH_2_ | a 1.35, m  b 1.45, m | | H-4, H-16  H-4, H-16 |  |
| 16 | 25.8, CH | 1.64, m | | H_3_-17, H_3_-18 |  |
| 17 | 23.6, CH_3_ | 0.96, d (6.7) | | H-16 | C-15, C-16, C-18 |
| 18 | 22.0, CH_3_ | 0.90, d (6.5) | | H-16 | C-15, C-16, C-17 |

*^a^* All assignments are based on extensive 1D and 2D NMR measurements (COSY, HSQC, HMBC).

*^b^* Multiplicities determined by HSQC.

Table S2. NMR data of cabrillospiral A (2) in CD_3_CN.

| 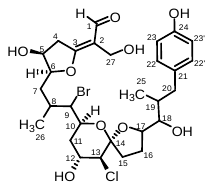 | | | | | | | |
| --- | --- | --- | --- | --- | --- | --- | --- |
| *^a^* | δ_C_ *^b^* | δ_H_ (*J* in Hz) | | COSY | | HMBC | NOESY |
| 1 | n. o., CHO | 9.58, s |  | | C-2, C-27 | | H_2_-4 |
| 2 | 116.2, C |  |  | |  | |  |
| 3 | 179.4, C |  |  | |  | |  |
| 4 | 39.6, CH_2_ | a 3.10, dd (4.9, 17.5)  b 3.29, d (17.5) | H-5 | | C-2, C-3  C-3, C-5, C-6 | | H-5, H-6  H-5 |
| 5 | 70.5, CH | 4.44, br s | H-4b | | C-3 (weak) | | H-4a, H-4b (weak), H-7a |
| 6 | 87.7, CH | 4.48, m | H-7a | |  | | H-8, H-9, H-4b, H_3_-26 |
| 7 | 32.7, CH_2_ | a 1.79, m  b 1.85, m | H-6 | | C-5, C-6, C-8,  C-9  C-5, C-6, C-8, C-9 | | H-6, H-9 |
| 8 | 30.2, CH | 2.35, m | H-9, H_3_-26 | | C-7, C-9, C-10 | |  |
| 9 | 65.7, CH | 4.13, m | H-10, | | C-10, C-11, C-26 | | H-6 |
| 10 | 67.3, CH | 4.32 br t (10.3) | H-9, H-11b | | C-9 (weak) | | H-11b, H_3_-26 |
| 11 | 39.7, CH_2_ | a 1.73, br t (12.9)  b 2.39, br t (12.9) | H-10  H-12 | | C-12 | | H-12, H-13 |
| 12 | 69.8, CH | 4.11, m | H-11a | | C-13 | |  |
| 12 | OH | 3.57 br d (7.5 Hz) | H-12 | |  | | H-10, H-12 |
| 13 | 63.2, CH | 4.23 m |  | | C-14 (weak) | | H-11a, H-15b |
| 14 | 109.6, C |  |  | |  | |  |
| 15 | 37.1, CH_2_ | a 1.92, m  b 2.37, m | H_2_-16 | | C-14, C-16, C-17  C-13, C-14, C-16 | |  |
| 16 | 27.3, CH_2_ | a 1.88, m  b 2.02, m | H_2_-15 | | C-15, C-17, C-18  C-14, C-15 | |  |
| 17 | 86.2, CH | 4.20, m | H_2_-16, H-18 | |  | | H-15b, H-16b |
| 18 | 78.3, CH | 3.46 m | H-17, H-19 | | C-16, C-17, C-19, C-20 | | H-16a, H_3_-25, H_3_-26 (transannular) |
| 19 | 39.8, CH | 1.89, m | H-17, H_2_-20, H_3_-25 | | C-20 | |  |
| 20 | 37.3, CH_2_ | a 2.25 m  b 2.89 br d (13.0) | H-19  H-19 | | C-21, C-22/22′, C-19, C-25 | |  |
| 21 | 133.7, C |  |  | |  | |  |
| 22, 22′ | 131.4, CH | 7.01, d (8.0) | H-23/23’ | | C-20, C-23/23′, C-24 | |  |
| 23, 23′ | 116.1, CH | 6.72 d (8.0) | H-22/22’ | | C-21, C-22/22′, C-24 | |  |
| 24 | 154.2 C |  |  | |  | |  |
| 25 | 16.4, CH_3_ | 0.80, d (6.8) | H-19 | | C-18, C-19, C-20 | |  |
| 26 | 16.6, CH_3_ | 1.04, d (6.2) | H-8 | | C-7, C-8, C-9 | |  |
| 27 | 54.7, CH_2_ | 4.24, br s |  | | C-3 (weak) | |  |

*^a^* All assignments are based on extensive 1D and 2D NMR measurements (COSY, HSQC, HMBC).

*^b^* Multiplicities determined by HSQC.

Table S3. NMR data of cabrillospiral A (2) in CD_3_OD.

| 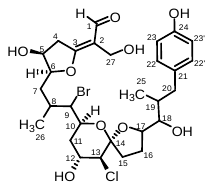 | | | | |
| --- | --- | --- | --- | --- |
| *^a^* | δ_C_ *^b^* | δ_H_ (*J* in Hz) | COSY | HMBC |
| 1 | 192.1, C | 9.57, s |  | C-2, C-27 |
| 2 | 115.8, C |  |  |  |
| 3 | 181.5, C |  |  |  |
| 4 | 39.3, CH_2_ | a 3.21, dd (4.9, 17.6)  b 3.36, d (17.6) | H-5 | C-5, C-6 |
| 5 | 70.5, CH | 4.43, dd (3.5, 4.9) | H-4b, H-6 | C-3, C-6 |
| 6 | 88.2, CH | 4.55, ddd (3.5, 4.5, 8.5) | H-5, H_2_-7 |  |
| 7 | 34.9, CH_2_ | a 1.85, m  b 1.92, m | H-6, H-8  H-6, H-8 | C-9 |
| 8 | 31.8, CH | 2.43, m | H-7, H-9, H-26 |  |
| 9 | 64.6, CH | 4.14, dd (2.2, 8.8) | H-8, H-10 | C-10, C-11, C-26 |
| 10 | 67.2, CH | 4.36 ddd (1.9, 8.8, 11.0) | H-9, H-11b |  |
| 11 | 39.4, CH_2_ | a 1.76, ddd (2.5, 11.0, 14.0)  b 2.50, ddd (1.9, 3.6, 14.0) | H-10, H-12  H-12 | C-10  C-12, C-13 |
| 12 | 69.8, CH | 4.17, ddd (2.5, 3.2, 3.6) | H_2_-11 | C-10, C-13, C-14 (weak) |
| 13 | 62.8, CH | 4.24, d (3.2) |  |  |
| 14 | 109.3, C |  |  |  |
| 15 | 36.9, CH_2_ | a 1.99, m  b 2.43, m | H_2_-16  H_2_-16 |  |
| 16 | 28.3, CH_2_ | a 1.90, m  b 2.11, m | H-15a, H-17  H-15a, H-17 |  |
| 17 | 85.8, CH | 4.21, m | H_2_-16, H-18 | C-16 |
| 18 | 79.0, CH | 3.48, dd (4.0, 8.0) | H-17, H-18 | C-16, C-17, C-20 |
| 19 | 39.9, CH | 2.00, m | H-18, H_2_-20, H_3_-25 | C-20 |
| 20 | 36.9, CH_2_ | a 2.24 dd (10.6, 13.3)  b 2.93 dd (3.5, 13.3) | H-19  H-19 | C-19, C-21, C-22/22’, C-25  C-19, C-21, C-22/22’, C-25 |
| 21 | 133.4, C |  |  |  |
| 22, 22′ | 131.1, CH | 7.02, d (8.4) | H-23/23′ | C-21, C-22/22’, C-24 |
| 23, 23′ | 115.9, CH | 6.70 d (8.4) | H-22/22′ | C-23/23′, C-24 |
| 24 | 156.2, C |  |  |  |
| 25 | 16.0, CH_3_ | 0.86, d (6.7) |  | C-18, C-19, C-20 |
| 26 | 15.9, CH_3_ | 1.07, d (6.5) |  | C-7, C-8, C-9 |
| 27 | 53.7, CH_2_ | a 4.30, d (11.3)  b 4.33, d (11.3) |  | C-1, C-2, C-3  C-1, C-2, C-3 |

*^a^* All assignments are based on extensive 1D and 2D NMR measurements (COSY, HSQC, HMBC).

*^b^* Multiplicities determined by HSQC.

Table S4. Comparison of ^1^H and ^13^C NMR data for cabrillospirals A (2) and B (3) in CD_3_OD.

|  | 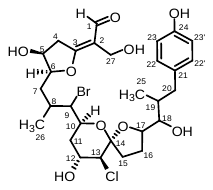 | | 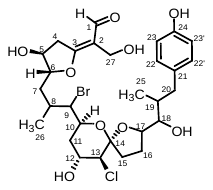 | |
| --- | --- | --- | --- | --- |
|  | Cabrillospiral A (**2**) | | Cabrillospiral B (**2**) | |
| *^a^* | δ_C_ *^b^* | δ_H_ (*J* in Hz) | δ_C_ *^b^* | δ_H_ (*J* in Hz) |
| 1 | 192.1, C | 9.57, s | n.o. | 9.57, s |
| 2 | 115.8, C |  | n.o. |  |
| 3 | 181.5, C |  | n.o. |  |
| 4 | 39.3, CH_2_ | a 3.21, dd (4.9, 17.6)  b 3.36, d (17.6) | n.o. | n.o. |
| 5 | 70.5, CH | 4.43, dd (3.5, 4.9) | 73.6, CH | 4.28, dd (2.9, 5.9) |
| 6 | 88.2, CH | 4.55, ddd (3.5, 4.5, 8.5) | 90.9, CH | 4.55, m |
| 7 | 34.9, CH_2_ | a 1.85, m  b 1.92, m | 38.9, CH_2_ | a 1.55, m  b 1.76, m |
| 8 | 31.8, CH | 2.43, m | 31.4, CH | 2.38, m |
| 9 | 64.6, CH | 4.14, dd (2.2, 8.8) | 63.7, CH | 4.14, dd (2.1, 9.0) |
| 10 | 67.2, CH | 4.36 ddd (1.9, 8.8, 11.0) | 67.1, CH | 4.35, ddd (1.7, 9.0, 11.0) |
| 11 | 39.4, CH_2_ | a 1.76, ddd (2.5, 11.0, 14.0)  b 2.50, ddd (1.9, 3.6, 14.0) | 39.4, CH_2_ | a 1.78, m  b 2.50, br d (13.5) |
| 12 | 69.8, CH | 4.17, ddd (2.5, 3.2, 3.6) | 69.7, CH | 4.18, dd (3.2, 6.5) |
| 13 | 62.8, CH | 4.24, d (3.2) | 62.7, CH | 4.26, d (3.2) |
| 14 | 109.3, C |  | n.o. |  |
| 15 | 36.9, CH_2_ | a 1.99, m  b 2.43, m | 36.9, CH_2_ | a 1.99, m  b 2.43, m |
| 16 | 28.3, CH_2_ | a 1.90, m  b 2.11, m | 28.3, CH_2_ | a 1.89, m  b 2.11, m |
| 17 | 85.8, CH | 4.21, m | 85.7, CH | 4.21, m |
| 18 | 79.0, CH | 3.48, dd (4.0, 8.0) | 79.0, CH | 3.46, dd (4.0, 8.0) |
| 19 | 39.9, CH | 2.00, m | 39.8, CH | 2.00, m |
| 20 | 36.9, CH_2_ | a 2.24 dd (10.6, 13.3)  b 2.93 dd (3.5, 13.3) | 36.7, CH_2_ | a 2.24 dd (10.6, 13.3)  b 2.92 dd (3.5, 13.3) |
| 21 | 133.4, C |  | n.o. |  |
| 22, 22′ | 131.1, CH | 7.02, d (8.4) | 131.1, CH | 7.01, d (8.4) |
| 23, 23′ | 115.9, CH | 6.70 d (8.4) | 115.9, CH | 6.70 d (8.4) |
| 24 | 156.2, C |  | n.o. |  |
| 25 | 16.0, CH_3_ | 0.86, d (6.7) | 15.9, CH_3_ | 0.86, d (7.0) |
| 26 | 15.9, CH_3_ | 1.07, d (6.5) | 15.7, CH_3_ | 1.02, d (6.6) |
| 27 | 53.7, CH_2_ | a 4.30, d (11.3)  b 4.33, d (11.3) | 53.6, CH_2_ | ab 4.31, s |

*^a^* All assignments are based on extensive 1D and 2D NMR measurements (COSY, HSQC, HMBC).

*^b^* Multiplicities determined by HSQC.

Table S5. LCMS results and MASST database matches for compounds captured using SMIRC.

| Compound name or *m/z* | calcd. MF | Occurrence in SMIRC extracts | Number of MASST datasets hits, type of study |
| --- | --- | --- | --- |
| Cabrillostatin (**1**) | C_18_H_33_NO_4_ | all | 11, DOM CA, deep water samples <500 m |
| Cabrillospirals **2-3** | C_29_H_40_BrClO_9_ | 2021 | No hits |
| Aplysiopsene A | C_12_H_16_O_3_ | 2021-2022 | 2, Indonesian coral reef colonization experiments on ceramic plates, *Salinispora* sp. |
| 757.447 [M+H]^+^ | C_43_H_64_O_11_ | all | 22, DOM, CA and tropical Pacific Ocean |
| 727.437 [M+H]^+^ | C_42_H_62_O_10_ | all | 2, DOM, CA |
| 743.431 [M+H]^+^ | C_42_H_62_O_11_ | all | 1, DOM, CA |
| 769.448, [M+H]^+^ | C_44_H_64_O_11_ | all | 5, DOM, CA |
| 785.462, [M+H]^+^ | C_41_H_68_O_14_ | all | No hits |
| 1031.550, [M+H]^+^ | C_62_H_78_O_13_ | all | 1, DOM, CA |
| 1211.64, [M+Na]^+^ | C_76_H_90_O_13_ | all | No hits |
| 864.3641, [M+H]^+^ | C_42_H_64_Cl_3_NO_11_ | 2021 | No hits |
| 577.381, [M+H]^+^ | C_27_H_52_N_4_O_9_ | 2021-2022 | 7, DOM, CA |
| 592.0393, [M+H]^+^ | C_26_H_24_Cl_2_BrN_3_O_4_ | 2021 | No hits |
| 1086.6190, [M+NH_4_]^+^ | C_55_H_88_O_20_ | all | 6, DOM, CA, South African Tunicate |
| 456.1317 [M+Na]^+^ | C_22_H_25_Cl_2_N_3_O_2_ | all | 3, DOM, CA |
| 606.3572 [M+H]^+^ | C_34_H_52_ClNO_6_ | 2022 | No hits |
| 644.3936 [M+H]^+^ | C_34_H_58_ClNO_8_ | 2022 | 5, DOM CA, Coral reefs (Hawaii, Moorea) |
| 488.2057 [M+H]^+^ | C_23_H_34_ClNO_8_ | 2021 |  |
| 913.4561 [M+Na]^+^ | C_47_H_70_O_16_ | 2022 |  |

Table S6. Cancer cell lines used for the high content screening.

| Name | Cancer type | Mutation status | Reference compound classification accuracy |
| --- | --- | --- | --- |
| 786-O | Kidney | VHL, p53, PTEN, and p16 | 0.942 |
| A549 | Lung | KRAS | 0.974 |
| AsPC1 | Pancreas | KRAS | 0.938 |
| DU145 | Prostate | p53 | 0.963 |
| HCT116 | Colon | KRAS | 0.906667 |
| HEPG2 | Liver | p53 | 0.94 |
| OVCAR4 | Ovarian | TP53 | 0.9 |
| U87MG | Brain | PTEN | 0.966 |
| WM164 | Skin | BRAF | 0.945 |

Table S7. Reference compounds used for the high content screening.

| **Compound** | **Category** | **Targets** | **Pathway** | **Source well conc. in µM** |
| --- | --- | --- | --- | --- |
| Colchicine | MT | B-Tubulin | MT polymerization | 5000 |
| Docetaxel | MT | Microtubules | MT depolymerization | 5000 |
| Epothilone B | MT | Microtubules | MT depolymerization | 5000 |
| Nocodazole | MT | Î²-tubulin | MT polymerization | 5000 |
| Vinblastine | MT | Tubulin | MT polymerization | 5000 |
| AUY922 | HSP90 | HSP90Î±/Î² | HSP | 5000 |
| 17-AAG | HSP90 | Pan-HSP | HSP | 5000 |
| 17-DMAG | HSP90 | Pan-HSP | HSP | 5000 |
| Geldanamycin | HSP90 | HSP90 | HSP | 5000 |
| SNX2112 | HSP90 | HSP90Î±/Î² | HSP | 5000 |
| AZD8055 | mTOR | mTOR | mTOR | 5000 |
| Deforolimus | mTOR | mTOR | mTOR | 5000 |
| INK128 | mTOR | mTOR | mTOR | 5000 |
| Rapamycin | mTOR | mTOR | mTOR | 5000 |
| Torin 1 | mTOR | mTORC1, mTORC2 | mTOR | 1000 |
| Camptothecin | DNA | Topo I | Topoisomerases | 5000 |
| Eloxatin | DNA | DNA | DNA Synthesis | 25000 |
| Etoposide | DNA | Topo II | Topoisomerases | 5000 |
| Gemcitabine | DNA | Nucleic Acids | Nucleic Acid Synthesis | 5000 |
| Pemetrexed | DNA | DHFR, TS, GARFT | Nucleic Acid Synthesis | 5000 |
| Apicidin | HDAC | HDAC3 | Epigenetic | 5000 |
| Oxamflatin | HDAC | HDAC | Epigenetic | 5000 |
| Panobinostat | HDAC | HDAC | Epigenetic | 5000 |
| SAHA | HDAC | HDAC | Epigenetic | 5000 |
| Aclacinomycin | Proteasome | Topo I, Topo II, 20S proteaome | Topoisomerases, Protein degradation | 5000 |
| Bortezomib | Proteasome | 20S proteasome | Protein degradation | 5000 |
| MG-115 | Proteasome | 20S Proteasome, 26S Proteasome | Protein degradation | 5000 |
| MG132 | Proteasome | proteasome, calpain | Protein degradation | 5000 |

Table S8. Results of bioactivity testing for 1-3 in the high content screening with nine cancer cell lines.

| Compound | Bioactive  p value | Cell line | Compound | Bioactive p value | Cell Line |
| --- | --- | --- | --- | --- | --- |
| Cabrillospiral A | 0.209294 | U87MG | Cabrillospiral B | 0.175548 | A549 |
| Cabrillospiral A | 0.304065 | U87MG | Cabrillostatin | 1.55E-15 | A549 |
| Cabrillospiral A | 0.1364 | U87MG | Cabrillostatin | 1.55E-15 | A549 |
| Cabrillospiral B | 0.141661 | U87MG | Cabrillostatin | 1.55E-15 | A549 |
| Cabrillospiral B | 0.045934 | U87MG | Cabrillospiral A | 0.434165 | OVCAR4 |
| Cabrillospiral B | 0.004458 | U87MG | Cabrillospiral A | 0.671568 | OVCAR4 |
| Cabrillostatin | 1.00E-16 | U87MG | Cabrillospiral A | 0.026876 | OVCAR4 |
| Cabrillostatin | 1.00E-16 | U87MG | Cabrillospiral B | 0.721262 | OVCAR4 |
| Cabrillostatin | 1.00E-16 | U87MG | Cabrillospiral B | 0.401609 | OVCAR4 |
| Cabrillospiral A | 0.190758 | AsPC1 | Cabrillospiral B | 0.5173 | OVCAR4 |
| Cabrillospiral A | 0.362119 | AsPC1 | Cabrillostatin | 3.28E-05 | OVCAR4 |
| Cabrillospiral A | 0.256499 | AsPC1 | Cabrillostatin | 1.28E-06 | OVCAR4 |
| Cabrillospiral B | 0.39188 | AsPC1 | Cabrillostatin | 0.000847 | OVCAR4 |
| Cabrillospiral B | 0.263472 | AsPC1 | Cabrillospiral A | 0.332002 | HEPG2 |
| Cabrillostatin | 4.77E-15 | AsPC1 | Cabrillospiral A | 0.376773 | HEPG2 |
| Cabrillostatin | 4.77E-15 | AsPC1 | Cabrillospiral A | 0.711658 | HEPG2 |
| Cabrillostatin | 4.77E-15 | AsPC1 | Cabrillospiral B | 0.194758 | HEPG2 |
| Cabrillospiral A | 0.45305 | WM164 | Cabrillospiral B | 0.399985 | HEPG2 |
| Cabrillospiral A | 0.00022 | WM164 | Cabrillospiral B | 0.64405 | HEPG2 |
| Cabrillospiral A | 2.45E-05 | WM164 | Cabrillostatin | 1.06E-08 | HEPG2 |
| Cabrillospiral B | 5.57E-05 | WM164 | Cabrillostatin | 0.039649 | HEPG2 |
| Cabrillospiral B | 0.008047 | WM164 | Cabrillostatin | 0.337498 | HEPG2 |
| Cabrillospiral B | 2.47E-05 | WM164 | Cabrillospiral A | 0.142646 | 786-O |
| Cabrillostatin | 1.55E-15 | WM164 | Cabrillospiral A | 0.191146 | 786-O |
| Cabrillostatin | 1.55E-15 | WM164 | Cabrillospiral A | 0.528463 | 786-O |
| Cabrillostatin | 1.55E-15 | WM164 | Cabrillospiral B | 0.014953 | 786-O |
| Cabrillospiral A | 0.236295 | DU145 | Cabrillospiral B | 0.285538 | 786-O |
| Cabrillospiral A | 0.216205 | DU145 | Cabrillospiral B | 0.260704 | 786-O |
| Cabrillospiral A | 0.331292 | DU145 | Cabrillostatin | 0.320406 | 786-O |
| Cabrillospiral B | 0.422186 | DU145 | Cabrillostatin | 0.007764 | 786-O |
| Cabrillospiral B | 0.02437 | DU145 | Cabrillostatin | 0.078936 | 786-O |
| Cabrillospiral B | 0.594261 | DU145 | Cabrillospiral A | 0.103913 | HCT116 |
| Cabrillostatin | 1.00E-16 | DU145 | Cabrillospiral A | 0.452732 | HCT116 |
| Cabrillostatin | 6.68E-08 | DU145 | Cabrillospiral A | 0.066401 | HCT116 |
| Cabrillostatin | 4.41E-08 | DU145 | Cabrillospiral B | 0.228838 | HCT116 |
| Cabrillospiral A | 0.059789 | A549 | Cabrillospiral B | 0.210873 | HCT116 |
| Cabrillospiral A | 0.093785 | A549 | Cabrillospiral B | 0.789221 | HCT116 |
| Cabrillospiral A | 0.103537 | A549 | Cabrillostatin | 3.35E-07 | HCT116 |
| Cabrillospiral B | 0.550484 | A549 | Cabrillostatin | 3.77E-12 | HCT116 |
| Cabrillospiral B | 0.276498 | A549 | Cabrillostatin | 5.08E-12 | HCT116 |

Table S9. Bioactivity profile for cabrillostatin (1) relative to reference compounds.

| Cell line | Compound name | Predicted compound category | Confidence score |
| --- | --- | --- | --- |
| U87MG | Cabrillostatin | HDAC | 0.097434563 |
| U87MG | Cabrillostatin | HDAC | 0.051281913 |
| U87MG | Cabrillostatin | HDAC | 0.065527331 |
| AsPC1 | Cabrillostatin | DMSO | 0.057007311 |
| AsPC1 | Cabrillostatin | DMSO | 0.068606556 |
| AsPC1 | Cabrillostatin | HDAC | 0.099184119 |
| WM164 | Cabrillostatin | HDAC | 0.003963972 |
| WM164 | Cabrillostatin | HDAC | 0.029267674 |
| WM164 | Cabrillostatin | HDAC | 0.016086107 |
| DU145 | Cabrillostatin | DMSO | 0.004070139 |
| DU145 | Cabrillostatin | DMSO | 0.002048989 |
| DU145 | Cabrillostatin | DMSO | 0.003589487 |
| A549 | Cabrillostatin | HDAC | 0.063783546 |
| A549 | Cabrillostatin | HDAC | 0.060002645 |
| A549 | Cabrillostatin | HDAC | 0.053045952 |
| HCT116 | Cabrillostatin | DMSO | 0.021928432 |
| HCT116 | Cabrillostatin | HDAC | 0.030559655 |
| HCT116 | Cabrillostatin | HDAC | 0.025467136 |

Table S10. Heart relevant drugs used as reference compounds in the IPSC-CM model.

| Compound | Category | Targets | Pathway | Source well conc. µM |
| --- | --- | --- | --- | --- |
| Metoprolol Tartrate | Neuronal Signaling | Adrenergic Receptor | Neuronal Signaling | 10000 |
| Dronedarone HCl | Transmembrane Transporters | Potassium Channel, Sodium Channel, Calcium Channel | Transmembrane Transporters | 10000 |
| Carvedilol | Neuronal Signaling | Adrenergic Receptor | Neuronal Signaling | 10000 |
| Amiodarone HCl | Transmembrane Transporters | Autophagy, Potassium Channel | Transmembrane Transporters | 10000 |
| Ibutilide Fumarate | Transmembrane Transporters | Sodium Channel | Transmembrane Transporters | 10000 |
| Diltiazem HCl | Transmembrane Transporters | Calcium Channel | Transmembrane Transporters | 10000 |
| Ajmaline | Cardiovascular Disease | Potassium Channel Sodium Channel | Cardiovascular Disease | 10000 |
| Atenolol | Neuronal Signaling | Adrenergic Receptor | Neuronal Signaling | 10000 |
| Adenosine 5'-monophosphate monohydrate | PI3K/Akt/mTOR | AMPK | PI3K/Akt/mTOR | 10000 |
| Sparteine | Others | Others | Others | 10000 |
| Adenosine disodium triphosphate | Cancer | Others | Cancer | 10000 |
| Entinostat (MS-275) | Epigenetics | HDAC | Epigenetics | 10000 |
| Nebivolol HCl | Neuronal Signaling | Adrenergic Receptor | Neuronal Signaling | 10000 |
| Bisoprolol fumarate | Neuronal Signaling | Adrenergic Receptor | Neuronal Signaling | 10000 |
| Lidocaine | Neuronal Signaling | Histamine Receptor | Neuronal Signaling | 10000 |
| Adenosine | GPCR & G Protein | Adenosine Receptor | GPCR & G Protein | 10000 |
| Dofetilide | Transmembrane Transporters | Potassium Channel | Transmembrane Transporters | 10000 |
| Lidocaine hydrochloride | Angiogenesis | EGFR | Angiogenesis | 10000 |
| Mexiletine HCl | Transmembrane Transporters | Sodium Channel | Transmembrane Transporters | 10000 |
| Digoxin | Transmembrane Transporters | Sodium Channel | Transmembrane Transporters | 10000 |
| Labetalol HCl | GPCR & G Protein | Adrenergic Receptor | GPCR & G Protein | 10000 |
| (R)-(+)-Atenolol HCl | Others | Adrenergic Receptor | Others | 10000 |
| Disopyramide Phosphate | Others | Others | Others | 10000 |
| Procainamide HCl | Transmembrane Transporters | DNA Methyltransferase, Sodium Channel | Transmembrane Transporters | 10000 |
| Verapamil HCl | Transmembrane Transporters | Calcium Channel | Transmembrane Transporters | 10000 |
| Propranolol HCl | GPCR & G Protein | Adrenergic Receptor | GPCR & G Protein | 10000 |
| Esmolol HCl | Neuronal Signaling | Adrenergic Receptor | Neuronal Signaling | 10000 |
| Timolol Maleate | Neuronal Signaling | Adrenergic Receptor | Neuronal Signaling | 10000 |
| Quinidine sulfate | Transmembrane Transporters | Sodium Channel | Transmembrane Transporters | 10000 |
| Propafenone HCl | Transmembrane Transporters | Sodium Channel | Transmembrane Transporters | 10000 |
| Sotalol HCl | Neuronal Signaling | Adrenergic Receptor | Neuronal Signaling | 10000 |
| Phenytoin Sodium | Transmembrane Transporters | Sodium Channel | Transmembrane Transporters | 10000 |
| Phenytoin | Transmembrane Transporters | Sodium Channel | Transmembrane Transporters | 10000 |
| Dronedarone | Others | Others | Others | 10000 |
| Carvedilol Phosphate | Others | Others | Others | 10000 |
| Esmolol | Neuronal Signaling | Adrenergic Receptor | Neuronal Signaling | 10000 |
| Nebivolol | Neuronal Signaling | Adrenergic Receptor | Neuronal Signaling | 10000 |
| (-)-Sparteine Sulfate | Transmembrane Transporters | Sodium Channel | Transmembrane Transporters | 10000 |
| Disopyramide | Transmembrane Transporters | Sodium Channel | Transmembrane Transporters | 10000 |
| Metoprolol | Neuronal Signaling | Adrenergic Receptor | Neuronal Signaling | 10000 |

Table S11. Compounds detected in MASST datasets and corresponding MassIVE identifiers.

| Compound name or *m/z* | MassIVE dataset identifiers |
| --- | --- |
| Cabrillostatin (**1**) | MSV000086926, MSV000087650, MSV000080176, MSV000088823, MSV000087735, MSV000087322, MSV000087588, MSV000084755, MSV000085852, MSV000085779 |
| Aplysiopsene A | MSV000085159, MSV000078811 |
| 757.447 [M+H]^+^ | MSV000086926, MSV000084755, MSV000084741, MSV000084158, MSV000087006, MSV000082082, MSV000080015, MSV000080507, MSV000085480, MSV000088164, MSV000083632, MSV000082312, MSV000087650, MSV000085025, MSV000082952, MSV000088021, MSV000084744, MSV000083889, MSV000084119, MSV000086236, MSV000081731, MSV000085779 |
| 727.437 [M+H]^+^ | MSV000084741, MSV000084755 |
| 743.431 [M+H]^+^ | MSV000086926 |
| 769.448, [M+H]^+^ | MSV000086926, MSV000084755, MSV000087006, MSV000082312, MSV000084741 |
| 1031.550, [M+H]^+^ | MSV000084741 |
| 577.381, [M+H]^+^ | MSV000085480, MSV000084741, MSV000082312, MSV000085025, MSV000085786, MSV000087005 |
| 1086.6190, [M+NH_4_]^+^ | MSV000086926, MSV000087006, MSV000085786, MSV000087449, MSV000083889, MSV000083888 |
| 456.1317 [M+Na]^+^ | MSV000082082, MSV000080015, MSV000085786 |
| 644.3936 [M+H]^+^ | MSV000084741, MSV000082084, MSV000082083, MSV000083889, MSV000085779 |

Table S12. Cell painting dyes used for the phenotypic screening.

| **Material** | **Stock concentration** | **Final concentration [µl/ml]** | **Excitation wavelength [nm]** |
| --- | --- | --- | --- |
| Phalloidin | 2 units | 0.5 | 568 |
| Concanavalin A | 5 mg/mL | 20 | 488 |
| SYTO 14 green | 5 mM | 0.6 | 490-515 |
| Hoechst | 10 mg/mL | 0.5 | 385 |
| Wheat-germ agglutinin | 1mg/mL | 1.5 | 555 |
| Mitotracker deep red | 5 µM | 0.5 µM | 647 |
